# Relaxation kinetics controls cell transport in crowded active monolayers

**DOI:** 10.64898/2026.11.03.763953

**Authors:** Steven A. Silber, Mikko Karttunen

## Abstract

Soft cancer cells embedded in stiffer epithelial layers migrate in bursts, which prior simulations have attributed to their deformability. In a dense layer, however, interfacial tension sets two things at once: the energy cost of a deformation and the rate at which it relaxes. Here we disentangle the two in an active phase-field model of a near-confluent monolayer in which relaxation rate and free energy can be varied independently. Crowding slows cells not by pushing them back bodily but through the relaxation of their own deformations against neighbours, which cancels most of their propulsion. Slowing this relaxation in a minority of cells, at unchanged energetics and propulsion, more than doubles their transport and speeds up the untouched majority by half. Conversely, the advantage of soft cells vanishes, and at high soft-cell fractions reverses, once their relaxation is made as fast as that of normal cells. Independent of composition, mixtures transport like a uniform layer with the geometric-mean interfacial tension. Passive relaxation kinetics thus emerges as an independent lever on transport, suggesting that the fluidity of heterogeneous tissues may depend on how quickly cells recover their shape, not only on how stiff they are.

---

In a confluent monolayer, cells cover the substrate with essentially no gaps. As a consequence, a cell cannot move far through the layer without changing shape and rearranging its contacts with neighboring cells. The elementary contact rearrangement, often called a T1 transition, occurs when one cell-cell contact shrinks and disappears while a new contact forms between a different pair of cells [1–3].

In multiphase-field (MPF) monolayers, T1 neighbor exchanges involve localized cell deformation and slow post-exchange energy relaxation, and can trigger nearby rearrangements [4]. Such rearrangements contribute to morphogenesis and epithelial repair [5, 6]. Their suppression is associated with rigidity and jamming in models of confluent tissues [7–9], with glass-like dynamics in epithelial monolayers [10], and cell shape serves as an experimental signature of jamming [11].

Real tissues are mechanically heterogeneous. For example, cancer cells are often more deformable than benign cells [12, 13], and tumors contain cells with widely different deformabilities and regions with distinct migratory states [14, 15]. This raises two questions: does a local mechanical difference affect only the altered cell or does it also reorganize the surrounding tissue, and does it act through the energetic cost of deformation or through the rate at which deformations relax? In MPF monolayers, the response to a single driven probe depends on the collective state: the force needed to set the embedded probe in motion falls to zero as the tissue reaches its solid–liquid transition [16]. Deformability and motility jointly control this transition [16, 17], and deformability lowers the motility at which motility-induced phase separation sets in [18].

Lee *et al*. observed pulsating motion of soft MDA-MB-231 carcinoma cells embedded in a monolayer of stiffer MCF10A epithelial cells [19]. Because the two cell lines differ in nuclear mechanics, adhesion, and active motility, the experiment did not isolate deformability as the sole cause of the observed motion. The two lines also differ in dissipative response: AFM microrheology shows a higher loss tangent *G*^*′′*^*/G*^*′*^ in MDA-MB-231 [20]. Palmieri *et al*. used a cell-resolved phase-field model to isolate the effect of changing a single cell’s interfacial coefficient [21]. A later phase-field study of Yin *et al*. found velocity bursts of the soft cell that follow its shape changes [22]. Using a related sharp-interface formulation, in which the cell boundaries are treated as mathematically sharp, Bresler *et al*. found substantial realization-to-realization variation in the altered-cell motion [23]. The same sharp-interface model has also been used to map where active cells jam, including mixed soft–stiff populations [24].

In the MPF representation [21, 25–29], each cell is described by a smooth field that is approximately one in the cell interior and zero outside, with a finite-width transition across the cell boundary. This representation resolves evolving cell shapes and contacts without tracking each boundary as a curve [30, 31]. The coefficient *γ* multiplies the free-energy contribution associated with this boundary (see Methods). At fixed interface width, *γ* is proportional to the effective line tension, i.e., the free-energy cost per unit length of the cell boundary [32]. Lowering *γ* makes deformations that increase the cell perimeter energetically less costly. simulated motion to an elasticity mismatch between the tagged cell and the surrounding monolayer. Within the model, however, *γ* is an interfacial-energy coefficient, not a bulk elastic modulus or a measured Young’s modulus. Moreover, the fields relax by non-conserved (Model A / Allen–Cahn) dynamics [33, 34], in which the kinetic coefficient *M* sets how fast each field relaxes toward lower free energy. Because the cell boundary then relaxes at a rate proportional to *Mγ* (Sec. 1.2), lowering *γ* at fixed *M* also slows shape relaxation. Thus, varying *γ* alone changed both the energetic cost and the relaxation rate for deformations.

How tissues flow and fail is not set by the energetic cost of deformation alone. Internal dissipation enables sustained flows in vertex models [35] and is required for spontaneous hole formation in MPF monolayers [36], whereas intercellular friction solidifies them [37]. Intercellular viscosity is biologically regulated [38], and whether cell–cell contacts fail in a ductile or a brittle way depends on both cortical tension and how quickly cells recover their shape [39]. Weakening adhesion can even make an epithelial monolayer more fluid while junctional tension, traction, density and cell shape stay unchanged [40]. In the MPF model, internal dissipation of this kind is set by 1*/M* (Sec. 3.1), a resistance to shape relaxation that is distinct from substrate and intercellular friction.

We use replicated paired simulations of the Palmieri *et al*. model [21], in which altered and control monolayers share initial fields and imposed polarity histories, to separate the energetic and kinetic consequences of a local mechanical change. The control with reduced *M* keeps the free-energy parameters fixed, compensated mixtures change *γ* while holding *Mγ* fixed, and homogeneous monolayers at the mean *γ* remove the contrast between cells.

We ask four questions: 1) Which contributions to centroid motion oppose propulsion in a crowded layer, and how do they depend on the kinetic coefficient *M*, which multiplies every passive relaxation term in the field equation? 2) When a cell is softened, how much of its transport change is energetic (line tension) and how much kinetic (relaxation rate)? 3) In mixtures, how much of the transport change follows the mean *γ* and how much the contrast between cells, that is, the difference between the *γ* values of the two cell types? 4) How much does softening raise a single cell’s own diffusivity, and does one soft cell change the motion of the rest of the monolayer? We find that crowding opposes propulsion mainly through field relaxation, that slowing relaxation alone raises the transport of both the modified and the unmodified cells, and that equalizing *Mγ* removes the higher diffusivity of soft cells. At a common *M*, population transport follows the geometric-mean *γ* with no resolved residual effect of contrast, and a single soft cell raises its own diffusivity by about 14% without a resolved effect on the rest of the monolayer.

## 1 Results

Cell *i* is represented by a smooth field *ϕ*_*i*_(***r***,*t*) with *ϕ*_*i*_ *≃*1 inside and *ϕ*_*i*_ *≃* 0 outside. The free energy penalizes cell boundary, deviations from target area and overlap between cells.

Normal and soft cells have interfacial coefficients *γ*_n_ =1 and *γ*_s_ = 0.35, respectively, and *f*_s_ denotes the soft-cell fraction. In the binary mixtures, both populations have relaxation coefficient of *M*_*i*_ =0.5. Figure 1 shows example configurations with one soft cell (Movie 1) and with finite soft-cell fractions (Movie 2). At the nominal packing fraction of 0.90, the monolayers are near-confluent with small gaps at multicellular junctions.

**Figure 1.**
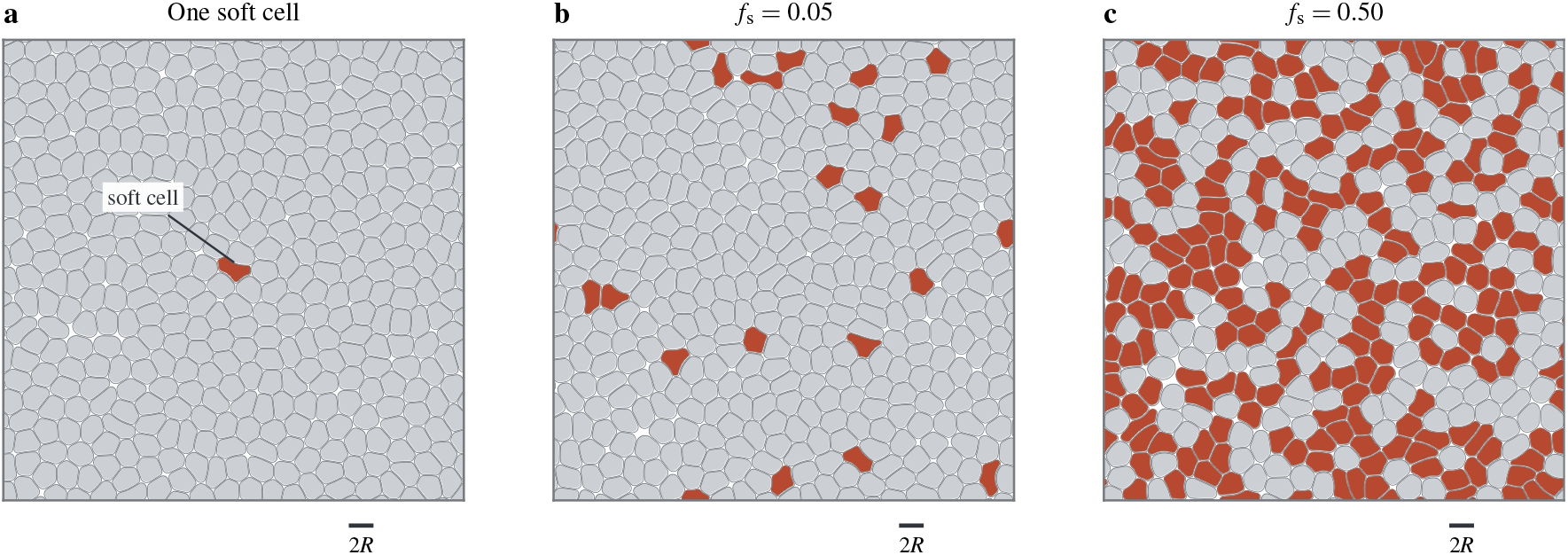
Simulated monolayers with interfacial heterogeneity. Example configurations after 50*τ* of active motion with (a) one soft cell, and (b) 5% and (c) 50% soft cells. All panels have 400 cells at nominal packing fraction 0.90. Gray and orange indicate normal (*γ*_n_ = 1) and soft (*γ*_s_ = 0.35) cells, respectively. Outlines show the *ϕ*_*i*_ = 0.5 contours.

We quantify transport by the 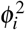-weighted centroid ***r***_*i*_(*t*) using finite-lag diffusivity

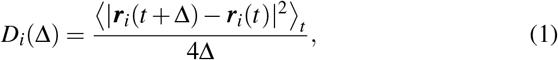

where the average includes all origins whose complete displacement interval lies within the active observation window. The polarity-persistence time *τ* is the mean waiting time between independent random reorientations of a cell’s imposed propulsion direction. We set *τ* = 10^4^ in model time units, following Palmieri *et al*. [21]; the reorientation process is detailed in Sec. 3.1. Our primary transport measure is *D*_8_,the finite-lag diffusivity at Δ = 8*τ*, with *D*_8,*i*_ = *D*_*i*_(8*τ*) for cell *i*.The choice of lag is motivated in Methods (Figs. S1 and S2). The symbols *D*_s_, *D*_n_ and *D*_pop_ denote averages over soft cells, normal cells and the full population.

### 1.1 Crowding opposes propulsion through field relaxation, not rigid advection

A free cell with propulsion speed *v*_*A*_ has 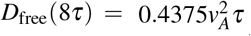 (Sec. S1). In all-normal monolayers, *D*_pop_ *≈*0.095*D*_free_(8*τ*). We first identify which contributions to centroid motion account for this reduction. The centroid displacement over a lag Δ is the sum of displacements due to imposed propulsion (***d***^*A*^), interaction advection (***d***^*I*^), and field relaxation (***d***^*R*^) (Methods; Secs. S2.1 and S2.2). The physical transport balance can be given in terms of their second moments as

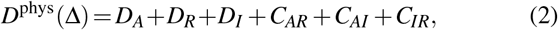

where *D*_*X*_ = ⟨|***d***^*X*^| ^2^ ⟩/(4Δ) and *C*_*XY*_ = ⟨ ***d***^*X*^ *·****d***^*Y*^⟩ /2Δ. Here *X,Y* ∈ {*A, I, R*} label propulsion, interaction advection and field relaxation, respectively. *D*_*X*_ is the contribution to diffusivity from *X*, and *C*_*XY*_ accounts for correlations between the respective displacement contributions.

In all-normal monolayers the propulsion contribution equals the free-cell value, 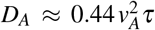 (Sec. S1), whereas 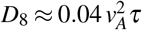. The difference is carried by field relaxation: 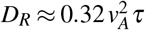 and 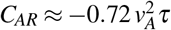, while the three interaction-advection terms sum to 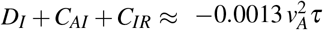 (Sec. S2.3, Table S1). Following a polarity reorientation, the relaxation velocity projected along the new direction averages ≈ −0.51 *v*_*A*_ over the first 0–0.025*τ* after a reorientation and reaches −0.81 *v*_*A*_ at 1–2*τ* (Sec. S2.4; Figs. S3 and S4). Relaxation cancels up to 80% of the propulsion velocity, and direct pushing by neighbors contributes a few per cent.

### 1.2 The kinetic coefficient controls transport at fixed free-energy parameters

To test whether slower passive relaxation increases transport, we reduce *M*_*i*_ from 0.5 to 0.175 in 30% of the cells (modified cells) while keeping all free-energy parameters fixed, with *γ*_*i*_ = 1 for every cell. The remaining cells retain *M*_*i*_ = 0.5 (unmodified cells). This changes interfacial, area and overlap relaxation together (Table 1; Sec. S3.1, Table S2). The reduction factor, 0.35, is the one applied to *γ* in the soft cells of the binary mixtures, i.e., modified cells and binary soft cells have the same interfacial relaxation rate *M*_*i*_*γ*_*i*_ = 0.175. Relative to the corresponding cells in paired all-normal references with *M*_*i*_ = 0.5, the geometric means of the transport ratios are 2.246 [2.193, 2.299] for modified and 1.498 [1.450, 1.547] for unmodified cells, and 1.723 [1.681, 1.766] for the population (Fig. 2a). Reducing *M*_*i*_ therefore increases transport at unchanged interfacial energy. The enhancement persists at four times higher advection mobility (*ξ* = 375): ten paired seeds give transport ratios 2.105 [2.050, 2.162] for modified, and 1.457 [1.407, 1.509] for unmodified cells (Fig. 2a; Sec. S3.2; Fig. S5). Relaxation still accounts for most of the gain (Table S3), although interaction advection partly offsets it in modified cells. Movie 3 shows this control and its all-normal reference at *ξ* = 375.

**Table 1.**
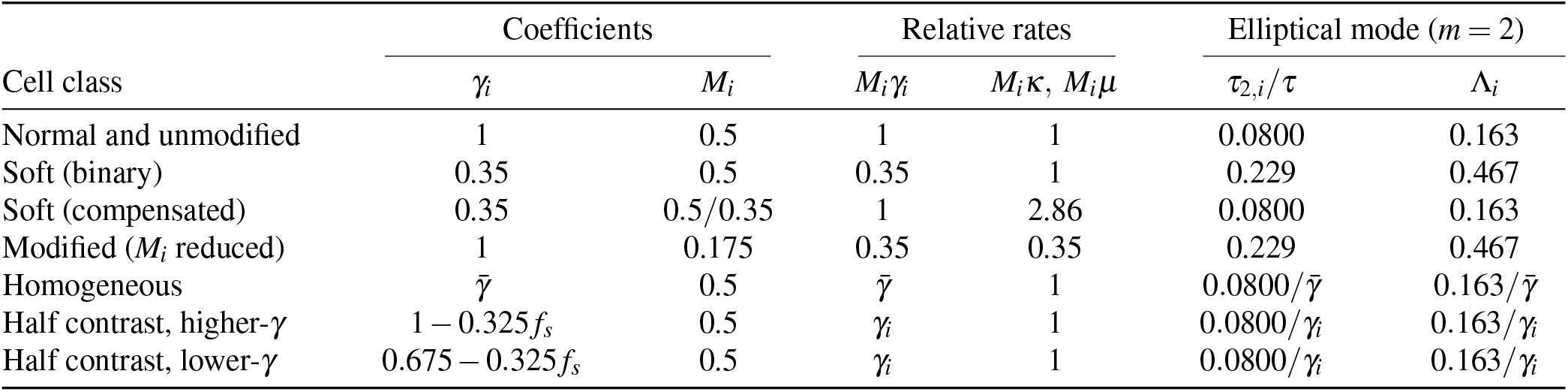
Passive relaxation coefficients. Binary mixtures contain normal and soft cells with different *γ*_*i*_ but the same *M*_*i*_. Compensated mixtures retain these *γ*_*i*_ values and increase *M*_*i*_ in soft cells so that *M*_*i*_*γ*_*i*_ is equal in both populations. In the control with reduced *M*_*i*_, every cell has *γ*_*i*_ = 1. Modified cells have reduced *M*_*i*_, while unmodified cells retain the normal-cell coefficients. The single-soft-cell and low-fraction systems use the binary-mixture coefficients. The homogeneous row describes the arithmetic-mean reference, 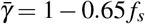, and the half-contrast rows follow Eq. S39. The columns for *γ*_*i*_ and *M*_*i*_ give their absolute values. The interfacial, overlap and area relaxation coefficients, *M*_*i*_*γ*_*i*_, *M*_*i*_*κ* and *M*_*i*_*µ*, are divided by their respective normal-cell values. All cells have *κ* = 10 and *µ* = 1. The advection coefficient *κ/ξ* is identical for every cell within a simulation, with *ξ* = 1500 except in the robustness test at *ξ* = 375 (Sec. S3.2). The elliptical relaxation time *τ*_2,*i*_ = *R*^2^*/*(6*M*_*i*_*γ*_*i*_) and the corresponding propulsion distance Λ_*i*_ = *v*_*A*_*τ*_2,*i*_*/R* are evaluated using *R* = 49, *v*_*A*_ = 0.01 and *τ* = 10^4^ (Eq. 4).

| Cell class | Coefficients |  | Relative rates |  | Elliptical mode (*m* = 2) |  |
|---|---|---|---|---|---|---|
|  | *γ*<sub>*i*</sub> | *M*<sub>*i*</sub> | *M*<sub>*i*</sub>*γ*<sub>*i*</sub> | *M*<sub>*i*</sub>*κ*, *M*<sub>*i*</sub>*μ* | *τ*<sub>2,*i*</sub>/*τ* | Λ<sub>*i*</sub> |
| Normal and unmodified | 1 | 0.5 | 1 | 1 | 0.0800 | 0.163 |
| Soft (binary) | 0.35 | 0.5 | 0.35 | 1 | 0.229 | 0.467 |
| Soft (compensated) | 0.35 | 0.5/0.35 | 1 | 2.86 | 0.0800 | 0.163 |
| Modified (*M*<sub>*i*</sub> reduced) | 1 | 0.175 | 0.35 | 0.35 | 0.229 | 0.467 |
| Homogeneous | *γ̄* | 0.5 | *γ̄* | 1 | 0.0800/*γ̄* | 0.163/*γ̄* |
| Half contrast, higher-*γ* | 1 − 0.325*f*<sub>*s*</sub> | 0.5 | *γ*<sub>*i*</sub> | 1 | 0.0800/*γ*<sub>*i*</sub> | 0.163/*γ*<sub>*i*</sub> |
| Half contrast, lower-*γ* | 0.675 − 0.325*f*<sub>*s*</sub> | 0.5 | *γ*<sub>*i*</sub> | 1 | 0.0800/*γ*<sub>*i*</sub> | 0.163/*γ*<sub>*i*</sub> |

**Figure 2.**
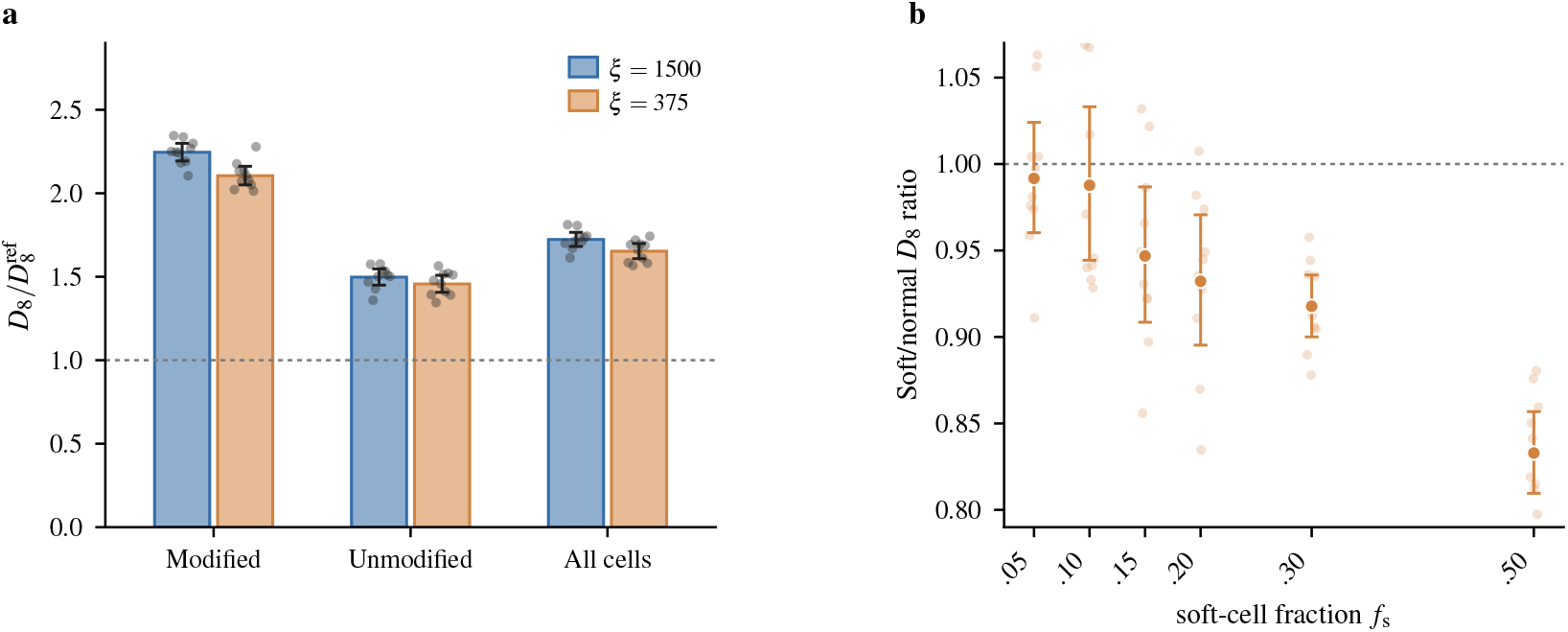
Transport under changes in *M*_*i*_ and *γ*_*i*_. (a) Reducing *M*_*i*_ in 30% of cells while keeping *γ*_*i*_ = 1 for every cell: transport of modified and unmodified cells, and the full population divided by 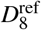 for the corresponding groups in paired all-normal monolayers with *M*_*i*_ = 0.5 at the same *ξ*. The two series compare advection mobilities 1*/ξ* at *ξ* = 1500 and 375. (b)Soft/normal *D*_8_ ratio within compensated mixtures, where *M*_*i*_*γ*_*i*_ is equal in the two populations. Faded symbols show individual simulations. Bars in (a) and filled symbols in (b) show geometric means, with error bars giving pointwise 95% confidence intervals over ten paired seeds in each case.

In the binary mixtures described above, soft cells have higher *D*_8_ than normal cells (Fig. S6a). To test whether this difference persists when the interfacial relaxation coefficients *M*_*i*_*γ*_*i*_ are equal, we use compensated mixtures that retain *γ*_n_ = 1 and *γ*_s_ = 0.35 but assign different *M*_*i*_ to the two cell types. Normal cells retain *M*_*i*_ = 0.5, while soft cells are assigned *M*_*i*_ = 0.5*/*0.35, giving *M*_*i*_*γ*_*i*_ = 0.5 for both cell types. In these compensated mixtures, no soft–normal transport difference was resolved at *f*_s_ = 0.05 or 0.10. From *f*_s_ = 0.15 onward, soft cells have smaller *D*_8_ than normal cells with *D*_s_*/D*_n_ = 0.833 [0.810, 0.857] at *f*_s_ = 0.50 (Fig. 2b). Compensated soft cells nevertheless move faster instantaneously: their mean-square centroid speed exceeds that of normal cells by 18% at 5% and by 1% at 50%, while their velocity-correlation integral is 16–19% smaller at every fraction (Fig. S6b). Higher instantaneous speed can therefore coexist with lower transport. In the noise-free dynamics of Eqs. (14) and (15) (Methods), the free-energy coefficients enter through the rate products *M*_*i*_*γ*_*i*_, *M*_*i*_*κ* and *M*_*i*_*µ*, and through the advection coefficient *κ/ξ*. The interfacial constant *γ*_*i*_ of cell *i* enters no other cell’s equation and its contribution to the interaction force vanishes by translational invariance. Here *κ* and *µ* set the overlap and area penalties, respectively, and 1*/ξ* controls interaction advection. The passive fixed points depend on the ratios *γ*_*i*_ : *κ* : *µ*. Thus, lowering *γ*_*i*_ at fixed *M*_*i*_ is dynamically identical to slowing only interfacial relaxation in the soft cells, whereas reducing *M*_*i*_ slows interfacial, overlap and area relaxation together in the modified cells.

For a homogeneous monolayer with kinetic coefficient *sM*, where *s >* 0 is a dimensionless scaling factor, rescaling time by *t*^*′*^ = *st* gives a system with kinetic coefficient *M* the same free energy, and transformed parameters

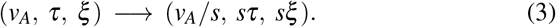

The two systems have the same persistence length *l*_*p*_ = *v*_*A*_*τ* and normalized diffusivity *D*(Δ)*/*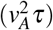 at matching Δ*/τ*. For *s <* 1, uniformly slower relaxation is thus equivalent to faster propulsion at fixed persistence length (*ξ* is reduced by the same factor), a possible route to fluidization [17].

The field equation weights interfacial and overlap relaxation by coefficients proportional to *M*_*i*_*γ*_*i*_ and *M*_*i*_*κ* (Eqs. 14 and 15). The corresponding interfacial scales are

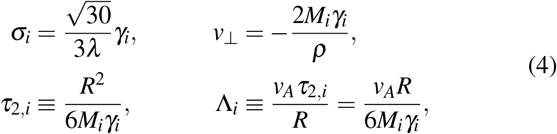

where *σ*_*i*_ is the planar line tension and *v*_⊥_is the Allen–Cahn normal velocity of an interface with local radius of curvature *ρ* [34, 41]. This coincides with the curvature term of the sharp-interface model of Ref. [23] at *M* = 1*/*2. A radial distortion *δ R* cos(*mθ*) of a cell of radius *R* therefore decays on *R*^2^*/*[2*M*_*i*_*γ*_*i*_(*m*^2^*−* 1)]; *τ*_2,*i*_ is the *m* = 2 (elliptical) value. The profile itself relaxes on *λ* ^2^*/*(45*M*_*i*_*γ*_*i*_), more than two orders of magnitude faster (Methods) meaning that cell-scale shape relaxation is the slow process. With the parameters used here, *τ*_2_ = 0.08*τ* for normal and unmodified cells and 0.23*τ* for soft and modified cells, comparable to the 0.1–0.5*τ* over which the opposing relaxation velocity builds up after a reorientation (Figs. S3 and S4). Λ_*i*_ is the fraction of a radius through which a cell is propelled before an elliptical distortion relaxes: 0.16 for normal and unmodified cells and 0.47 for soft and modified cells. At fixed *M*_*i*_, lowering *γ*_*i*_ lowers the line tension and raises Λ_*i*_ by the same factor. The compensated control keeps *M*_*i*_*γ*_*i*_, and hence *τ*_2,*i*_ and Λ_*i*_, fixed, whereas reducing *M*_*i*_ changes the overlap and area relaxation rates as well.

### 1.3 Reduced propulsion–relaxation cancellation in binary mixtures

Reducing *M*_*i*_ increases transport at fixed free-energy parameters. We next ask which contributions to cell displacement account for the transport increase in the binary mixtures of soft and normal cells. Relative to the paired all-normal reference, *D*_*A*_ (Eq. 2) is unchanged, while the magnitude of the negative cross term *C*_*AR*_ decreases (Fig. 3a). The increase in *C*_*AR*_ exceeds the decrease in *D*_*R*_, accounting for the population transport gain.

**Figure 3.**
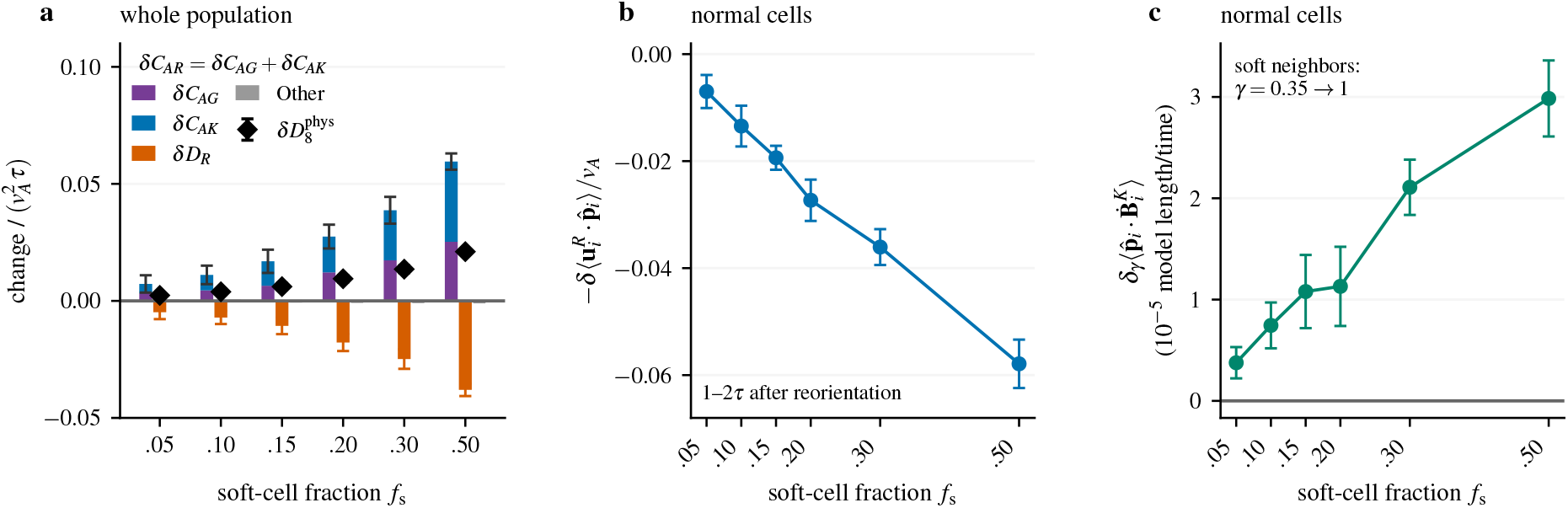
Reduced propulsion–relaxation cancellation in binary mixtures. (a) Changes in the population displacement balance at 8*τ. δ* denotes the difference between binary and all-normal. Stacked contributions give *δC*_*AR*_ with the paired interval for their sum. Signed contributions sum to 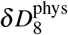(diamonds). Other terms are *δ* (*D*_*I*_ +*C*_*AI*_ +*C*_*IR*_) and *δ D*_*A*_ = 0. (b) Change in normal-cell opposing relaxation velocity 1–2*τ* after reorientation, conditional on no subsequent orientation event. Negative values indicate weaker opposition. (c) Difference in the initial rate of change of normal cells’ overlap imbalance 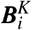 projected along propulsion after increasing the soft-cell coefficient from *γ*_s_ = 0.35 to 1. Both evaluations use the same fields and polarities, saved after 50*τ* of activity. Here, *δ*_*γ*_ denotes the difference between the modified evaluation and the original one. Positive values indicate faster initial development of overlap-induced centroid motion opposite propulsion. Means and pointwise 95% Student intervals use 10 paired seeds at each fraction. Displacement windows lie within 8–50*τ*. Reused seeds correlate estimates across fractions.

To relate this change in *C*_*AR*_ (Eq. 2) to the interfacial and overlap terms in the free energy (Eq. 14 in Methods), we write their contributions to the relaxation velocity as

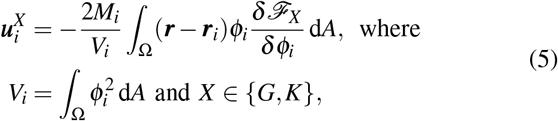

and *ℱ* _*G*_ and *ℱ* _*K*_ are the interfacial and overlap terms in Eq. 14. The area penalty has zero direct contribution to centroid velocity, so 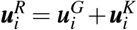 (Sec. S4) and their displacement correlations give *C*_*AR*_ = *C*_*AG*_ +*C*_*AK*_. Both cross-term contributions increase on average, and normal- and soft-cell means show the same balance. At 1–2*τ* after reorientation, the mean relaxation velocity opposing propulsion is smaller for normal cells in binary mixtures than in the all-normal reference with both interfacial and overlap contributions reduced (Fig. 3b). No systematic shortening of the time to reach 90% of this late response was resolved.

The overlap contribution to the relaxation velocity depends on where overlap occurs relative to the cell centroid. With 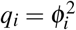 and *V*_*i*_ = ∫_Ω_ *q*_*i*_ d*A*,

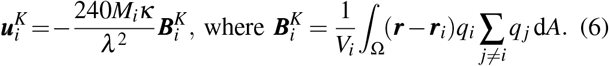

The vector 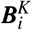 weights each region of overlap by its displacement from the cell centroid. Let 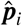 denote the unit vector along the imposed propulsion direction. A positive projection 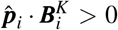, means that the distance-weighted overlap is greater ahead of the centroid than behind it, so Eq. 6 gives a relaxation component opposite propulsion. Its connection to transport follows by integrating over the same displacement window *w* =[*t*_0_,*t*_0_ +Δ]:

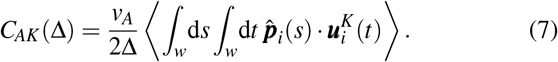

The average uses the same cells and windows as Eq. 2. Thus, *C*_*AK*_ depends on 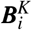 throughout the displacement window, including its correlations with propulsion at different times.

Using configurations saved after 50*τ* of activity, we evaluated the field equations with *γ*_s_ = 0.35 and again with *γ*_s_ =1. All cell fields, polarities and other parameters were kept unchanged. Increasing the soft-cell coefficient increased the mean initial rate of change of 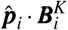 in normal cells (Fig. 3c; Sec. S4.1). Thus, Eq. 6 implies faster initial development of overlap-induced centroid motion opposite propulsion. The smaller values of 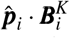 and weaker backward velocities were measured separately in the evolved mixtures. Together, these results support a mechanism in which weaker interfacial restoration changes the evolving overlap distribution, reducing its backward contribution during continued propulsion. Interfacial relaxation also produces less backward motion, and the two displacement correlations account for the measured reduction in cancellation.

The control with reduced *M*_*i*_ reveals a change in the temporal response of unmodified cells. Rescaling the propulsion–relaxation correlation of the corresponding cells in the all-normal reference to match the equal-time amplitude measured in the simulation with reduced *M*_*i*_ predicts a decrease in *C*_*AR*_ (Eq. 2), whereas the measured change is positive. Approximately 88% of this increase comes from correlations between propulsion and relaxation in subsequent polarity intervals. After reorientation, relaxation is less opposed to the preceding propulsion direction, even though its early opposition to the new direction strengthens. Thus, changing the passive kinetics of one population alters how the other population’s relaxation correlates with its propulsion history (Sec. S5; Figs. S3, S7 and S8).

Binary soft cells and modified cells in the control with reduced *M*_*i*_ have the same *M*_*i*_*γ*_*i*_ = 0.175, but different overlap and area relaxation coefficients, and different transport responses. In the control with reduced *M*_*i*_, most of the unmodified-cell *C*_*AR*_ increase involves propulsion before the most recent orientation event. In binary mixtures, correlations within the same interval of constant polarity account for most of the increase in the population average of *C*_*AR*_ relative to the 5% mixture. In compensated mixtures, *M*_*i*_*γ*_*i*_ = 0.5 for both cell types, and soft cells no longer have higher diffusivity than normal cells. Relative to all-normal controls, compensated soft cells instead gain transport through an increase in *D*_*R*_ despite a more negative *C*_*AR*_. Matching *Mγ* therefore does not fix either the displacement balance or its temporal correlations (Sec. S5.3; Figs. S9 and S10).

### 1.4 A single soft cell retains its higher diffusivity as system size increases

To establish whether the soft-cell transport advantage persists in large monolayers, we compared systems containing one soft cell with all-normal controls using *N* = 100–6400. Each pair shares its initial configuration and polarity history. The tagged-cell *D*_8_ ratio, pooled over *N* = 100–1600, is 1.137 [1.074, 1.204] (Fig. 4a). Regression of paired log ratios against log_2_ *N* resolved no systematic size dependence. The population-mean *D*_8_ shows no resolved increase relative to the all-normal controls with a pooled ratio of 1.0004 [0.9945, 1.0064] (Fig. 4b). The tagged-cell ratio remains above unity for lags from 0.5*τ* to 16*τ* (Fig. 4c).

**Figure 4.**
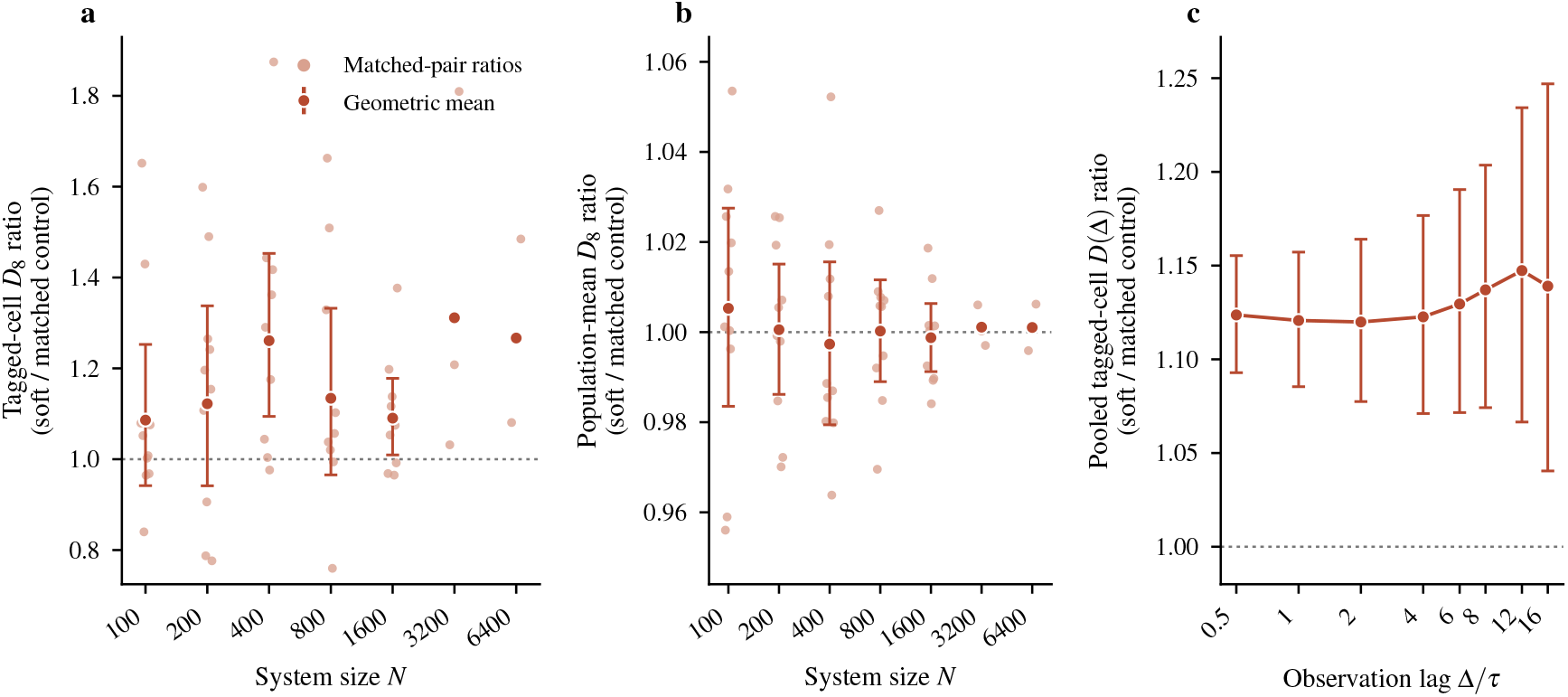
Finite-size transport response. (a) Tagged-cell transport (*D*_8_) in the soft branch relative to its paired all-normal control. (b) Corresponding population-averaged ratio. Faded symbols: individual replicates; filled symbols and error bars: geometric means and 95% confidence intervals with 10 replicates. The three *N* = 3200 and two *N* = 6400 replicates are shown individually. (c) Pooled tagged-cell ratio for *N* =100–1600 as a function of observation lag. Dotted lines: transport ratio of one.

Palmieri *et al*. reported an approximately 1.5-fold increase from one pair of 72-cell simulations [21]. They matched initial positions and reorientation sequences between their two 72-cell runs. The new elements here are replication over initial conditions and system size, and the mechanistic decomposition. We recover the same tendency with a smaller mean ratio and substantial variation among initial conditions. This is consistent with the run-to-run spread in the sharp-interface model, where the diffusion constant varied by *≈* 9% for the soft cell and *≈* 3% for normal cells across 20 runs [23]. The relative uncertainty of a time-averaged MSD grows with the ratio of lag to record length because overlapping displacement windows are correlated [42, 43].

### 1.5 Transport increases with soft-cell fraction

To test whether a larger soft-cell population also increases normal-cell transport, we varied the soft-cell fraction in *N* = 400 monolayers from *f*_s_ = 1*/*400 to 0.50 with finer sampling between 0.5% and 4% to resolve the response near the single-cell limit (Fig. 5a). Between 5% and 50%, mean *D*_8_ increased by approximately one third for both cell types. The soft/normal ratio ranged from 1.11-1.14 across the six fractions (Fig. S6a), close to the single cell ratio measured at 1.137. No difference between their slopes was resolved, and soft cells retained larger mean *D*_8_. The normal-cell increase persisted over all examined observation lags.

**Figure 5.**
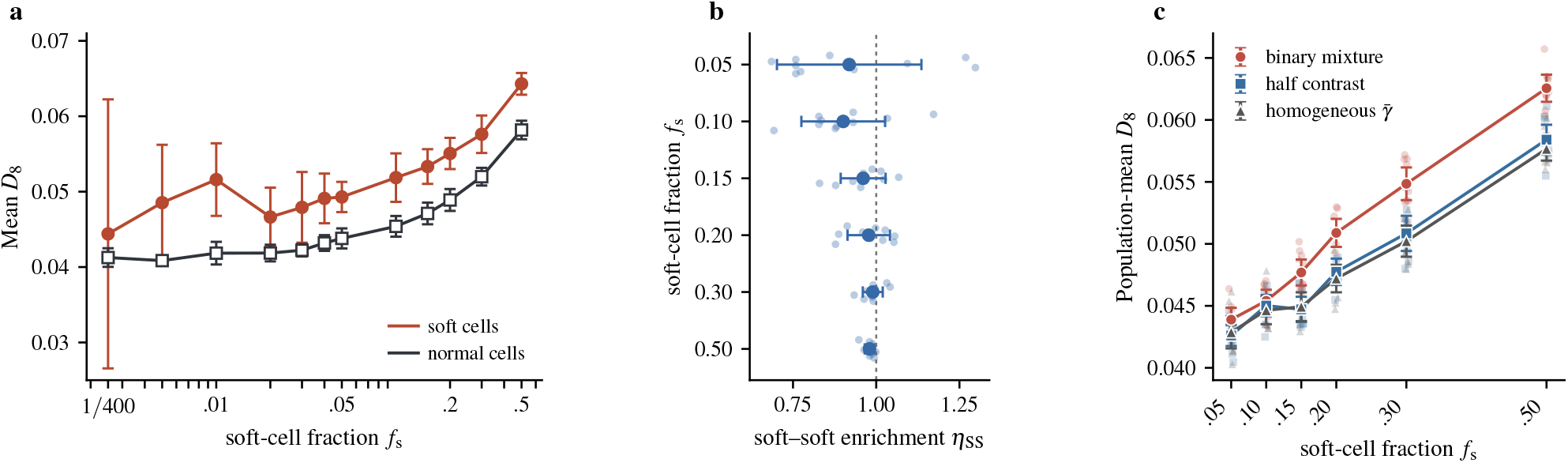
Composition-dependence of *D*_8_. (a) Mean soft- and normal-cell *D*_8_ in *N* = 400 monolayers. Symbols and error bars: means and 95% confidence intervals across independent initial conditions. (b) Soft–soft Delaunay-edge counts relative to their random-label expectations. Background symbols show individual simulations. Values near unity indicate random mixing. (c) Population-mean *D*_8_ in binary and half-contrast mixtures, and homogeneous monolayers at the same mean interfacial coefficient 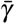, with matched polarity histories and common displacement windows within 0.01–50*τ*. Faded symbols: individual simulations; filled symbols and error bars: arithmetic means and pointwise 95% confidence intervals over ten paired seeds.

With one soft cell, no population transport increase was resolved. This is consistent with the finite-size results. Normal-cell transport rose across the 0.5–4% interval, reaching approximately 5% above the all-normal control at 4% (Fig. 5a). We identify geometric neighbors using Delaunay triangulation of the cell centroids. Two centroids are connected by an edge when the regions closest to them share a boundary. Across 5–50%, mean soft–soft Delaunay-edge counts did not exceed their random-label expectations, indicating no positive soft-cell sorting (Fig. 5b).

### 1.6 Population transport follows the geometric-mean interfacial coefficient

To assess how much of the transport increase with *f*_s_ can be explained by the changing mean interfacial coefficient 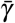, we compared each binary mixture with a homogeneous monolayer at

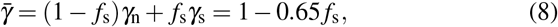

where *γ*_n_ = 1 and *γ*_s_ = 0.35. These homogeneous monolayers with 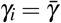 reproduce most of the composition-dependent increase in population transport, but with matched polarity histories and common 8*τ* displacement windows the full-contrast mixtures retain a higher mean transport. This was resolved at five of six fractions (Fig. 5c; Sec. S6; Fig. S11). We call the difference between the two interfacial coefficients, Δ*γ* = *γ*_n_*− γ*_s_, the contrast.

At each fraction, we compared three populations with the same 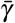: homogeneous monolayers with Δ*γ* = 0, half-contrast mixtures in which each coefficient deviates from 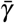 by half as much as in the binary mixture (Δ*γ* = 0.325; Eq. S39), and the binary mixtures of Sec. 1.5, which have full contrast (Δ*γ* = 0.65). We interpolated the measured population transport *D*_8_ of homogeneous monolayers as a function of *γ* to determine the uniform interfacial coefficient *γ*_eff_ that reproduces each mixture’s measured population transport, *D*_hom_(*γ*_eff_) = *D*_mix_ (Sec. S6). The effective coefficient *γ*_eff_ agrees with the geometric mean *γ*_*G*_ =exp ln *γ*_*i*_ within 1% at five of the six fractions and within 2.2% at *f*_*s*_ = 0.2, and lies below 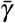 and above the harmonic mean 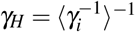(Fig. S12). The half-contrast mixtures agree with *γ*_*G*_ within their uncertainties.

At uniform *M*_*i*_, population transport therefore follows the geometric-mean interfacial coefficient, equivalently the mean logarithm of the interfacial relaxation rate *M*_*i*_*γ*_*i*_ with no resolved residual effect of contrast.

Contrast separates the two classes. At *f*_*s*_ = 0.20, 0.30 and 0.50, the lower-/higher-*γ* transport ratio in half-contrast mixtures exceeds that of the same labeled groups in the homogeneous reference by 4–6% (pointwise 95% intervals; Fig. S13). A class difference linear in Δ*γ* gives the first-order prediction that the half-contrast transport ratio equals the square root of the full-contrast ratio (1.11–1.14),i.e. 1.05–1.07, consistent with these values.

### 1.7 Normal-cell gain is spread across the layer

To test whether enhanced transport is concentrated near a soft cell, we measured normal-cell displacements as a function of their initial distance from the tagged cell in the finite-size series. The soft-branch near/far contrast was normalized by the corresponding contrast around the tagged normal cell in the paired control. The profile did not show any systematic radial enhancement (Fig. 6a). The pooled near/far contrast is 1.012 [0.971, 1.055] (Fig. 6b). Because each profile is normalized by its own far-field value, this test detects a response concentrated near the soft cell, not a spatially uniform one. The pooled result resolved no common near-cell enhancement across *N* = 400–1600, although the *N* = 800 estimate is 1.079 [1.012, 1.150] with a pointwise interval (Fig. 6b).

**Figure 6.**
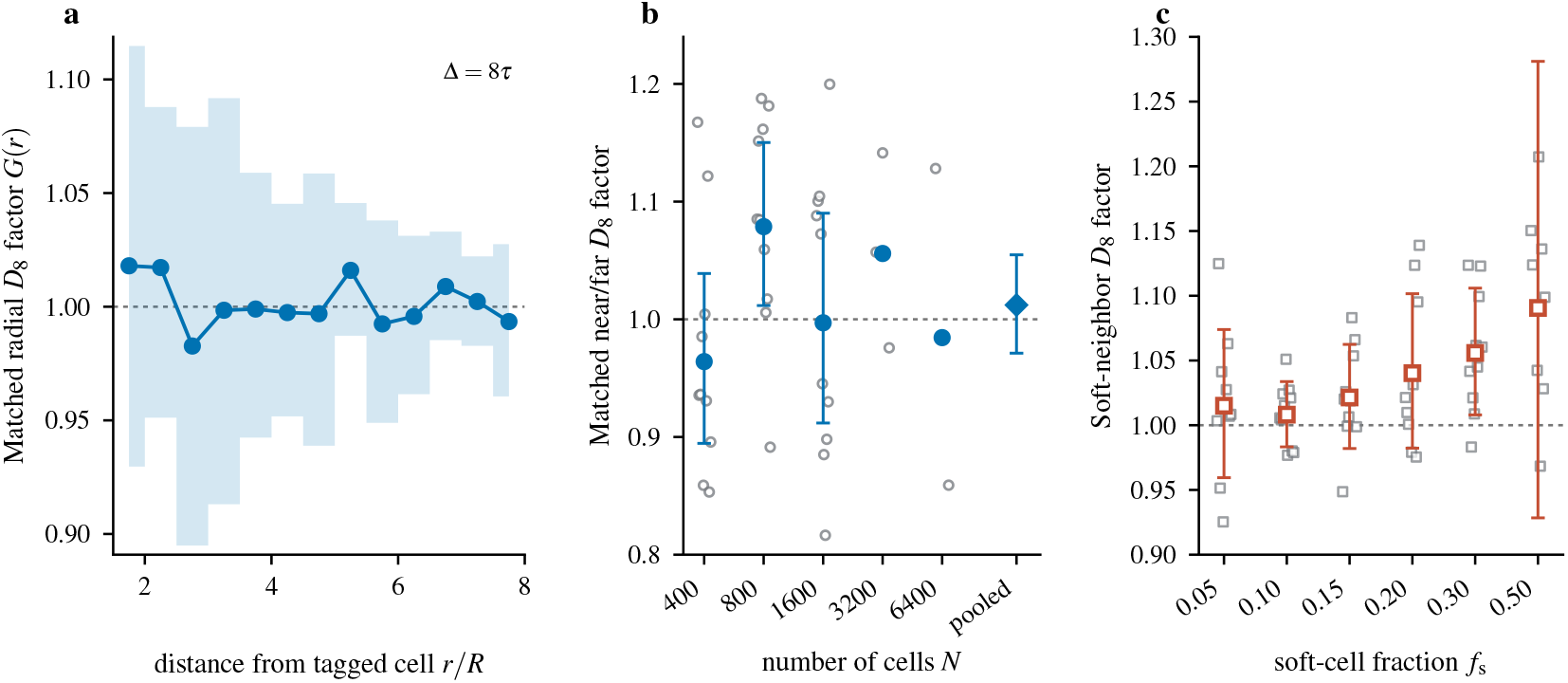
Spatial association of normal-cell transport with soft cells. (a) Distance-resolved comparison of normal-cell *D*_8_ between simulations containing one soft cell and paired all-normal controls. Distances are measured from the tagged cell at the start of each displacement interval. Each simulation’s radial profile is divided by its mean normal-cell *D*_8_ over the entire far region, 6*R* ≤ *r<* 8*R. G*(*r*) is the resulting normalized profile from the single-soft-cell simulation divided by the normalized profile from its paired control. The curve pools 10 paired simulations at each of *N* =400, 800 and 1600. Shading: simultaneous 95% confidence band. (b) The same comparison using the near region 1.5*R*≤ *r<* 3*R* in place of individual distance bins. Open circles: individual pairs; filled circles: geometric means with pointwise 95% confidence intervals for *N* = 400–1600; the diamond: pooled estimate for these three sizes. Estimates at *N* = 3200 and 6400 use three and two pairs, respectively, and are shown without intervals. Unity in (a,b) means that the soft/control transport ratio equals its far-region value. (c) Transport of normal cells with at least one soft Delaunay neighbor relative to normal cells without one in *N* = 400 mixtures. Neighbors are identified at the start of each 8*τ* displacement interval. Each squared displacement is normalized by the normal-cell mean at the same starting time before calculating the ratio of the two group averages. This ratio is divided by the geometric mean of the ratios obtained using random sets of reference cells, each equal in size to the soft population. Gray squares: individual simulations; red squares and bars: geometric means and simultaneous 95% confidence intervals across 10 simulations per fraction.

At finite fractions, normal cells that have a soft Delaunay neighbor have modestly higher *D*_8_ over the following 8*τ* than normal cells without one (Fig. 6c). Across the six fractions, this neighbor/non-neighbor ratio was 1.038±0.025 times the ratio obtained with equal-size random label sets. Its point estimate rised through *f*_s_ = 0.30 and remained above unity at *f*_s_ = 0.50, where the estimate is less precise because few normal cells lack a soft neighbor. This spatial association accompanies the population-wide increase, which is also present in homogeneous monolayers.

## 2 Discussion

Three main results emerge. First, at the baseline advection mobility, crowding opposes propulsion predominantly through field relaxation, including overlap-induced interactions, while uniform interaction advection contributes little to the net transport reduction; interaction advection contributes a few per cent of the opposing displacement product, whereas field relaxation cancels up to ~ 80% of the propulsion velocity at 1–2*τ* after reorientation (Sec. 1.1). Second, changing passive kinetics at fixed free-energy parameters and matched propulsion histories substantially alters transport: reducing *M* by a factor 0.35 (at unchanged free energy) raises the transport of the modified cells 2.2-fold and of the unmodified cells 1.5-fold. Conversely, increasing *M* in the soft cells of binary mixtures removes, and at higher soft-cell fractions reverses, the diffusivity difference between soft and normal cells (Sec. 1.2). Third, normal-cell transport also increases with soft-cell fraction, and homogeneous monolayers at the same mean *γ* reproduce most of the population increase (Secs. 1.5 and 1.6). These findings complement deformability-driven fluidization in multiphase-field monolayers [16, 17] and the shape-controlled unjamming of vertex and Voronoi models [8, 44].

The distinction between energetic and kinetic effects also arises when cell–cell adhesion is altered experimentally. Adhesion contributes both to interfacial energy and to viscous resistance during contact deformation. Bera *et al*. observed increased cell diffusivity after reducing adhesion with little change in cell shape or junctional tension [40]. Their vertex model analysis attributes this increase primarily to reduced viscous resistance, while the parameter governing interfacial energetics remains nearly unchanged.

Optogenetic E-cadherin clustering in the *Drosophila* embryo has been reported to reduce neighbour exchanges by about 70% with no resolved change in cell shape index and myosin levels, and only a small reduction in junctional tension [45]. An anisotropic vertex model qualitatively reproduced this response by increasing junctional viscous friction, whereas the standard energetic implementation of adhesion predicts the opposite. Experiments also show that changing intercellular viscosity alters collective migration without appreciably changing junctional tension [38]. In vertex and MPF models with internal dissipation, reducing substrate friction strengthens spatial correlations between cell velocities [35, 36]. In the vertex models used by Bera *et al*. and Lerchbaumer *et al*., junctional viscous forces oppose relative vertex motion. In the clustering experiments [45] the slowed kinetics are, likewise, junctional: E-cadherin clustering slows the shortening and growth of the very contacts whose myosin-driven remodelling constitutes a rearrangement, and T1 transitions become fewer and slower. The MPF model has no has no junctional friction, 1*/M*_*i*_ is a cell-internal resistance to shape relaxation (Eq. 15) that leaves the imposed propulsion and the advection mobility unchanged. Thus, reducing *M*_*i*_ at fixed free energy parameters acts on the relaxation that cancels propulsion rather than on propulsion itself, and transport increases rather than decreases through the resulting change in the displacement correlations (Sec. 1.2).

Our simulations suggest testing whether slower passive shape recovery at fixed cortical tension and single-cell propulsion increases transport in a confluent monolayer. Recovery measurements, such as those used in junction-rupture experiments [39], could be combined with cell tracking and measurements of junctional dissipation [38]. Mixed monolayers would then test whether changing one population’s recovery also alters transport in the other.

In our binary mixture simulations, normal cells with a soft Delaunay neighbor show modestly higher transport over the following 8*τ* than normal cells without one. This is consistent with local response. In mixed MDCK monolayers, two cell types whose traction and intercellular stress differ when grown separately reach almost the same levels once mixed [46].

Contact-exchange measurements distinguish contact remodeling from centroid transport: deforming cells can change contact length without an equivalent centroid displacement. Faster contact shortening or growth therefore need not imply faster centroid separation during an exchange. The measurements connect relaxation to exchange kinetics, but do not establish that T1 exchanges cause the population transport increase (Sec. S7.2; Fig. S14). Movie 4 illustrates one neighbor exchange in an *N* = 36 monolayer.

In the MPF model studied by Jain *et al*., cell speed on average peaks locally around a T1 exchange, and the relative dispersion of cells increases linearly with the number of T1 events [47], and Palmieri *et al*. traced soft-cell bursts to T1 swaps closing behind the soft cell [21]. In MPF monolayers, where successive T1s occur depends on cell shape: a cell that loses a neighbor becomes rounder, one that gains a neighbor becomes more elongated, and the resulting asymmetry biases cells towards directed migration [48].

The overlap contribution connects reduced slowing to the repulsive-particle theory of Bialké *et al*. [49], in which density-dependent slowing of self-propelled particles drives motility-induced phase separation [50, 51]. In the limit of identical circular profiles with common kinetic coefficient and field-weighted area *V* =*∫*_Ω_ *q*_*i*_ d*A*, overlap is centered halfway between the centroids. With *O*(*r*) =*∫ q*_*i*_*q* _*j*_ d*A*, Eq. 6 gives

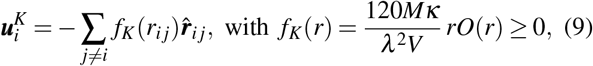

where 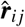 points from cell *i* to *j*. Let *n* denote the mean cell number per unit area and *g*(*r, θ*) the pair distribution of cell centroids at separation *r* with *θ* measured from cell *i*’s propulsion direction 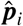. Averaging over the corresponding neighbor density *ng*(*r, θ*) gives

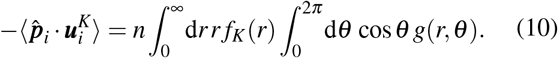

This has Bialké’s force-imbalance form: stronger repulsive contributions ahead than behind cause slowing, and reducing the imbalance weakens opposition. At packing fraction 0.5 and Péclet number ≈ 15–75, deformability instead promotes motility-induced phase separation in MPF suspensions. This occurs because deformable cells stay in contact for longer times during two-body collisions when interaction forces enter the advection velocity [18]. Here, at packing fraction 0.90 and Pe = *l*_*p*_*/R*≈ 2 (*l*_*p*_ = *v*_*A*_*τ* for run-and-tumble propulsion [52]), opposition is carried by field relaxation rather than by interaction advection (Sec. S2.3). Hopkins *et al*. also found that without passive forces in the advection velocity, more deformable cells are slowed less by crowding [18]. Here *f*_*K*_ in Eq. 10 describes relaxation-induced centroid velocity, and interaction advection remains separate (Sec. S7.3). For deforming cells, Eq. 6 retains the actual overlap geometry, and Eq. 7 retains its time-dependent contribution to transport.

At fixed 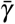, write 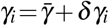 with Σ_*i*_ *δγ*_*i*_ =0. For sufficiently small contrast,

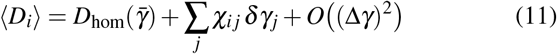

where 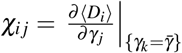. The coefficient *χ*_*i j*_ measures the response of the mean transport of cell *i* to a change in *γ* _*j*_. In the homogeneous reference, this response summed over all cells *i* is the same for every perturbed cell *j*, i.e., Σ_*i*_ *χ*_*i j*_ = *c* is independent of *j*. Averaging Eq. 11 over cells then gives a linear contribution (*c/N*) Σ _*j*_ *δγ*_*j*_ = 0 at fixed 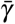, whereas the difference between the two classes can contain a linear term. The mixture and the homogeneous monolayer at 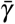 therefore have the same population transport to first order in Δ*γ*. We ask whether a different mean interfacial coefficient can improve the agreement between the homogeneous reference and the measured population transport of the mixture. For a homogeneous reference at *γ*_*g*_ = *g*^−1^(*g*(*γ*_*i*_)) with smooth monotone *g*, we obtain

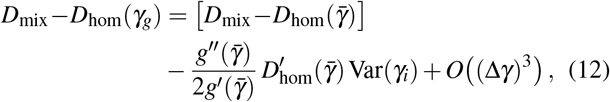

where Var(*γ*_*i*_) = *f*_*s*_(1 −*f*_*s*_)(Δ*γ*)^2^. Combining Eq. 12 with the geometric-mean description *D*_mix_ ≃ *D*_hom_(*γ*_*G*_) in Sec. 1.6 predicts, for small contrast,

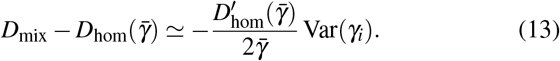

The predicted excess over the arithmetic-mean reference is positive because 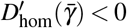, and its magnitude is set by the coefficient variance and the slope of the homogeneous transport curve. At uniform *M*_*i*_, geometric averaging gives the same homogeneous coefficient when applied to the interfacial rates or the corresponding relaxation times (Eq. 4). Within the sampled range, this relation predicts equal changes in population transport for equal fractional changes in any cell’s interfacial rate.

Mechanical heterogeneity can alter rigidity [53] and produce small-scale demixing [54] in vertex models, and fluctuating junctional adhesion is a further route to fluidizing MPF monolayers [55]. The transport increase in our mixtures occurs without positive soft-cell sorting (Fig. 5b). Different imposed noise strengths can drive demixing in mixtures of Brownian disks [56, 57]. McCarthy *et al*. found no large-scale demixing at high packing fractions or in their confluent Voronoi model [57]. Here, the ≈12% difference in *D*_8_ arises from mechanical heterogeneity under identical propulsion parameters. The absence of positive sorting also differs from the small scale demixing of shape-disparate vertex model mixtures [54]. Equation 14 contains no heterotypic interfacial term, and the overlap coefficient *κ* is the same for all cell pairs. Cell sorting has been explained by differential adhesion [58, 59], while an extra tension at contacts between unlike cells can keep the boundary between two confluent populations sharp [60].

In a confluent phase-field monolayer, Graham *et al*. found small-scale sorting between cells with extensile and contractile intercellular active stresses, and attributed this sorting to different cell diffusivities [61]. The late-time mean-square displacements gave an extensile-to-contractile ratio of ≈2.7 in their initially mixed system (estimated from their Fig. 3b). Segregation that depends on activity has also been found in a phase-field model in which elasticity is the only difference between the two cell types [62]. Growth is a further route: in a proliferating force-based tissue, soft cells with a higher growth potential form a compact domain that displaces the stiff matrix [63]. Our monolayers have neither growth nor division, and the transport difference alone produces no sorting. Kinetic and dissipative parameters also change collective motion in other phase-field [36, 37, 64] and vertex models [35, 65], and MPF collective behavior depends qualitatively on how activity is implemented [66]. Here 1*/M*_*i*_ is an internal shape-relaxation resistance, distinct from the intercellular friction that solidifies MPF monolayers [37] and from the substrate friction parameter that sets the advection mobility (Eq. 16). Biological mixtures also differ in adhesion, nuclear mechanics and traction [19, 46]. Here, evaluating the field equations for the same saved simulation snapshots shows that a lower soft-cell interfacial coefficient reduces the mean initial growth rate of 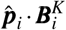 in normal cells. The displacement correlations measured in the evolving mixtures connect reduced relaxation-induced motion opposite propulsion to increased transport.

## 3 Methods

### 3.1 Active phase-field model

Cell *i* is represented by a scalar field *ϕ*_*i*_(***r***,*t*), with *ϕ*_*i*_ ≃1 inside the cell and *ϕ*_*i*_ ≃ 0 outside [21, 27, 28]. The free energy over the periodic domain Ω combines interfacial energy with penalties for deviations from the target cell area and for overlap between cells:

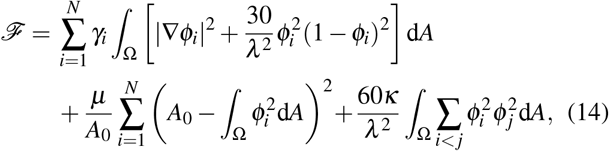

where *A*_0_ = *πR*^2^ is the target area, *λ* sets the interface width, *µ* controls the penalty for changing the cell area, and *κ* the penalty for overlap between cells. The fields obey non-conserved (model A) relaxation with advection [21, 33],

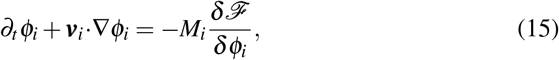

where *M*_*i*_ is the kinetic coefficient for field relaxation. The advection velocity is uniform within each cell and is given by

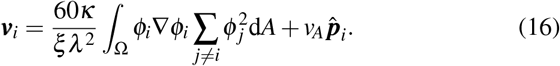

Here *v*_*A*_ is the propulsion speed and 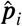 the polarity direction. The interaction-induced velocity equals the force associated with a rigid cell translation divided by 2*ξ* (2*ξ* follows the convention of Palmieri *et al*.). The advection prefactor 1*/ξ* is independent of *M*_*i*_. Reducing *M*_*i*_ therefore slows shape relaxation at fixed advection mobility. Without propulsion, both field relaxation and interaction advection dissipate *F* (Sec. S2.1, Eqs. S5–S6).

Each cell’s propulsion direction remains fixed between reorientation events, which follow a Poisson process with mean waiting time *τ* [21, 52]. We use *τ* = 10^4^ in model time units. At each timestep of duration *δt*, the cell reorients with probability *p*_t_ = −1 exp(− *δt/τ*). After an event, the new angle is drawn from a uniform random distribution in [0, 2*π*). The time between successive reorientation events is denoted by *T*_run_. Then, for the underlying Poisson process and *s* ≥0, *P*(*T*_run_ *> s*) = *e*^−*s/τ*^ and 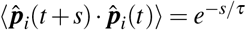. Numerical evaluation of this probability, and the checks that paired simulations share the same polarity histories, are described in Secs. S3.2, S6 and S8.

Results are given in model units. Using the calibration of Refs. [21, 23], one model time unit corresponds to about 0.36 s, so *τ*≈ 1 h and 200*τ* corresponds to about eight days. The model does not have cell growth or division. The values *γ*_n_ = 1 and *γ*_s_ = 0.35 follow Palmieri *et al*. [21]. Unless stated otherwise, we use *M*_*i*_ = 0.5, *R* = 49, *λ* = 7, *κ* = 10, *ξ* = 1500, *µ* = 1, *v*_*A*_ = 0.01 and *τ* = 10^4^. The relaxation of the interface profile is characterized by *τ*_*ϕ,i*_ below, while *τ*_2,*i*_ characterizes relaxation of an elliptical cell deformation (Eq. 4).

Next, we derive the stationary planar-interface profile, line tension and intrinsic interfacial relaxation time from Eqs. 14 and 15. Minimizing the interfacial energy gives a stationary profile independent of *γ*_*i*_, which multiplies both interfacial terms in Eq. 14:

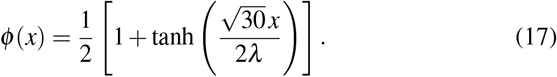

The interface width is fixed by *λ*. The planar equilibrium line tension and the intrinsic interfacial relaxation scales are

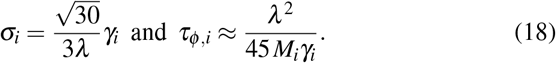

The first relation follows from integrating the interfacial energy using the profile in Eq. 17, and the second follows by linearizing the interfacial relaxation dynamics about that profile [41]; the derivation is given in Sec. S3.1. The eigenvalues of the linearized operator give the decay rates of small perturbations to the planar profile. The slowest decaying profile mode has rate 45*M*_*i*_*γ*_*i*_*/λ* ^2^, while the continuum of faster modes begins at 60*M*_*i*_*γ*_*i*_*/λ* ^2^. Thus *τ*_*ϕ,i*_≈ *λ* ^2^*/*(45*M*_*i*_*γ*_*i*_) ≈2 model time units for normal cells.

Changes in the overall cell shape relax more slowly. In the sharp-interface limit, the interfacial term of Eq. 15 gives the Allen–Cahn normal velocity *v*_⊥_ = −2*M*_*i*_*γ*_*i*_*/ρ* for a boundary with local radius of curvature *ρ* [34]. For an area-constrained circular cell of radius *R*, a distortion of angular order *m* decays with *τ*_*m*_ = *R*^2^*/*[2*M*_*i*_*γ*_*i*_(*m*^2^ −1)]; the elliptical mode has *m* = 2 (Eq. 4). At fixed *R* and *λ*, both *τ*_*ϕ,i*_ and *τ*_2,*i*_ scale as 1*/*(*M*_*i*_*γ*_*i*_). In the compensated control, *M*_s_ is increased so that *M*_s_*γ*_s_ = *M*_n_*γ*_n_, keeping both relaxation times equal to their normal-cell values. The corresponding free-cell persistence length is *l*_*p*_ = *v*_*A*_*τ* = 100 ≃ 2.04*R*, that is, approximately one cell diameter.

### 3.2 Numerical simulations

The *N* cells are simulated in a periodic square box of side length *L*, chosen so that *NπR*^2^*/L*^2^≃ 0.90 [21]. The monolayer is then near-confluent, with small gaps at multicellular junctions (Fig. 1). Time integration uses the forward Euler method with timestep *δt* = 0.01 on a unit-spaced periodic lattice, and spatial derivatives use centered first differences and an isotropic nine-point Laplacian [67]. We use a GPU-accelerated phase-field solver implemented in CUDA C++, building on the CUDA backend of SymPhas 2.0 [68]. Fields and pointwise stencil operations use single-precision arithmetic, and cell areas, moments and interaction-force sums are accumulated in double precision. Production calculations used Nvidia H100, A100 and GH200 GPUs.

The initial cell centers were placed at random. Each center initialized a circular phase field (Sec. S8, Eq. S38). During preparation, propulsion was disabled (*v*_*A*_ = 0), while field relaxation and interaction-driven advection remained active. The simulations at different soft-cell fractions *f*_*s*_ started from passively relaxed all-normal configurations. A randomly selected subset of *f*_*s*_*N* cells was assigned *γ*_s_, followed by a further 1*τ* of passive relaxation and 50*τ* of active motion.

Parameter assignments for the mixtures and controls are specified in Secs. 1.2 and 1.6, and Sec. S3.1 and Table S2.

Paired conditions shared the initial fields and selected cell identities before passive (*v*_*A*_ = 0) preparation, and used identical imposed polarity histories during active motion. Table S4 lists the system sizes, compositions, replicate counts and active durations. The low-fraction and paired-control preparation procedures are described in Sec. S8.1. A double-precision control implementation reproduced one update with a maximum relative *L*_2_ field difference of 4.81 ×10^−8^. Timestep sensitivity, domain-support and restart checks are reported in Sec. S8.2 and Table S5.

### 3.3 Observables and statistical analysis

Cell positions are 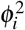-weighted centroids. We evaluate *D*_8_ at Δ = 8*τ* (=80000), the shorter end of the range 8*τ*–16*τ* over which the tagged-cell MSD is approximately proportional to lag (Fig. S1). The enhancement and its dependence on system size and composition persist across the tested lags (Sec. S1, Fig. S2), and the free-cell expression is Eq. S2.

Uncertainties are given as 95% confidence intervals. Square brackets give the lower and upper bounds when the interval is asymmetric. Each independently generated initial condition defines one statistical replicate. Ratios between paired conditions were calculated within each replicate and summarized by geometric means with two-sided Student-*t* intervals calculated on the logarithmic scale. Composition means and slopes were likewise calculated within each simulation before combining independent replicates. The contributions to diffusivity are reported as arithmetic means across replicates with pointwise Student-*t* confidence intervals. For control comparisons, the differences were first calculated within each pair over common observation windows.

For the spatial analysis, the distance to the tagged cell was measured at the beginning of each displacement interval. The near and far regions were 1.5*R* ≤ *r <* 3*R* and 6*R* ≤ *r <* 8*R*, respectively. The near region extends to the first minimum of the radial distribution function, while the far region spans three to four nominal cell diameters. The pooled near-to-far comparison remains consistent with unity when the region boundaries are varied (Fig. S15). We calculated the near-to-far transport ratio in each simulation, then divided the single-soft-cell ratio by its paired all-normal ratio to account for the radial structure already present around the tagged cell. Distance-resolved analysis, random-label comparisons and simultaneous confidence bands are described in Secs. S9 and S10, Eqs. S40–S42.

## Acknowledgements

The authors thank Profs. Gonca Erdemci-Tandogan and Matteo Rauzi for helpful suggestions and critical reading of the manuscript. M.K. thanks the Foundation PS and the Research Council of Finland Flagship of Advanced Mathematics for Sensing, Imaging and Modeling (grant 358944), and S.A.S. the NSERC Postgraduate Scholarships–Doctoral program. Computational resources were provided by the Digital Research Alliance of Canada and CSC–IT Center for Science, Finland.

## Author contributions

S.A.S. and M.K. designed research and analyzed data; S.A.S. performed the simulations, developed software, and performed the numerical analysis of the data; S.A.S. wrote the first draft of the paper.

## Data and code availability

The simulation code and parameters are publicly available at the GitHub repository.

## Supplementary Information

### S1 Observation-lag sensitivity

We express time in units of the polarity-persistence time *τ* = 10^4^ and report the finite-lag diffusivity *D*_8_ = *D*(8*τ*). The choice Δ = 8*τ* is motivated in Methods, and here we document the lag dependence. For lag Δ, let *t*^′^ be the stored time nearest *t* + Δ. We calculate

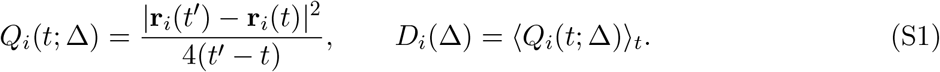

We average over overlapping time windows within the active observation period [1]. Positions are unwrapped across periodic boundaries. Displacements are measured in the laboratory frame without subtracting the population drift.

The free-cell result provides a reference for lag dependence arising from propulsion persistence alone. For free motion at speed *v*_*A*_ with independent isotropic Poisson reorientations, 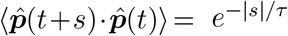. Integrating this correlation gives 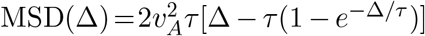 and 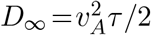, hence

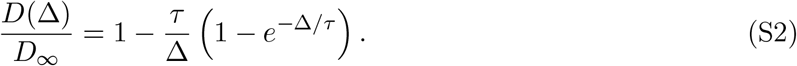

At Δ = 4*τ* and 8*τ*, *D*(Δ)*/D*_∞_ reaches 0.755 and 0.875, respectively. Figure S1 shows the pooled tagged-cell MSD and finite-lag diffusivity. The log–log slope between 8*τ* and 16*τ* is 1.04 in both the all-normal and single-soft-cell systems. The pooled tagged-cell enhancement persists across lags (Fig. **??**c). Figure S2 shows that the absence of a resolved system-size dependence also persists. Transport increases with soft-cell fraction for both cell types at every lag tested. The difference between their slopes remains consistent with zero within uncertainty. Restricting the 8*τ* displacement intervals to the first or second half of each composition trajectory gives no resolved early-to-late change in the soft-cell slope, normal-cell slope, or their difference. The *N* = 3200 and 6400 comparisons also span 200*τ* of activity and are shown individually. The pooled estimate and system-size regression use *N* = 100–1600.

### S2 Centroid displacement and relaxation

#### S2.1 Derivative of the field-defined centroid

To separate motion due to advection from motion due to field relaxation, we differentiate the centroid defined by the phase field. Let 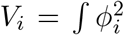 d*A* and 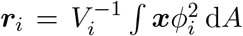. Differentiating both the numerator and its normalization gives

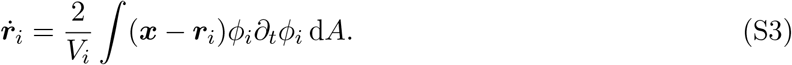

The advection velocity is spatially uniform within each cell. Substituting Eq. **??** and integrating the advection term by parts with a vanishing surface term for a localized cell field, gives

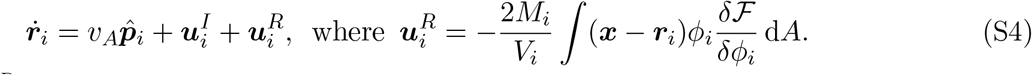

Here 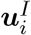 and 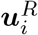 are the contributions to centroid velocity from interaction advection and field relaxation, respectively. The area penalty for cell *i* in Eq. **??** is *µ*(*V*_*i*_ *A*_0_)^2^*/A*_0_. Since *δV*_*i*_*/δϕ*_*i*_ = 2*ϕ*_*i*_, its variational derivative is 4*µ*(*V*_*i*_ *A*_0_)*ϕ*_*i*_*/A*_0_. Its direct contribution to Eq. S4 vanishes because 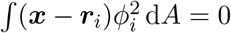 by the definition of the centroid. Area regulation enters the centroid dynamics through the evolution of *ϕ*_*i*_.

##### Advection convention and passive dissipation

The force associated with rigid translation of a cell determines its interaction-advection velocity. With *ψ*_*i*_ = *δ*ℱ*/δϕ*_*i*_, this force and velocity are

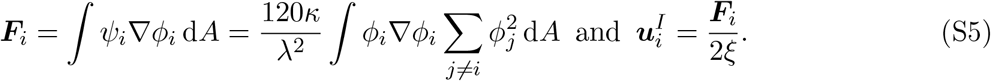

The self-interfacial and area energies are translationally invariant, and thus only the overlap term contributes to this force. With Palmieri *et al*.’s convention [2], the effective translational friction is 2*ξ*. The parameter *ξ* sets the advection prefactor, proportional to 1*/ξ*, independently of *M*_*i*_. Changing *M*_*i*_ can therefore alter 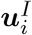 through field evolution while leaving the advection coefficient fixed. Both advection and field relaxation dissipate free energy in the absence of propulsion. With *v*_*A*_ = 0, the continuum dynamics obey

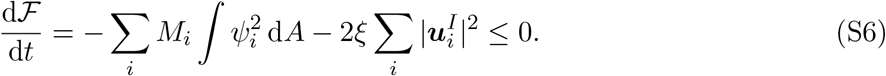

Numerical tests at finite timesteps are reported in Sec. S8.2.

#### S2.2 Signed contributions and their interpretation

To determine the contribution of each velocity to transport, we integrate it over a displacement interval before forming the mean-square displacement. We define 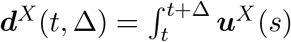 d*s* with *X* = *A, I, R* for propulsion, interaction advection and physical field relaxation, respectively, and 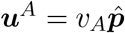. Summing these displacement contributions gives

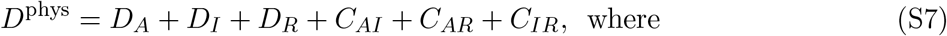

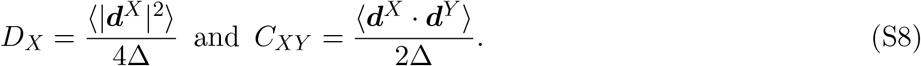

Positive cross-terms increase the mean-square displacement, whereas negative cross-terms reduce it. We evaluate these averages without subtracting the mean displacements. Transport contributions are reported in units of 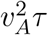. The free-cell value at the same lag is derived in Sec. S1.

Interaction advection translates the cell field uniformly through the net overlap force. Passive relaxation changes the field locally and includes the overlap term in ℱ. Its contribution to the propulsion cross term is

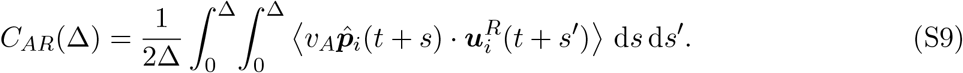

For stationary dynamics, *C*_*AR*_ can be written in terms of *K*_*αβ*_(*s*) = ⟨***u***^*α*^(*t*) · ***u***^*β*^(*t* + *s*)⟩ as

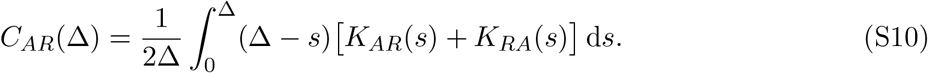

We calculate the displacement products directly, using Eq. S9 without a stationarity approximation. Palmieri *et al*.’s supplementary ellipse calculation relates short speed bursts to centroid shifts during relaxation with one side constrained [2]. Yin *et al*. relate speed peaks to preceding deformation [3]. The derivation in Sec. S4 extends the centroid calculation to arbitrary shapes and all displacement windows. Section S7.3 derives the connection between overlap-induced opposition and the repulsive-particle force imbalance of Bialké *et al*. [4].

#### S2.3 Numerical evaluation and inference

The transport analysis covers binary mixtures, homogeneous systems at the same mean *γ*, half-contrast mixtures and compensated mixtures at soft-cell fractions *f*_*s*_ = 0.05, 0.10, 0.15, 0.20, 0.30, 0.50. The conditions are defined in Secs. **??** and **??** of the paper and Table **??**. The control with reduced *M*_*i*_ uses 30% modified cells. We also analyze all-normal references. All simulations had *N* = 400 and a 50*τ* active observation window with polarity-persistence time *τ* = 10^4^.

We integrate the recorded propulsion and interaction-advection velocities over the same time intervals used to calculate cell displacements. Subtracting these two contributions from the total cell displacement gives the relaxation estimate

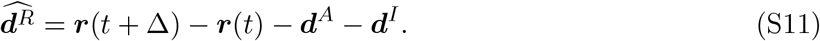

The hat denotes the estimate obtained from the recorded displacements and velocities. The relaxation velocity is 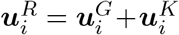, where 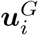 and 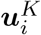 are the interfacial and overlap contributions, respectively.

We use the estimated displacement to calculate *D*_*R*_ and *C*_*AR*_ for the control comparisons and the analysis in Sec. S5.2. When the interfacial and overlap velocities are recorded separately, we also integrate their sum and compare the resulting displacement with this estimate.

For each simulation, we averaged over cells and over all available starting times for the displacement intervals. For paired comparisons, we calculated the contribution difference within each seed and report the arithmetic mean with a pointwise 95% Student-*t* interval. Positive transport ratios are reported as geometric means. Compensated mixtures and the control with reduced *M*_*i*_ use the same designated cell indices and recorded polarity histories as their all-normal references.

The principal responses persist at lags 4*τ*, 8*τ* and 16*τ*. We tested the numerical integration of the recorded propulsion and interaction-advection velocities using trapezoidal, left-endpoint and right-endpoint rules, and repeated the trapezoidal calculation using every second or fourth recorded frame. At 8*τ*, these variations change the mean *D*_*R*_ and *C*_*AR*_ differences for the control with reduced *M*_*i*_ and the compensated mixtures at 30% by at most 2.63 × 10^−5^, preserving their signs. Independent displacement, correlation and polarity-partition calculations agree within 1.4 × 10^−13^. For the separately recorded physical sources, the normal-cell transport gain at 50% differs from the sampled-centroid gain by 1.82 × 10^−4^, or 1.06% of the centroid gain (Sec. S7). Table S1 gives the displacement balance obtained from the directly recorded velocity contributions in the all-normal reference simulations.

We calculate *D*_*A*_ from the recorded propulsion histories. The ideal free-cell expression is in Sec. S1. Centroids measured over the same windows give *D*_8_ = 0.04157 ± 0.00091, which is 0.00051 below the sum of the velocity contributions. The net relaxation reduction −(*D*_*R*_ + *C*_*AR*_) accounts for 99.68% of 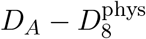 (the mean of seed-level shares). Using only the negative propulsion cross terms gives *C*_*AR*_*/*(*C*_*AR*_ + *C*_*AI*_) = 98.33%.

##### Simulation sets and relaxation estimates

The all-normal balance above, and the binary-minus-all-normal comparisons, use directly recorded interfacial and overlap velocities. For comparisons between binary-mixture fractions, we estimated relaxation from centroid displacement and recorded advection, using the 5% mixture as the reference. The simulation sets and analysis windows are listed in Sec. S8.

#### S2.4 Relaxation after reorientation

We measure how relaxation opposes propulsion after a cell reorients, using 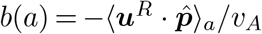 calculated from the recorded interfacial and overlap contributions. The simulations and averaging conventions are specified in Sec. S7. Time *a* = 0 is the first recording of the new direction. We include measurements only until the next reorientation. Within each time bin, we first average the measurements for each cell, then average equally over the contributing cells. Let *b*_0_ be the first resolved value in 0–0.025*τ* and *b*_late_ the equal mean of the 1–1.5*τ* and 1.5–2*τ* bins. We normalize as 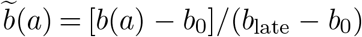 and interpolate between bin midpoints to find its 50% and 90% crossing times, *t*_50_ and *t*_90_. In the 10 distinct all-normal populations, *b*_0_ = 0.5085 ± 0.0011 and *b*_late_ = 0.8138 ± 0.0023 with pointwise 95% Student intervals across seed means. The projection therefore cancels about 50.8% of *v*_*A*_ in the first resolved 0–0.025*τ* bin and 81.4% at ages 1–2*τ*. The first value is an average over a finite recording bin, not the instantaneous response at the turn. The mean relaxation-velocity projection along the new polarity, normalized by *v*_*A*_, is denoted by 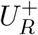. Its curves in Fig. S3 use narrower time bins immediately after reorientation and weight each recorded interval by its duration.

The normal-cell late level decreases from 0.807 at 5% to 0.755 at 50%, while all-normal levels remain near 0.814. The 50% crossing occurs earlier in mixtures at *f*_*s*_ ≥ 0.10. All paired pointwise intervals for the *t*_90_ ratio include one. At 50%, the interfacial and overlap contributions to late opposition decrease by 0.029*v*_*A*_ and 0.028*v*_*A*_, respectively. The response curves and crossing times are shown in Fig. S4.

### S3 Passive relaxation coefficients and control comparisons

#### S3.1 Passive coefficients and relaxation scales

To compare field relaxation in the binary mixtures, compensated mixtures and control with reduced *M*_*i*_, we identify the coefficients multiplying the interfacial, overlap and area contributions. Write ℱ = ∑_*i*_ *γ*_*i*_ℰ _*i*_ + *κ* ℰ_*K*_ + *µ*ℰ_*V*_. The passive field equation contains the coefficients *M*_*i*_*γ*_*i*_, *M*_*i*_*κ* and *M*_*i*_*µ*. Relative to a normal cell (*γ* = 1, *M* = 0.5), the interfacial and overlap/area coefficient ratios are listed in Table S2.

A cell’s relaxation also depends on the neighboring fields, even when its own material coefficients remain fixed. At fixed parameters, a small variation of the fields gives

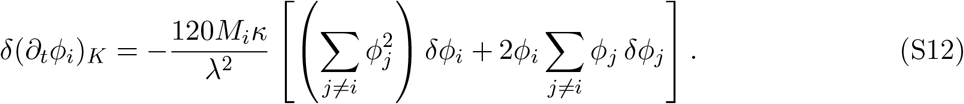

The term proportional to *δϕ*_*j*_ couples deformation of neighboring cells to cell *i*’s relaxation.

##### Planar-interface and cell-scale relaxation

For a planar equilibrium interface, integration of the interfacial energy gives the line tension 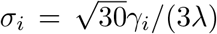. A small perturbation of this profile evolves according to *∂*_*t*_*δϕ*_*i*_ = −ℒ_*i*_*δϕ*_*i*_ when only interfacial relaxation is retained. Writing 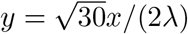, the linearized relaxation operator is

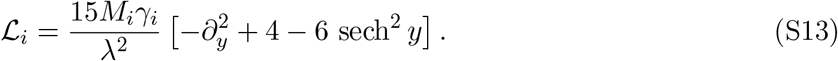

The eigenvalues of ℒ_*i*_ give the decay rates of the perturbations. The translation mode sech^2^ *y* has eigenvalue zero because it shifts the interface without changing its profile. The slowest decaying profile mode is sech *y* tanh *y* with eigenvalue 45*M*_*i*_*γ*_*i*_*/λ*^2^. The continuum of faster modes begins at 60*M*_*i*_*γ*_*i*_*/λ*^2^. The corresponding profile-relaxation time is *τ*_*ϕ,i*_ = *λ*^2^*/*(45*M*_*i*_*γ*_*i*_), approximately 2.18 model time units for a normal cell.

For changes in the overall shape of an area-constrained circular cell, the sharp-interface approximation gives *τ*_*m,i*_ = *R*^2^*/*[2*M*_*i*_*γ*_*i*_(*m*^2^ −1)] for angular modes *m* ≥ 2. The *m* = 2 mode describes an elliptical deformation with *τ*_2,*i*_ ≃ 800 model time units for a normal cell. These are reference times for an isolated interface and an area-constrained cell. Relaxation in the crowded monolayer also involves interactions with neighboring cells.

#### S3.2 Robustness to increased advection mobility

To test whether relaxation remains the dominant contribution to the transport gain when interaction advection is stronger, we repeated the control with *M*_*i*_ reduced in 30% of cells and its all-normal reference at *ξ* = 375, reducing the baseline value by four and multiplying the interaction-advection prefactor by four. Ten matched seeds each supply two *N* = 400 recordings of the control with reduced *M*_*i*_ and its all-normal reference as defined in Sec. **??** (120 modified cells). Starting fields before the additional 1*τ* passive preparation and cell identities are shared, followed by 50*τ* of activity. For each seed, the simulations with reduced *M*_*i*_ and the all-normal simulations use the same propulsion histories at both values of *ξ*.

The primary centroid estimates use every complete 8*τ* window within 0.01–50*τ* of activity. The geometric transport ratios relative to the all-normal reference are 2.105 [2.050, 2.162] for modified cells, 1.457 [1.407, 1.509] for unmodified cells and 1.653 [1.608, 1.699] for the population (pointwise 95% Student intervals across 10 log ratios, Fig. S5a). The same 10 seeds at *ξ* = 1500 give 2.246, 1.498 and 1.723, respectively.

The paired ratios of enhancements, 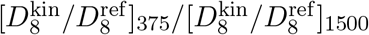, are 0.937 [0.912, 0.964], 0.973 [0.937, 1.010] and 0.959 [0.933, 0.986] for modified cells, unmodified cells and the population.

Their point estimates decrease by 6.3%, 2.7% and 4.1%.

We calculate the relaxation displacement by integrating the interfacial and overlap velocities and retain all six terms in Eq. S7. Define *R*_net_ = *D*_*R*_ + *C*_*AR*_ and *I*_net_ = *D*_*I*_ + *C*_*AI*_ + *C*_*IR*_. Population means for the control with reduced *M*_*i*_ and its all-normal reference at *ξ* = 375 are given in Table S3. Figure S5 shows confidence intervals and changes for each cell type. For modified cells, *δR*_net_ = +0.06833 and *δI*_net_ = − 0.02545: interaction advection offsets part of the relaxation-associated gain. For unmodified cells these changes are +0.01606 and +0.00129, and for the population they are +0.03174 and − 0.00673. Since *δD*_*A*_ = 0, the two changes sum to the transport change calculated from the recorded velocity contributions within every seed. Thus relaxation remains the dominant contribution to the gain, while the interaction-advection contribution differs quantitatively from the baseline. In the control with reduced *M*_*i*_, the mean shares *C*_*AR*_*/*(*C*_*AR*_ + *C*_*AI*_) are 83.2% for modified cells and 93.6% for unmodified cells. The corresponding reference shares are both 93.7%.

Integrating the recorded velocity contributions gives population transport estimates of 0.06364 and 0.03863, compared with centroid estimates 0.06304 and 0.03815 over the same windows. The two estimates differ by approximately 1%. Checks of the transport calculations are given in Sec. S8.2.

### S4 Physical origin of relaxation-induced centroid motion

The relaxation velocity 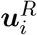 in Eq. S4 contains interfacial and overlap contributions. Each contribution is obtained by weighting the corresponding local rate of change of 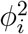 by displacement from the centroid, integrating over space and dividing by the cell area. Writing 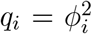 and ***s***_*i*_ = ***x*** − ***r***_*i*_, Eq. **??** gives

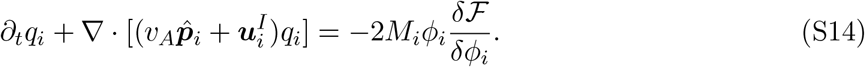

The overlap term in Eq. **??** and its contribution to Eq. S14 are

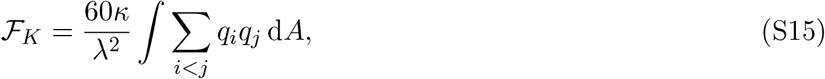

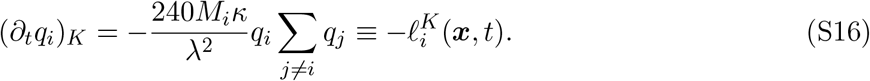

Thus 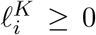 is the local rate at which overlap relaxation decreases 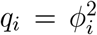, the field used to calculate the centroid. Let 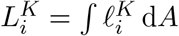 and, when 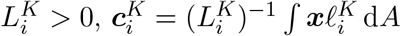. The overlap contribution to centroid velocity is

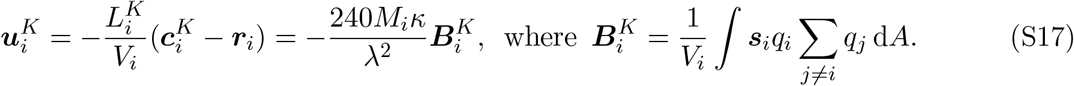

At zero overlap the integral gives 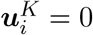. Otherwise this relaxation contribution shifts the centroid away from the overlap-weighted center 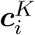. Overlap concentrated ahead of the centroid therefore makes overlap relaxation oppose propulsion. An overlap distribution symmetric about the centroid gives 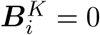.

For ℱ_*G,i*_ = *γ*_*i*_ℰ_*i*_ with 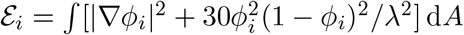 the interfacial contribution is

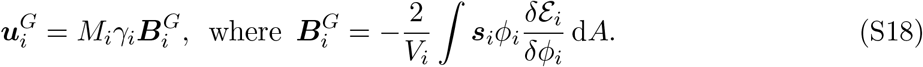

Interfacial relaxation can increase or decrease *q*_*i*_ locally. The area term makes no direct contribution to the centroid velocity, so 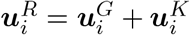 in the continuum equation.

To relate overlap relaxation to finite-lag transport, we integrate its velocity over the displacement window 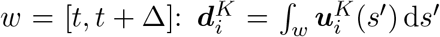. The propulsion displacement is 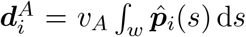.

Substituting Eq. S17 into 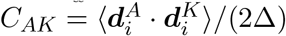 then gives

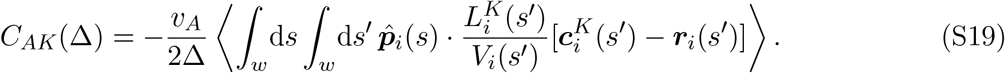

Equation S19 connects 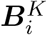 to transport through its correlations with propulsion at all times in the window. Together with interfacial relaxation, *C*_*AR*_ = *C*_*AG*_ + *C*_*AK*_ and *D*_*R*_ = *D*_*G*_ + *D*_*K*_ + *C*_*GK*_. Their contributions to transport are combined with propulsion and interaction advection in Eq. S7.

#### S4.1 Response to neighboring interfacial relaxation

The saved-configuration test in main Fig. **??**c uses *N* = 400 binary mixtures with ten seeds at each soft-cell fraction. We average over normal cells within each seed and then over seeds. To isolate coupling through neighbors, we evaluate the field equation twice at the same saved configuration *t*_0_: with the binary coefficients and with only soft cells’ *γ* restored from 0.35 to 1. Every field, polarity and other parameter is held fixed. Write 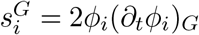 for the interfacial source in *∂*_*t*_*q*_*i*_ and *δ*_*γ*_ for restored minus original coefficients. A normal cell has the same instantaneous source and centroid velocity in both evaluations, but the neighboring sources alter the initial rate of change of 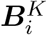:

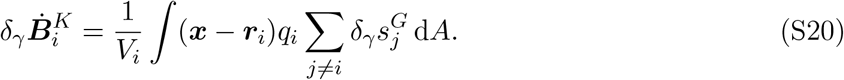

The mean forward projection 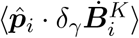 is positive at the saved active configurations at all six fractions.

For normal cells, the initial fields and the overlap coefficient are identical, so 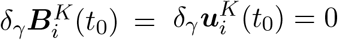. For a short time *h* before any subsequent orientation event, Eq. S17 gives

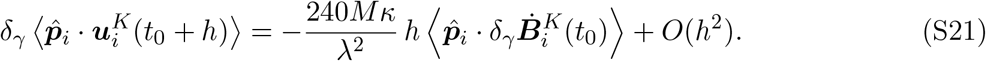

At short times, restoring the neighbors’ interfacial coefficient therefore strengthens the overlap contribution opposing propulsion.

##### Sustained relaxation in the evolved mixtures

At ages 1–2*τ* after a normal cell reorients, both *G* and *K* produce smaller backward velocities in binary mixtures than in all-normal controls with paired pointwise intervals excluding zero at every fraction. The fixed overlap coefficient makes the reduction in backward *K* velocity proportional to the reduction in the projection of 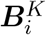 along propulsion.

### S5 Temporal correlations of cell motion

#### S5.1 Time correlations and the persistence of centroid motion

We resolve displacement products by time separation to test which correlations produce the transport change. This analysis covers binary and compensated mixtures across all fractions and the control with reduced *M*_*i*_, all-normal references, homogeneous controls and half-contrast mixtures at 30%. Displacement intervals begin throughout the active trajectory and are not aligned with reorientation events. The following calculations use *A, I, R* for propulsion, interaction advection and relaxation with the latter evaluated by Eq. S11. Let 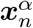 denote the displacement due to contribution *α* between recordings *n* and *n* + 1. Summing the propulsion, interaction-advection and relaxation contributions gives the cell displacement over that interval. For the set of displacement origins *s* and endpoints *e*_*s*_, define

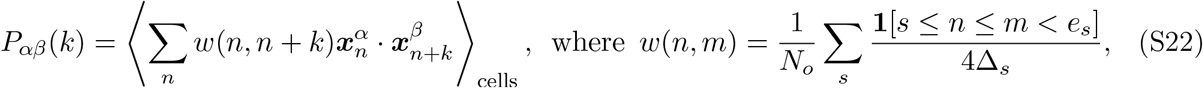

where *N*_*o*_ is the number of origins and Δ_*s*_ is the actual duration of the corresponding displacement interval. These weights give

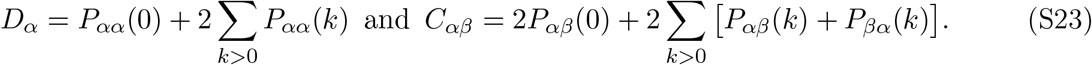

We evaluate the weighted displacement products using zero-padded Fourier correlations and check the results against explicit pair sums.

We also separate the contributions of centroid speed and persistence to transport. For each class within each seed, *S*_*v*_ is the mean square of centroid velocities averaged over the recorded time intervals, and *T*_*v*_(Δ) = 2*D*(Δ)*/S*_*v*_. Here *T*_*v*_ measures the finite-lag integral of the normalized velocity correlation. The identity *D* = *S*_*v*_*T*_*v*_*/*2 holds separately in every seed. Geometric means of class ratios consequently obey the same product identity. In compensated mixtures, the soft-to-normal ratio of *S*_*v*_ declines from approximately 1.18 at 5% to 1.01 at 50%, whereas the *T*_*v*_ ratio remains approximately 0.81–0.84 (Fig. S6). At 5% and 10%, the *D*_8_ class difference is unresolved. At 15–50%, soft-cell *D*_8_ is smaller. At the higher fractions, the shorter persistence of soft-cell motion outweighs its higher mean-square speed.

For modified cells in the control with *M*_*i*_ reduced in 30% of cells, 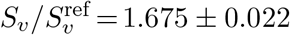 and 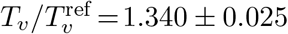 at 8*τ*. For compensated soft cells relative to all-normal controls, the ratios are 1.315 ± 0.016 and 0.891 [0.867, 0.916]. Under matched propulsion histories, reducing *M*_*i*_ increases mean-square speed and persistence. Compensation increases mean-square speed but decreases persistence. Compensated soft cells retain a higher *D*_8_ than all-normal cells but have a lower *D*_8_ than normal cells in the same mixture.

To test whether a change in correlation amplitude alone explains the transport response, we compare the control with reduced *M*_*i*_ and the compensated mixtures with their paired all-normal simulations. We analyze modified and unmodified cells separately in the control with reduced *M*_*i*_, and soft and normal cells separately in the compensated mixture. For each comparison, the reference correlation is calculated from cells with the same labels in the paired all-normal simulation. For each seed, we multiply this reference correlation by 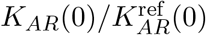, where *K*_*AR*_(0) is the propulsion–relaxation correlation at zero time separation in the mixture and 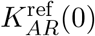 is the same quantity in its all-normal reference. This changes the amplitude of the reference correlation while retaining its time dependence, giving the prediction 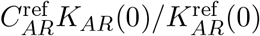. For unmodified cells in the control with *M*_*i*_ reduced in 30% of cells, the predicted change is − 0.00351 ± 0.00248, whereas the measured change is +0.00813 ± 0.00337 (Fig. S7). Since the velocities are recorded at discrete times, we checked whether the choice of numerical integration rule affects this comparison. Trapezoidal, left-endpoint and right-endpoint integration all give a negative predicted change and a positive measured change. The increased cross term therefore requires a change in temporal correlations beyond equal-time amplitude.

#### S5.2 Relaxation after reorientation and contributions within polarity intervals

Projecting the relaxation velocity onto the new and preceding propulsion directions shows how its alignment changes after a turn. This analysis covers all the simulation conditions listed above. We identify intervals of constant polarity from each cell’s recorded direction. Recording intervals containing a direction change are treated separately. The time origin is the first frame recording the new direction. For each subsequent interval between recordings, *a* is the time from the first recording of the new direction to the interval midpoint. We project the mean relaxation velocity over that interval onto the new direction 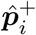 or the preceding direction 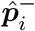. The plotted conditional means are 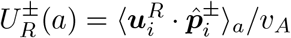. We include only intervals before the next reorientation.

Age bins have width 0.0125*τ* up to 0.1*τ*, followed by boundaries at 0.15, 0.2, 0.3, 0.5, 0.75, 1, 1.5, 2, 3, 4*τ*. Within each time bin, we first average each cell’s response, weighting intervals by their duration. We then average equally over the contributing cells within each class. We also average responses over 0.025–0.1, 0.2–0.5, 1–2 and 2–4*τ*. Seed means and pointwise 95% intervals are shown in Fig. S3. The restriction to cells that have not reoriented again applies only to these response curves. The transport estimates include all cells.

To quantify how these responses contribute to transport, we separate the propulsion–relaxation displacement product into contributions within and between intervals of constant polarity. For observation window *w*, let *T*_*irw*_ and 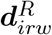 denote the duration and relaxation displacement of the portion of resolved polarity interval *r* of cell *i* lying inside the window. Propulsion has a constant direction over this portion, giving

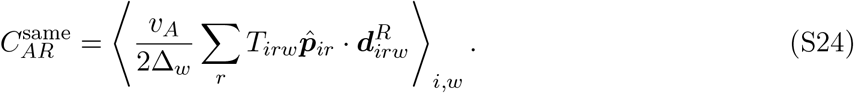

We calculate these products directly, weighting each polarity interval by its duration inside the observation window. The conditional response curves average at fixed age and do not carry these observation-window weights.

Every remaining pair is assigned to one of three contributions: 1) earlier propulsion with relaxation in a subsequent polarity interval, 2) later propulsion with relaxation in a preceding polarity interval, or 3) a pair containing at least one recorded interval in which the cell reorients. Thus

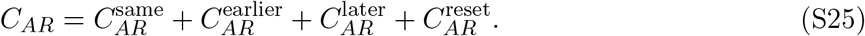

The earlier and later labels specify the propulsion interval relative to the relaxation interval and include separations spanning multiple reorientations.

Within each term of Eq. S25, we group relaxation ages into 0–0.1, 0.1–0.5, 0.5–1, 1–2, 2–4 and ≥ 4*τ*. The initial interval with unknown age and intervals containing reorientations are retained separately, so all groups sum to the original *C*_*AR*_. Figure S8 shows projection onto the preceding polarity. Figure S9 shows the fraction and class comparisons.

In the binary population comparison between 5% and 50%, 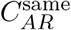 increases by 0.04641 ± 0.00164, compared with the full *C*_*AR*_ change of 0.04866. The contributions from earlier propulsion, later propulsion, and intervals containing a reorientation change by 0.00204, − 0.00042, and 0.00064, respectively. Relaxation intervals at ages *a* ≥ 0.5*τ* contribute 0.03583 of the same-interval change. Most of the increase in the same-interval term therefore comes from these later relaxation increments. Over 1–2*τ*, the normal-cell projection on the new direction changes by (0.05051 ± 0.00453)*v*_*A*_ between these fractions. The soft-cell change is (0.04467 ± 0.01036)*v*_*A*_.

For unmodified cells in the control with reduced *M*_*i*_, the early (0.025–0.1*τ*) projection on the new direction changes by (−0.03782 ± 0.00315)*v*_*A*_ relative to the all-normal reference, whereas the projection on the preceding direction changes by (0.03721 ± 0.00223)*v*_*A*_. The same-interval cross-term change is unresolved, 0.00127 ± 0.00301, while correlations with earlier propulsion contribute 0.00717 ± 0.00175. Together with the remaining contributions, these give the full increase 0.00813 ± 0.00337. Most of the increase in *C*_*AR*_ therefore comes from correlations with propulsion before the most recent reorientation.

Modified cells in the control with reduced *M*_*i*_ instead have a same-interval change of 0.19744± 0.00377 relative to matching all-normal labels. For compensated soft cells, the same-interval change is negative at every fraction, ranging in mean from −0.08761 to −0.05235. Homogeneous softening increases the population same-interval cross term at every fraction. Relative to the matched homogeneous mean-*γ* condition, half contrast also increases that contribution in the lower-*γ* class at every fraction. Its population change is unresolved at 5–30%. The 50% point is small and positive, 0.00248 ± 0.00237 (Fig. S9g–l).

#### S5.3 Displacement contributions in the control comparisons

The coefficient ratios for these controls are given in Sec. S3.1. Compensation preserves the binary free-energy functional while changing its kinetics. The control with reduced *M*_*i*_ preserves the all-normal free-energy functional.

##### Calculation of the displacement contributions

For the control comparisons, relaxation displacement is estimated from centroid displacement and integrated advection, using the actual elapsed time of each window (Sec. S2.3). Homogeneous controls, compensated mixtures and the control with reduced *M*_*i*_ are compared with matched all-normal references. Half-contrast mixtures are compared with the homogeneous control at the same mean coefficient. Both compensated mixtures and the control with reduced *M*_*i*_ have *δD*_*A*_ = 0. Recorded propulsion differences are retained in the homogeneous comparison.

##### Homogeneous softening

Homogeneous softening at 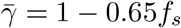 gives *δC*_*AR*_ *>* 0, *δD*_*R*_ *<* 0 and *δD*_8_ *>* 0 relative to all-normal controls throughout the series. The cross-term increase exceeds the decrease in *D*_*R*_, and each change has a pointwise interval excluding zero. The binary mixtures show the same balance between these two terms.

##### Compensated mixtures

Compensated soft cells have larger mean *D*_*R*_ and more negative mean *C*_*AR*_ than matching all-normal cells at every fraction. Normal cells have the opposite changes. Both classes retain positive mean transport gains. At 30%, the geometric transport ratios are approximately 1.17 for soft cells and 1.29 for normal cells. Thus compensation reverses the within-mixture class ordering while both classes remain faster than their all-normal references. Figure S10 compares these signed contributions at Δ = 8*τ*. Here the remaining terms are *δ*(*D*_*A*_ + *D*_*I*_ + *C*_*AI*_ + *C*_*IR*_). Cell labels in each reference match those in its paired system.

##### Temporal correlations after reducing *M*_*i*_

For unmodified cells in the control with *M*_*i*_ reduced in 30% of cells, *D*_*R*_ and *C*_*AR*_ both increase relative to all-normal controls. Equal-time amplitude rescaling predicts the wrong sign for the cross-term change (Sec. S5.1). The polarity-interval partition in Sec. S5.2 locates most of that increase in correlations with earlier propulsion.

### S6 Fixed-mean controls and geometric-mean calibration

The homogeneous and half-contrast systems distinguish a change in the mean interfacial coefficient from a distribution of coefficients about the same mean. Full-contrast mixtures have *γ*_n_ = 1 and *γ*_s_ = 0.35, giving 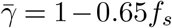. Half contrast replaces each coefficient by 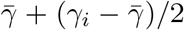, while homogeneous populations have 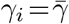. The conditions therefore share the same mean at each fraction.

#### Matched transport balance across the fraction series

Figure S11 compares the three conditions across the fraction series with 10 matched seeds at each fraction and corresponding cell indices, as in main Fig. **??**c. The three conditions use matching propulsion histories. We integrate propulsion and interaction-advection velocities at each simulation’s recorded time points, then select displacement origins and endpoints shared by all three conditions within 0.01–50*τ* of activity. Total relaxation is calculated from these integrals and centroid displacement using the estimator described above. The primary lag is 8*τ*, and both mixtures are compared with the same homogeneous reference. Here *δ* is the difference between mixture and the homogeneous reference.

At 8*τ*, the mean full-contrast population transport exceeds the homogeneous reference at every fraction. Five of the six pointwise 95% intervals exclude zero. The interval at 10% includes zero. The mean increase in *C*_*AR*_ exceeds the mean decrease in *D*_*R*_ throughout. The *C*_*AR*_ increase is resolved at every fraction. The *D*_*R*_ decrease is unresolved at 5% and 15%. Confidence intervals are calculated from differences formed within each seed. The remaining terms oppose the gain by at most 6.7% of its mean. The mean transport excess stays positive at 4*τ* and 16*τ*. The mean half-contrast excess is smaller at every fraction. For the common-window comparison in Fig. S11, its interval includes zero throughout the series. The direct full-minus-half transport difference is resolved at five fractions with the 10% interval including zero. The homogeneous and half-contrast recordings provide total relaxation without separating its interfacial and overlap contributions.

#### Effective coefficient and geometric-mean calibration

We define *γ*_eff_ as the interfacial coefficient of a homogeneous monolayer with the same population-mean transport as the mixture. We compare this coefficient with the arithmetic, geometric and harmonic means of the interfacial coefficients. We use population-mean centroid transport *D*_8_ from matched seed recordings. Each recording averages all 400 cells with equal weight over the common complete 8*τ* displacement windows within 0.01–50*τ* of activity. Recording means are then averaged arithmetically across seeds. The homogeneous coefficients are 0.9675, 0.9350, 0.9025, 0.8700, 0.8050, 0.6750 and 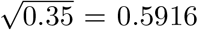.

The last coefficient is the geometric mean of the 50% full-contrast mixture. Their ensemble-mean transports are monotonic. We interpolate these means linearly in *γ* to obtain 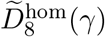 and solve 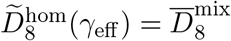.

For the full-contrast mixtures, the geometric and harmonic means are 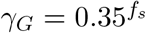 and *γ*_*H*_ = (1 *f*_*s*_ + *f*_*s*_*/*0.35)^−1^, respectively. For half contrast, the geometric mean uses the coefficients 1 − 0.325*f*_*s*_ and 0.675 − 0.325*f*_*s*_ with weights 1 − *f*_*s*_ and *f*_*s*_. At *f*_*s*_ = 0.05, 0.10, 0.15, 0.20, 0.30 and 0.50, respectively, the full-contrast estimates of *γ*_eff_ are 0.9488, 0.8956, 0.8594, 0.7933, 0.7242 and 0.5972. The corresponding geometric means are 0.9489, 0.9003, 0.8543, 0.8106, 0.7298 and 0.5916.

The maximum absolute relative difference,

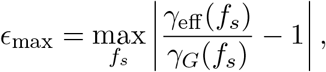

is 2.13%, attained at *f*_*s*_ = 0.20. All six full-contrast point estimates lie between *γ*_*H*_ and 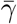. All six full-contrast inverse estimates lie within the sampled homogeneous range. Of the half-contrast inverse estimates, only the 5% value requires extrapolation. As a sensitivity check, interpolation linear in ln *γ* gives a maximum full-contrast difference of 2.25%. Monotone piecewise-cubic interpolation in *γ* gives 2.29% across all six fractions.

We estimate the uncertainty in the transport difference between a mixture and the homogeneous calibration at *γ*_*G*_. If *γ*_*k*_ and *γ*_*k*+1_ bracket *γ*_*G*_, let *w* = (*γ*_*G*_ − *γ*_*k*_)*/*(*γ*_*k*+1_ − *γ*_*k*_). Within seed *j*, define

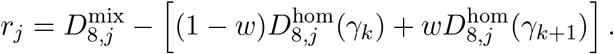

At 50% full contrast, *r*_*j*_ subtracts the directly measured homogeneous transport at 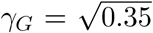. The other geometric-mean coefficients lie inside the measured calibration range and use linear interpolation. Pointwise 95% Student-*t* intervals use the paired residuals and the corresponding number of seeds. All twelve intervals include zero (Fig. S12c). They include finite-seed variation in both mixture and calibration. Interpolation error is not estimated. The same seeds are used across fractions, so estimates at different fractions are correlated. Neither contrast level shows a resolved transport difference from its geometric-mean reference.

Figure S13 compares the cell-class transport ratio in the half-contrast mixture with that of the corresponding labels in the homogeneous reference:

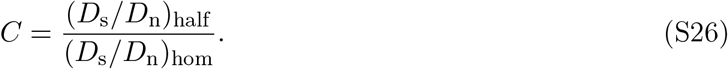

In the half-contrast mixture, s and n label the lower- and higher-*γ* classes. Both coefficients vary with fraction. In the homogeneous control, they label the corresponding cell indices. This reference removes differences between the designated groups that are already present without material contrast.

#### Weak-contrast expansion

Averaging Eq. **??** over cells and using ∑_*i*_ *χ*_*ij*_ = *c*, the term linear in the contrast is

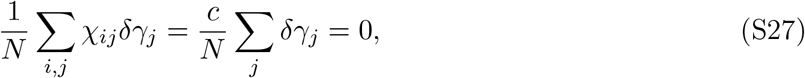

where the last equality follows from *∑* _*j*_ *δγ*_*j*_ = 0 at fixed 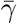.

Population transport in the homogeneous reference is unchanged by relabeling cells after averaging over initial configurations and polarity histories. We can therefore write its second-order response with two coefficients, *a* and *b*:

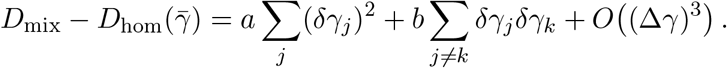

Using ∑_*j*_*δγ*_*j*_ = 0 gives ∑_*j≠k*_*δγ*_*j*_*δγ*_*k*_ = −∑_*j*_ (*δγ*_*j*_)^2^, and hence

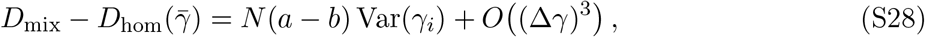

where Var(*γ*_*i*_) = *N* ^−1^ ∑_*i*_(*δγ*_*i*_)^2^ = *f*_*s*_(1 − *f*_*s*_)(Δ*γ*)^2^ for a binary mixture.

The single-soft-cell limit differs from fixed-mean weak contrast. At fixed finite contrast and *f*_*s*_ = 1*/N* → 0, a finite change in the tagged cell’s transport contributes only *O*(*N* ^−1^) to the population mean. The same scaling holds if only finitely many neighbors have finite responses. A non-vanishing population change instead requires the sum of surrounding-cell responses to grow with *N*.

### S7 Interfacial relaxation and contact exchange

Direct recordings of interfacial and overlap relaxation allow us to examine how softening one cell population affects the motion of neighboring normal cells. We use paired *N* = 400 binary and all-normal simulations. The fraction series is *f*_*s*_ = 0.05, 0.10, 0.15, 0.20, 0.30 and 0.50. Each pair shares its fields before passive preparation and its imposed polarity history. Ten paired seeds are used for the displacement balances, responses to cell reorientation and field-equation evaluations at saved configurations. Five pairs per fraction are used for the accepted T1 measurements. Cells and time origins are averaged within each seed before forming paired differences. Population averages give equal weight to every cell. We then average equally over seeds. Unless stated otherwise, *δX* = *X*^binary^ − *X*^0^, Δ = 8*τ*, and displacement contributions are reported in units of 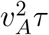.

**Binary-mixture displacement balance.** For complete 8*τ* windows within 8–50*τ* of activity, the mean increase in *C*_*AR*_ exceeds the mean decrease in *D*_*R*_ in both cell classes. Both *C*_*AG*_ and *C*_*AK*_ contribute to the increase. Population changes are shown in main Fig. **??**a. With matched propulsion histories, *δD*_*A*_ = 0. The remaining interaction-advection terms are retained, so that 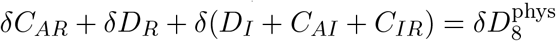 within each seed. Numerical differences from centroid transport are assessed in Sec. S2.3.

#### S7.1 Interfacial and overlap relaxation

The phase-field equation is *∂*_*t*_*ϕ*_*i*_ + ***v***_*i*_· ∇ *ϕ*_*i*_ = −*M*_*i*_ *δℱ /δϕ*_*i*_. The interfacial (*G*) and overlap (*K*) contributions to field relaxation are

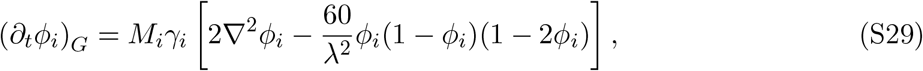

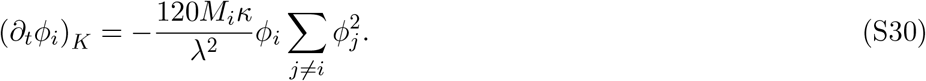

At fixed fields and *M*_*i*_, lowering *γ*_*i*_ reduces the interfacial source while leaving the overlap source unchanged. The resulting field evolution changes both sources.

For 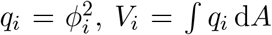 and *S*_*i*_ = ∑_*j ≠ I*_ *q*_*j*_, Eq. S17 defines 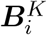 and the associated overlap contribution 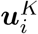 to centroid velocity.

#### S7.2 Neighbor exchanges and displacement correlations

To relate the transport changes to local cell rearrangements, we measure centroid motion and contact-length changes during four-cell neighbor exchanges (T1 events). Cell boundaries are defined by the *ϕ*_*i*_ = 0.5 contours. For each pair of cells, we measure the length of each contour lying within seven grid units of the other, then average the two lengths. Contact-on and contact-off thresholds are three and 1.5 grid units, respectively. These boundary contacts are distinct from the centroid Delaunay neighbors used in Sec. S9.

An accepted four-cell T1 exchange loses one diagonal contact and forms the other while preserving the four-cell ring. The old diagonal and ring must persist for 0.1*τ* before contact loss, the ring must remain intact during exchange, and the new diagonal and ring must persist for 0.1*τ* after completion. Events include participants of either cell type. Each event receives equal weight within its seed. Seed means are the independent observations. Accepted events need not be identical in the two systems of a pair. Five paired seeds are used at every fraction.

##### Relative advance during an exchange

Let *d*_old_ and *d*_new_ be the centroid separations of the cell pair losing contact and the pair gaining contact, respectively, and define the relative advance *q* = (*d*_old_ − *d*_new_)*/*(*v*_*A*_*τ*). This quantity increases as the old neighbors separate or the new neighbors approach. In the final 0.1*τ* of intact old contact, its dimensionless rate *u*_*q*_ = d*q/*d(*t/τ*) increases from 0.878 in all-normal controls to 0.998 in 50% binary mixtures.

The increase persists after grouping events by *q* immediately before contact loss and by the propulsion contribution to relative advance over the pre-loss interval. Bin boundaries are set by the pooled all-normal controls: five equal-population bins for *q* and either five or 10 for propulsion. Within each seed pair, we retain bins with at least 10 events in each condition and average both conditions with weights proportional to their control event counts.

During completion, the rate is instead 1.409 in controls and 1.372 in mixtures with smaller integrated advance in mixtures. Four-normal-cell exchanges at 30% also show less opposing interfacial motion before loss. Soft cells may lie outside the quartet. We repeat the comparison after excluding events with participant reorientations and matching polarity ages. The additional relative centroid advance precedes contact loss.

##### Contact shrinkage, growth and exchange duration

The contact-length measurement follows the old boundary as it shortens and the new boundary after it forms. Junction lengths and the interval between contact loss and formation are distinct observables in phase-field descriptions of T1s [5]. Here *L* denotes the contact length defined above. Let *t*_*o*_ be the last saved time with the old contact present and *t*_*n*_ the first with the new contact present. With *h* = 0.1*τ*, the mean shortening and growth speeds are

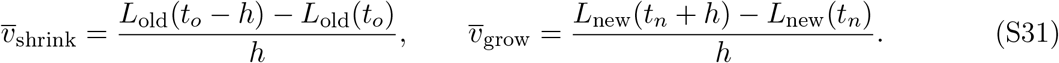

The recorded exchange interval is *T* = *t*_*n*_ − *t*_*o*_. Each contact-change time is bracketed by frames separated by 0.01*τ*. The same accepted events enter the centroid and boundary comparisons. The old contact is present throughout the shortening interval, and the new contact throughout the growth interval.

At *f*_*s*_ = 0.50, old-contact shortening is (2.0 ±0.5)% faster and new-contact growth is (4.2± 0.9)% faster than in all-normal controls. The mean exchange interval decreases from 0.307*τ* to 0.283*τ*, a paired reduction of (7.8 ±1.9)% (Fig. S14). These uncertainties denote pointwise 95% paired-seed confidence intervals. Percentages are averaged over seed-pair ratios. Faster new-contact growth and the shorter exchange interval persist when participant reorientations are excluded.

During exchange at 50%, mean relative centroid advance Δ*q* decreases from 0.368 to 0.332 and mean speed from 1.409 to 1.372. For each event, the mean rate and duration give the relative advance through Δ*q* = *ū*_*q*_*T/τ*. Contact length also depends on cell deformation, so centroid speed and contact-growth speed are distinct measurements. Figure S14 shows the kinematic comparisons.

#### S7.3 Relation to repulsive-particle force imbalance

**Identical circular profiles**. For identical circular cell profiles, overlap-induced relaxation can be written as a sum of pairwise repulsive velocities. Consider instantaneous fields *q*_*i*_(***x***) = *q*_0_(|***x*** − ***r***_*i*_|) with identical, nonnegative circular profiles, common *M*, and common *V = ∫q*_0_ d*A*. The profiles are localized, and the integrals extend over the plane. Define ***r***_*ij*_ = ***r***_*j*_ − ***r***_*i*_, *r*_*ij*_ = |***r***_*ij*_| and

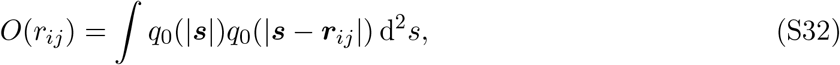

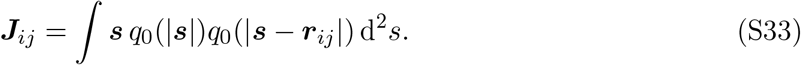

The change of variables ***s***↦***r***_*ij*_ −***s*** leaves the product of the two profiles unchanged. Consequently ***J***_*ij*_ = ***r***_*ij*_*O*(*r*_*ij*_) −***J***_*ij*_: the overlap centroid is exactly at the pair midpoint whenever the overlap is nonzero. Equation S17 becomes

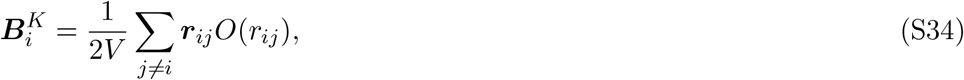

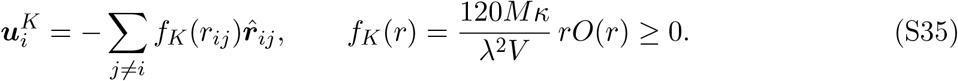

At zero separation the vector contribution is zero. For sufficiently localized profiles, the kernel also admits a repulsive potential representation, 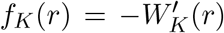 with 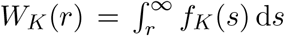.

Here *f*_*K*_ has units of velocity: it represents centroid drift generated by field relaxation. It is distinct from the model’s interaction-advection contribution *I*. Translating the overlap energy gives a derivative of *O*(*r*), whereas this relaxation drift is proportional to *rO*(*r*). Both contributions enter the displacement balance separately. In the binary mixture, interfacial relaxation changes the evolving profiles and areas. Equation S17 gives the overlap velocity for these general fields.

##### Angular average and force imbalance

In a spatially uniform ensemble, write *n* for cell number density and define *g*(*r, θ*) so that the mean number of neighbors in the element *r* d*r* d*θ* is *ng*(*r, θ*)*r* d*r* d*θ*, in a frame aligned with the tagged cell’s propulsion. The angle *θ* = 0 points forwards.

Projecting Eq. S35 and averaging gives

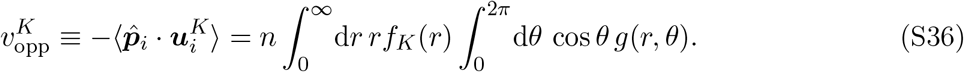

Bialké *et al*. [4] obtain the opposing drift *nζ* with exactly this angular integral and radial weight *r*[−*u*^′^(*r*)], where *u* is the repulsive pair potential and translational mobility is absorbed into their units. Replacing −*u*^′^(*r*) by the derived velocity kernel *f*_*K*_(*r*) gives Eq. S36. An isotropic neighbor distribution has zero cosine integral. A positive force-weighted excess ahead gives positive opposition. The integral selects the first angular cosine component of the interaction distribution, weighted by the radial velocity kernel.

#### S8 Numerical implementation and simulation conditions

Unless stated otherwise, the simulations use (*κ, ξ*) = (10, 1500) and the spatial and temporal discretization of Palmieri *et al*. [2]. The square periodic lattice has unit spacing and integer side 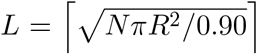, where *N* is the number of cells, *R* is the target cell radius, and 0.90 is the target value of the nominal area fraction *NπR*^2^*/L*^2^. The ceiling brackets round up to the smallest integer not less than their argument. Time integration and stencils are as stated in Methods. The interface-width parameter is *λ* = 7 lattice units and the cell radius is *R* = 7*λ*.

The reorientation rule is defined in Sec. **??**. We evaluate its probability in double precision. The simulations with reduced *M*_*i*_ at both *ξ* = 1500 and 375 use *p*_t_ = −expm1(−*δt/τ*), where expm1(*x*) = *e*^*x*^ −1 evaluates the exponential difference accurately for small arguments. The paired simulations use matching polarity histories (Sec. S3.2). At each timestep, each cell’s polarity is updated according to the reorientation rule above. The interaction-induced velocity and field-relaxation terms are calculated from the fields at the beginning of the step with the overlap contribution obtained by subtracting 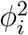 from the aggregate field 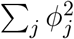. Advection and relaxation are combined in a single forward-Euler update. The updated fields are stored separately, and their areas, centroids, support bounds and aggregate overlap field are calculated for the next step. CUDA thread blocks process different cells concurrently. Within each block, threads share the grid-point calculations for one cell.

Each phase field is stored in single precision on a 288 ×288 tile and advanced on a moving local domain. A similar moving-subdomain construction is described by Chiang *et al*. [6]. Cell-based parallelization on adaptively refined full-domain meshes is described by Praetorius and Voigt [7]. The domain is recentered using the 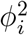-weighted centroid and enlarged according to the bounding box of points for which *ϕ*_*i*_ *>* 10^−5^, up to an active domain of 286 ×286 sites. For a locally planar equilibrium interface, the exterior tail a distance *d* from the *ϕ*_*i*_ = 1*/*2 contour is

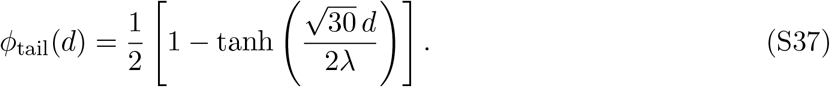

At *λ* = 7, the support threshold lies 14.7 lattice sites outside the half-height contour.

To generate initial configurations, trial cell centers are drawn uniformly in the periodic box and added sequentially. A trial center is accepted when its minimum-image distance from every previously accepted center exceeds an exclusion distance, initially 2*R*. After 10,000 unsuccessful trials, this distance is reduced by 5%, but never below *R*, allowing the target number of cells to be placed at the prescribed density. Each accepted center then initializes a circular diffuse phase field using the periodic profile in Eq. S38.

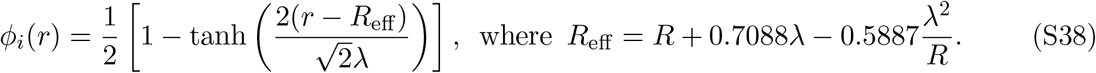

For *R* = 49 and *λ* = 7, Eq. S38 gives *R*_eff_ = 53.3729 and 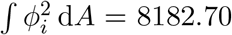0, compared with the target *A*_0_ = *πR*^2^ = 7542.96. Before passive relaxation, the circular profile therefore has a 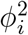-weighted area 8.48% above the target.

The systems in the finite-size series are relaxed passively for 8*τ* before activity is enabled. The composition simulations begin from equilibrated all-normal *N* = 400 configurations and are relaxed for a further 1*τ* after the soft labels are assigned. The finite-size transport estimates use a 200*τ* active window, and the composition estimates use a 50*τ* active window. Table S4 lists the simulation sets. The trajectories used for the spatial estimates are specified in Sec. S9. We calculate each displacement by summing the shortest periodic displacements between successive recorded positions. Restarted trajectory segments are joined chronologically, replacing an earlier copy of any overlapping stored time.

#### S8.1 Simulation conditions and paired controls

The single-cell limit uses paired *N* = 400 systems at *f*_s_ = 0 and 1*/*400, prepared with the same binary/all-normal protocol. After 8*τ* of all-normal passive equilibration, one predetermined cell is assigned *γ* = 0.35 in the binary mixture and retains *γ* = 1 in the all-normal control. Both systems then relax passively for 1*τ* and evolve actively for 50*τ* with matched polarity histories. The tagged-cell *D*_8_ ratio is 1.187 [0.873, 1.613], and the ratio for the other cells is 0.997 [0.972, 1.021].

The same binary/all-normal protocol is also evaluated at *N* = 400 with soft-cell proportions from 0.5% to 4% to resolve the low-fraction crossover. Each cell type’s *D*_8_ is normalized by the value for the same cell labels in its all-normal control. The same all-normal reference simulations are used at every soft-cell fraction, and comparisons between fractions are paired by seed.

Normal-cell *D*_8_ increases over this interval (main Fig. **??**a). At *f*_s_ = 0.04, its ratio to the all-normal control is 1.046 ±0.020, while the population ratio is 1.050 [1.028, 1.071]. The positive within-seed trend is retained at lags of 4*τ* and 16*τ*.

##### S8.1.1 Controls at fixed mean interfacial coefficient

The half-contrast controls retain the original fraction and population mean 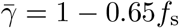, but use

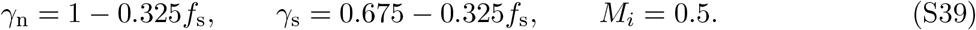

The half-contrast mixtures and homogeneous controls share the fields before their 1*τ* passive preparation, designated cell labels and imposed polarity histories. Their fields at the start of active evolution can differ because passive preparation uses the respective material coefficients. Each active window spans 50*τ*. The compensated mixtures and the control with reduced *M*_*i*_ are defined in Sec. **??** and Table **??**. Their all-normal references have *M*_*i*_ = 0.5 and *γ*_*i*_ = 1 for every cell and share designated cell identities and recorded polarity histories with the paired condition within each seed.

#### S8.2 Numerical validation

We compared the CUDA update with an independent double-precision implementation on the complete periodic lattice. The test states were an *N* = 12 monolayer containing one deformed soft cell and two elongated variants of that configuration. They tested the stencil, interaction and area terms, including the largest moving domain. Activity and reorientations were disabled for the one-step comparison. The relative *L*_2_ difference between CUDA and double-precision updates is 1.9 × 10^−8^–4.8 × 10^−8^ (Table S5a). Rounding the overlap density to the solver’s fixed-point representation changes the update by at most 1.2 × 10^−9^.

With fixed polarity, we evolved the deformed state for 0.1*τ* at timesteps 0.01, 0.005 and 0.0025. Relative to the finest calculation, the production timestep gives a field difference of 1.26 × 10^−4^ in relative *L*_2_ norm and a centroid RMS difference of 3.47 ×10^−5^*R* (Table S5b). The soft-cell and population RMS displacement differences are 0.033% and 0.009%.

With propulsion disabled, the free energy decreased at every recorded step in the validation runs. Restarting from a saved checkpoint reproduced the same final fields as uninterrupted evolution.

For the simulations at *ξ* = 375, we independently recalculate centroid transport from unwrapped trajectories using Fourier correlations of displacements. The calculation covers the three populations, lags 4, 8, 16*τ* and both observation windows (0.01–50*τ* and 8–50*τ*). All estimates agree to an absolute difference below 1.4 ×10^−10^. We also checked the time integrals of the recorded velocity contributions against recordings taken at shorter intervals. The relative net-*R* discrepancy was below 1.2 ×10^−6^ over that 0.1*τ* test.

Errors in recorded positions can affect MSD-derived diffusivities [8]. We assessed their effect on the tagged-cell transport ratio by assuming a maximum error of one lattice spacing in each coordinate at every recorded time. The resulting worst-case bounds on the pooled soft/control *D*_8_ ratio are 1.032 and 1.252, compared with the measured ratio of 1.137. The lower bound remains above unity, so position errors of this size cannot fully account for the observed enhancement.

### S9 Spatial dependence of transport

#### S9.1 Single-soft-cell systems

To test whether a single soft cell changes the transport of nearby normal cells, we measure their diffusivity as a function of distance from the tagged cell. For each interval in Eq. S1, the minimumimage distance from normal cell *i* to the tagged cell at the start of the interval is *r*_*i*_(*t*) = |**r**_*i*_(*t*) − **r**_0_(*t*)| _min_. Within each simulation, we average *Q*_*i*_ over all cell observations whose interval-start distance falls in a bin *B*, obtaining *D*(*B*). The plotted factor is

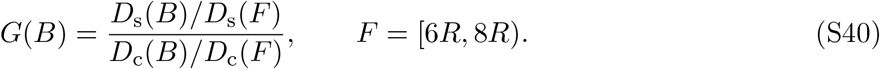

Here the subscripts denote the single-soft-cell simulation and its all-normal control. The tagged cell is excluded from both measurements. Thus each profile is normalized by its own far-region value before division by the control profile. The control accounts for spatial variation also present around a normal tagged cell. Thirteen bins of width 0.5*R* cover 1.5*R* ≤ *r <* 8*R*. The prespecified near/far comparison replaces *B* by [1.5*R*, 3*R*).

The spatial comparison uses 200*τ* active windows with 10 paired seeds at each of *N* = 400, 800 and 1600, three at *N* = 3200, and two at *N* = 6400. Restricting it to *N* ≥ 400 keeps the reference radius 8*R* below one quarter of the box side. The pooled profile gives equal weight to *N* = 400, 800 and 1600: one replicate from each size is averaged within each of 10 predefined groups. Larger systems are shown individually.

##### S9.1.1 Choice of distance bands

We assess the near band against the first-neighbor shell, whose outer extent can be identified from the first minimum of the radial distribution function *g*(*r*) [9]. At the displacement origins used above, we calculate *g*(*r*) from the mean number of normal-cell centroids in each annulus, divided by its area and the mean density (*N*− 1)*/L*^2^. In both the single-soft-cell and all-normal systems, the first peak lies in 1.5*R*≤ *r <* 2*R* and the first minimum in 2.5*R* ≤*r <* 3*R* (Fig. S15a). The near band therefore covers the principal first-neighbor peak. The far band samples separations of three to four nominal cell diameters.

We repeat Eq. S40 for all 30 combinations of the five near bands and six far bands shown in

Fig. S15b,c. Regional diffusivities are recalculated from the 0.5*R* bins, weighted by their observation counts. The lag, paired comparisons and averaging over sizes and replicate groups are unchanged. The pooled factors range from 1.004 to 1.020, compared with 1.012 for the original bands. Every pointwise 95% interval includes unity. Thus no preferential increase in nearby normal-cell transport is resolved for any of these choices.

#### S9.2 Finite soft-cell fractions

At finite soft-cell fractions, we ask whether the increase in normal-cell transport is concentrated near soft cells. For *N* = 400 binary mixtures at *f*_s_ = 0.05, 0.10, 0.15, 0.20, 0.30 and 0.50, we measure each normal cell’s distance to the nearest soft cell at the start of an 8*τ* displacement window. Window starts occur at *t/τ* = 1, 3, …, 41, relative to the first stored active frame. We normalize by the mean displacement contribution of normal cells at the same start time,

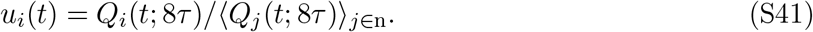

Immediate neighbors are defined by a periodic centroid Delaunay triangulation. We triangulate a 3 ×3 tiling of the centroid positions (using the Rust delaunator library, version 1.1.0), retain edges associated with the central box, and remove duplicate edges between cell identities. Each sampled graph has 3*N* distinct edges. Let *A* be the mean *u*_*i*_ for normal cells with at least one soft neighbor divided by the mean for normal cells without one. All cell observations are pooled within their respective groups in each simulation.

To remove geometrical associations unrelated to cell type, we repeat the classification using 32 distinct uniformly selected sets of exactly *Nf*_s_ reference labels. Each set is fixed throughout a trajectory. The measured cells and the normalization in Eq. S41 remain the actual normal cells. A cell is never its own reference neighbor. The corrected neighbor factor is

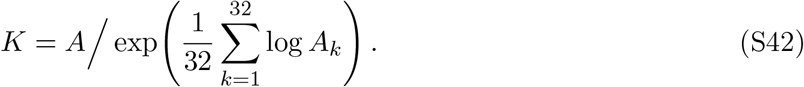

For the distance-resolved finite-fraction profile, the mean *u*_*i*_ in each distance bin is divided by the arithmetic mean of the 32 reference-label bin means. A bin is retained only if it contains at least 1% of the normal-cell observations in every replicate and every reference-label set at that fraction. No difference from the random-label reference is resolved across the retained bins and six fractions (simultaneous global *p* = 0.307).

#### S9.3 Cell-type mixing

To test whether the composition-dependent *D*_8_ response is accompanied by cell sorting, we construct the periodic centroid Delaunay graph at *t/τ* = 1, 3, …, 47. If the graph contains *M* edges, *n*_s_ soft cells, and *M*_SS_ soft–soft edges, the soft–soft enrichment is

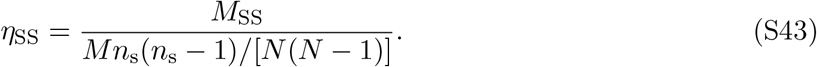

Uniform reassignment of exactly *n*_s_ labels on the same graph gives⟨*η*_SS_⟩ = 1. We average the enrichment over the sampled times within each simulation, then calculate the mean and confidence interval across the 10 independent simulations. No fraction shows a resolved positive enrichment or increase in enrichment over the 50*τ* production window (main Fig. **??**b). At *f*_s_ = 0.50, the time-averaged enrichment is 0.980 ± 0.011 (95% confidence interval).

#### S10 Statistical inference

Interval, replicate and geometric-mean conventions are given in Methods. Here we describe the sign-flip test and the simultaneous confidence bands.

We use a sign-flip test to determine whether transport differs from the reference in any distance bin or at any soft-cell fraction. For each replicate, the test applies the same sign, +1 or −1, to the effects measured in all distance bins or at all fractions. For the ten-replicate comparison sets, we enumerate all 2^10^ sign assignments and compare the observed maximum absolute *t* statistic across the displayed comparisons with this null distribution. The test assumes that replicate-level effects are symmetric under the null.

For simultaneous confidence bands, we first subtract each comparison’s mean from its replicate values. Under each sign assignment, we divide the resulting mean by its standard error and take the largest absolute value across comparisons. The 95th percentile of these maxima sets the common multiplier of the original standard errors. Mixing enrichment uses pointwise intervals. Sections S2 and S7 use pointwise Student intervals across paired seeds.

**Table S1:** Displacement balance at lag 8*τ* in the all-normal reference. Displacement windows lie within 8–50*τ* after activity begins. Values are means± half-widths of 95% Student-*t* confidence intervals across 10 independent simulations, in units of 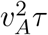.

| Contribution | Value |
|---|---|
| *D*<sub>*A*</sub> | 0.43943 ± 0.00506 |
| *D*<sub>*R*</sub> | 0.31976 ± 0.00480 |
| *C*<sub>*AR*</sub> | −0.71584 ± 0.00996 |
| *D*<sub>*I*</sub> | 0.0000935 ± 0.0000014 |
| *C*<sub>*AI*</sub> | −0.01216 ± 0.00017 |
| *C*<sub>*IR*</sub> | 0.01079 ± 0.00016 |
| *D*<sub>*I*</sub> + *C*<sub>*AI*</sub> + *C*<sub>*IR*</sub> | −0.00127 ± 0.00003 |
| $D_8^{\text{phys}} = D_A + D_R + C_{AR} + D_I + C_{AI} + C_{IR}$ | 0.04208 ± 0.00092 |

**Table S2:** Passive field coefficients relative to a normal cell. The interfacial column is *M*_*i*_*γ*_*i*_*/*(0.5 ×1), and the overlap/area column is *M*_*i*_*/*0.5. The advection coefficient is unchanged in these comparisons.

| Cell condition | *γ<sub>i</sub>* | *M<sub>i</sub>* | Interfacial | Overlap/area |
|---|---|---|---|---|
| Normal and unmodified | 1 | 0.5 | 1 | 1 |
| Binary mixture, soft | 0.35 | 0.5 | 0.35 | 1 |
| Compensated soft | 0.35 | 0.5/0.35 | 1 | 1/0.35 |
| Modified (*M<sub>i</sub>* reduced) | 1 | 0.175 | 0.35 | 0.35 |

**Table S3:** Population displacement balance at *ξ* = 375 and lag Δ = 8*τ* in units of 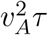.

| Contribution | Reduced *M*<sub>*i*</sub> | All-normal reference |
|---|---|---|
| *D*<sub>*A*</sub> | 0.43989 | 0.43989 |
| *D*<sub>*R*</sub> | 0.25759 | 0.29655 |
| *C*<sub>*AR*</sub> | −0.62152 | −0.69223 |
| *D*<sub>*I*</sub> | 0.00286 | 0.00137 |
| *C*<sub>*AI*</sub> | −0.06202 | −0.04676 |
| *C*<sub>*IR*</sub> | 0.04684 | 0.03980 |

**Table S4:**
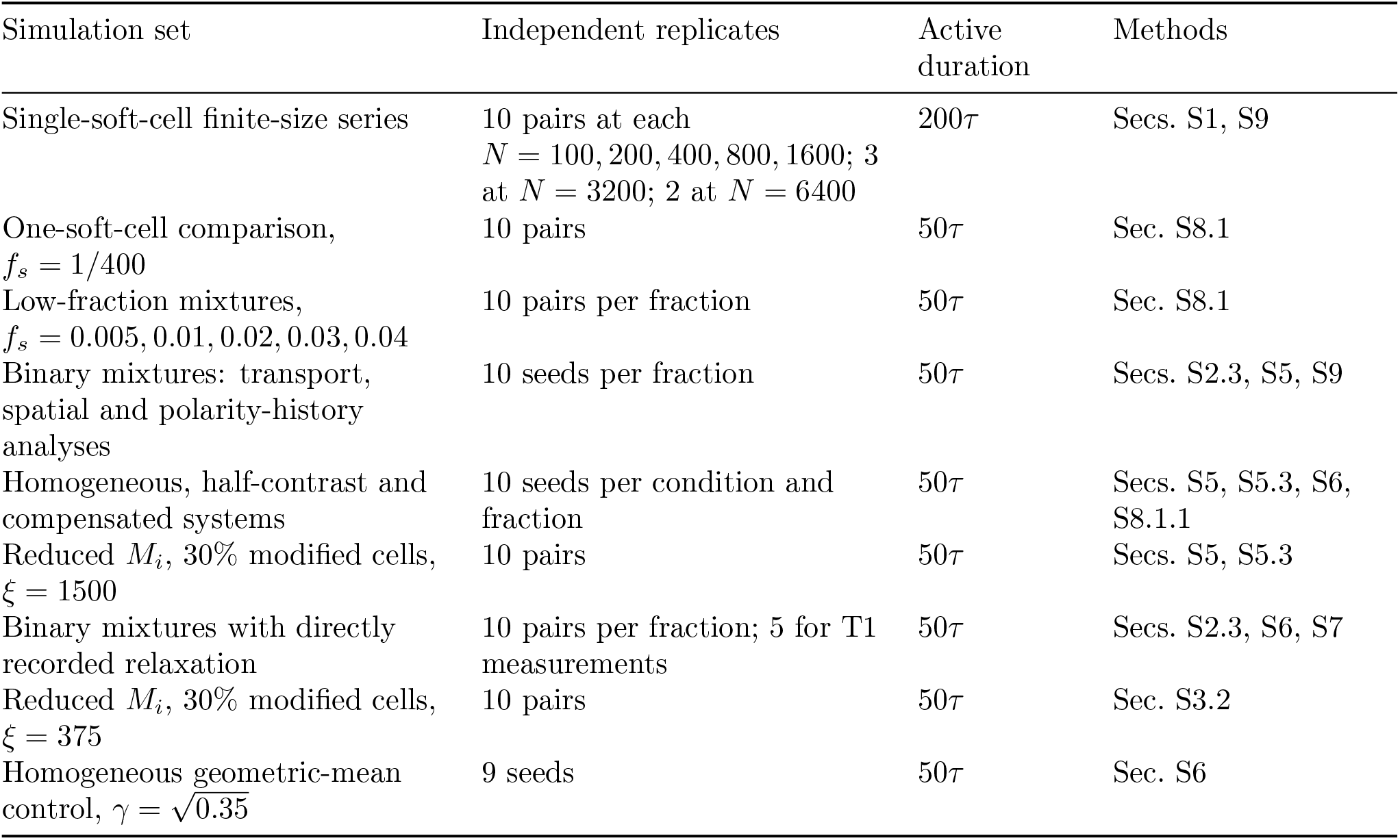
Summary of production simulations. Except for the finite-size series, all systems contain *N* = 400 cells. The six-fraction series uses *f*_*s*_ = 0.05, 0.10, 0.15, 0.20, 0.30 and 0.50. Counts refer to independent seeds or matched seed pairs. Active durations exclude passive preparation.

| Simulation set | Independent replicates | Active duration | Methods |
|---|---|---|---|
| Single-soft-cell finite-size series | 10 pairs at each *N* = 100, 200, 400, 800, 1600; 3 at *N* = 3200; 2 at *N* = 6400 | 200*τ* | Secs. S1, S9 |
| One-soft-cell comparison, *f<sub>s</sub>* = 1/400 | 10 pairs | 50*τ* | Sec. S8.1 |
| Low-fraction mixtures, *f<sub>s</sub>* = 0.005, 0.01, 0.02, 0.03, 0.04 | 10 pairs per fraction | 50*τ* | Sec. S8.1 |
| Binary mixtures: transport, spatial and polarity-history analyses | 10 seeds per fraction | 50*τ* | Secs. S2.3, S5, S9 |
| Homogeneous, half-contrast and compensated systems | 10 seeds per condition and fraction | 50*τ* | Secs. S5, S5.3, S6, S8.1.1 |
| Reduced *M<sub>i</sub>*, 30% modified cells, *ξ* = 1500 | 10 pairs | 50*τ* | Secs. S5, S5.3 |
| Binary mixtures with directly recorded relaxation | 10 pairs per fraction; 5 for T1 measurements | 50*τ* | Secs. S2.3, S6, S7 |
| Reduced *M<sub>i</sub>*, 30% modified cells, *ξ* = 375 | 10 pairs | 50*τ* | Sec. S3.2 |
| Homogeneous geometric-mean control, *γ* = $\sqrt{0.35}$ | 9 seeds | 50*τ* | Sec. S6 |

**Table S5:**
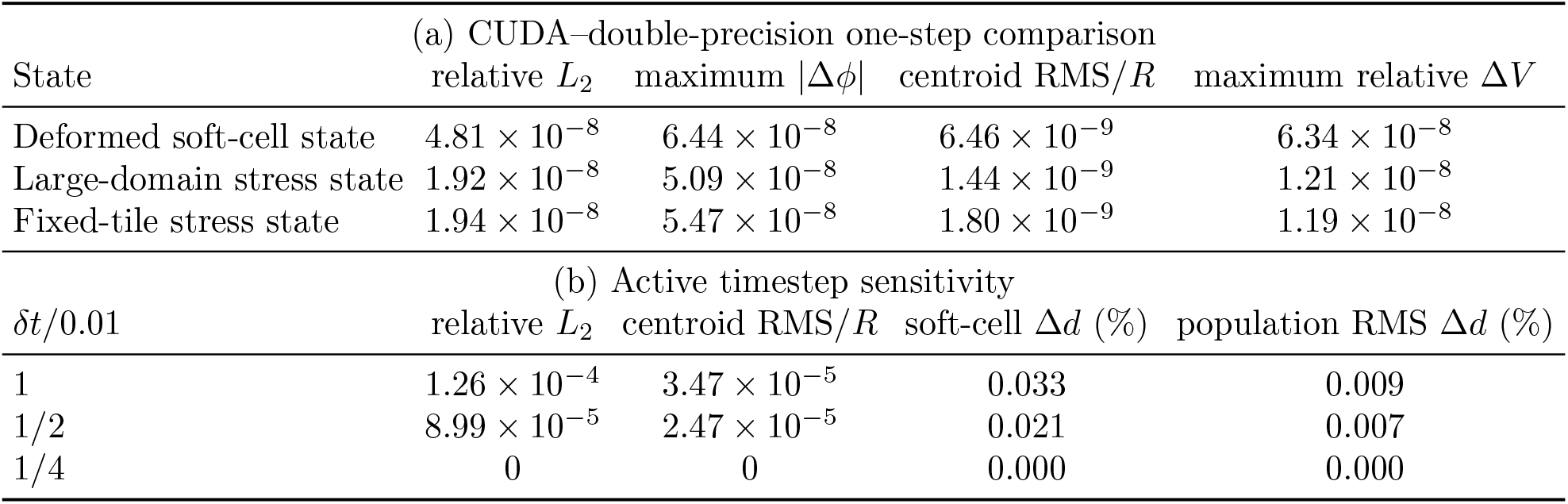
Numerical validation of the mixed-precision CUDA solver. Part (a) compares one CUDA update at *δt* = 0.01 with an independent full-domain double-precision evaluation from the identical checkpoint. Part (b) measures timestep sensitivity over 0.1*τ* with fixed polarity, using *δt* = 0.0025 as the reference. The centroid difference is normalized by the cell radius *R*. The two displacement columns give relative differences from the *δt* = 0.0025 values.

**Figure S1:**
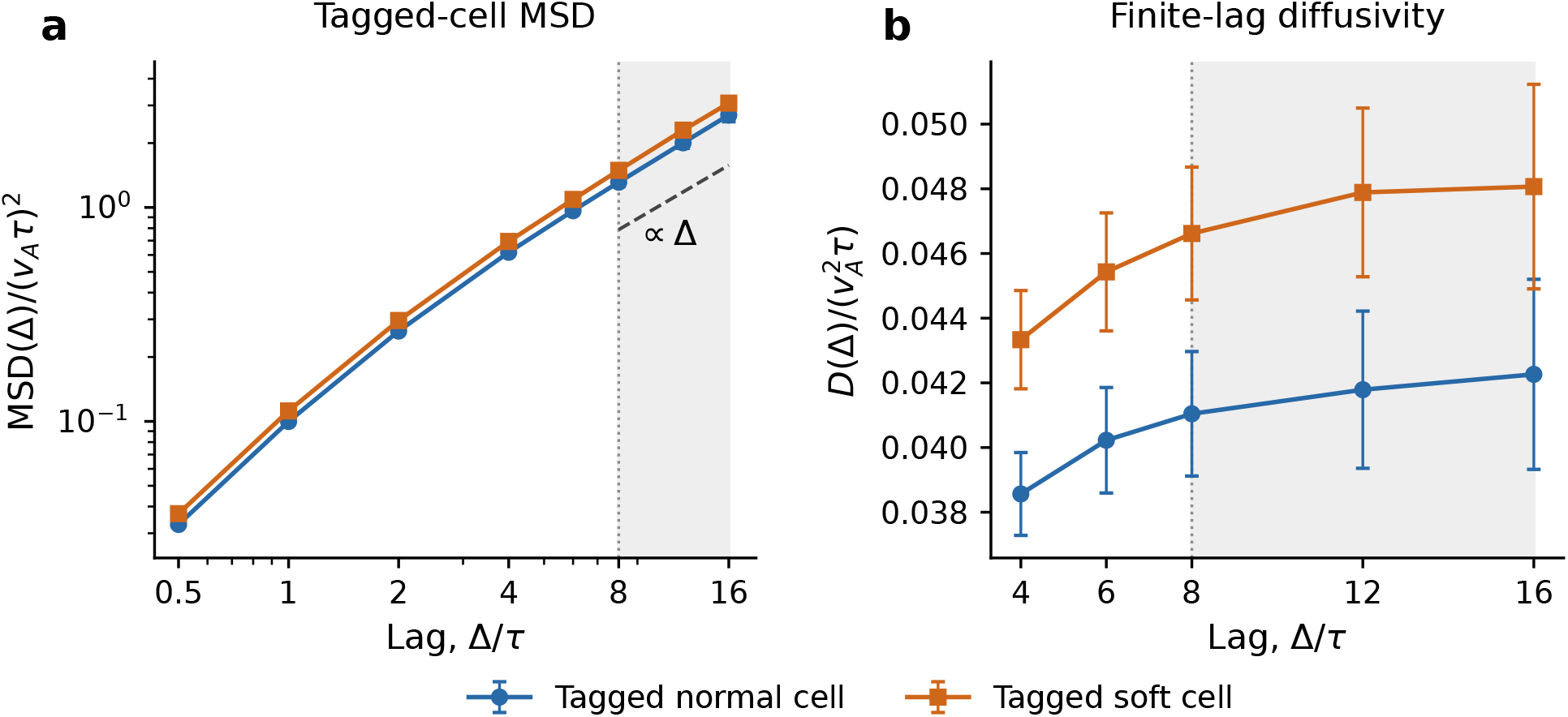
Tagged-cell mean-square displacement and lag dependence of diffusivity. (a) Mean-square displacement of the tagged normal cell in all-normal references and the tagged soft cell in single-soft-cell systems. The dashed line is a unit-slope guide. (b) Finite-lag diffusivity calculated from the same trajectories using Eq. S1. Shading marks the 8*τ* –16*τ* interval. Points show arithmetic means over 50 simulations per condition: 10 at each of *N* = 100, 200, 400, 800 and 1600. Each simulation receives equal weight. Bars give pointwise 95% Student-*t* confidence intervals. Slopes quoted in the text are calculated from the pooled MSD at 8*τ* and 16*τ*. Each lag uses overlapping displacement windows within the 200*τ* active record.

**Figure S2:**
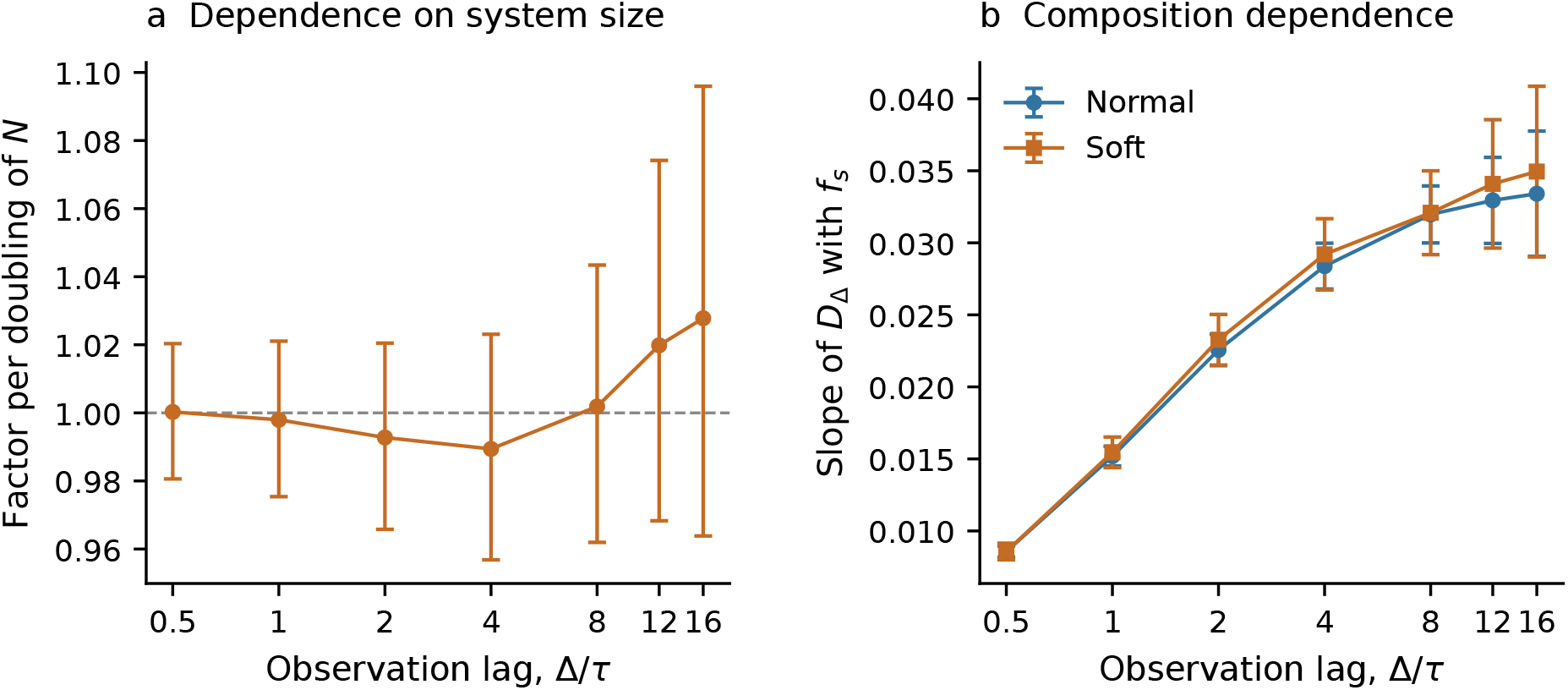
Dependence of transport estimates on observation lag. (a) Multiplicative change in the tagged-cell soft/control transport ratio per doubling of *N*, fitted by regressing the log ratio on log_2_(*N/*100) over *N* = 100–1600. The dashed line marks unity. (b) Soft- and normal-cell slopes of *D*_Δ_ with *f*_*s*_, fitted separately within each seed across the six fractions. Points show means across 10 seeds. Bars give pointwise 95% confidence intervals. Each lag uses overlapping displacement windows from the same trajectories. The pooled tagged-cell ratio is shown in Fig. **??**c.

**Figure S3:**
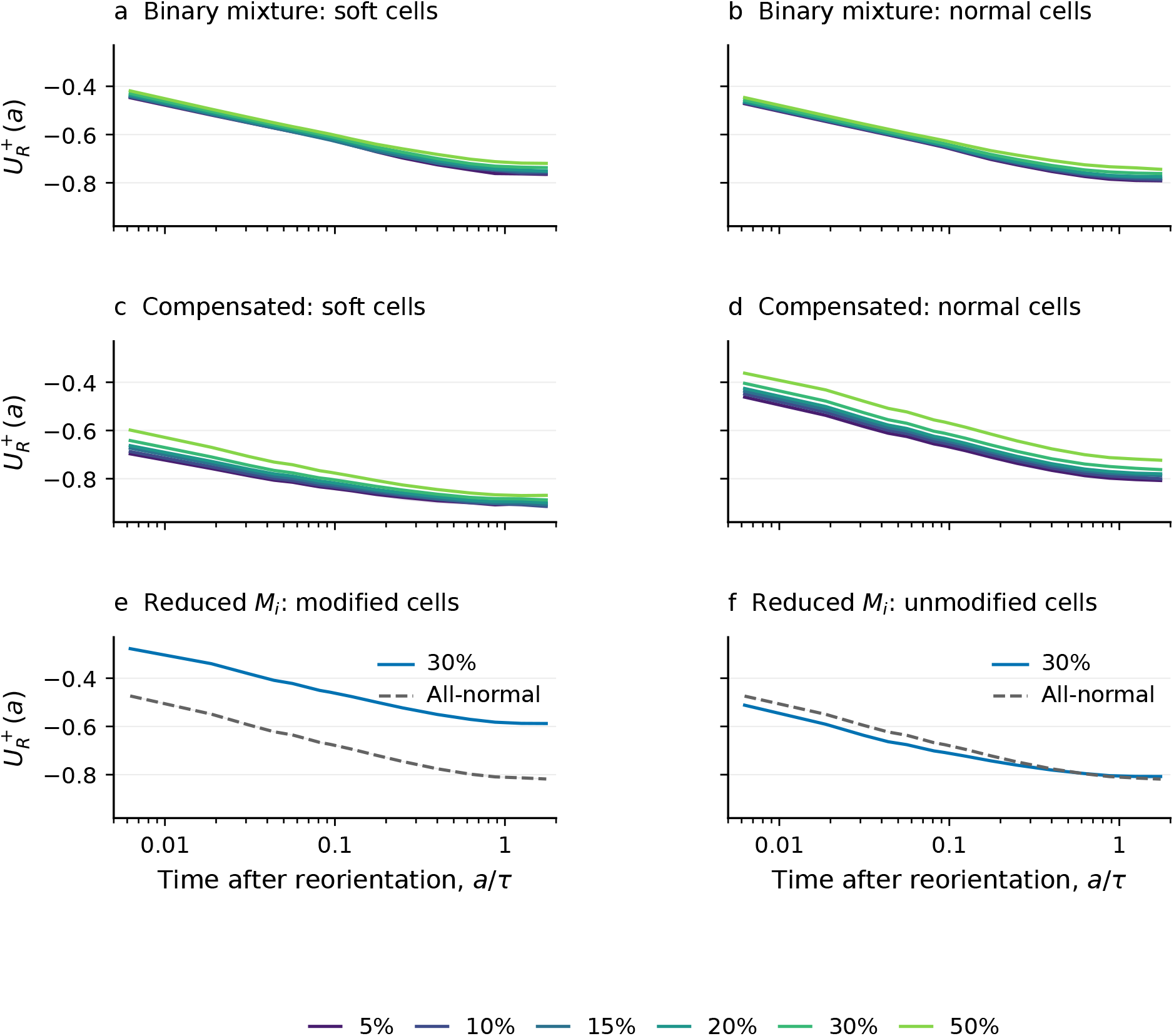
Relaxation along the new propulsion direction. 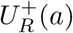 is the mean relaxation-velocity projection onto the cell’s new polarity, divided by *v*_*A*_. Left and right columns show soft and normal cells in the binary and compensated mixtures (a–d), and modified and unmodified cells in the control with reduced *M*_*i*_ (e,f). Colors show all six fractions for the binary and compensated mixtures. The control with reduced *M*_*i*_ uses 30% modified cells with dashed curves for the corresponding labels in the all-normal reference. The time axis is logarithmic. Shading is a pointwise 95% interval across 10 seeds. At each time, we average over cells that have not reoriented again.

**Figure S4:**
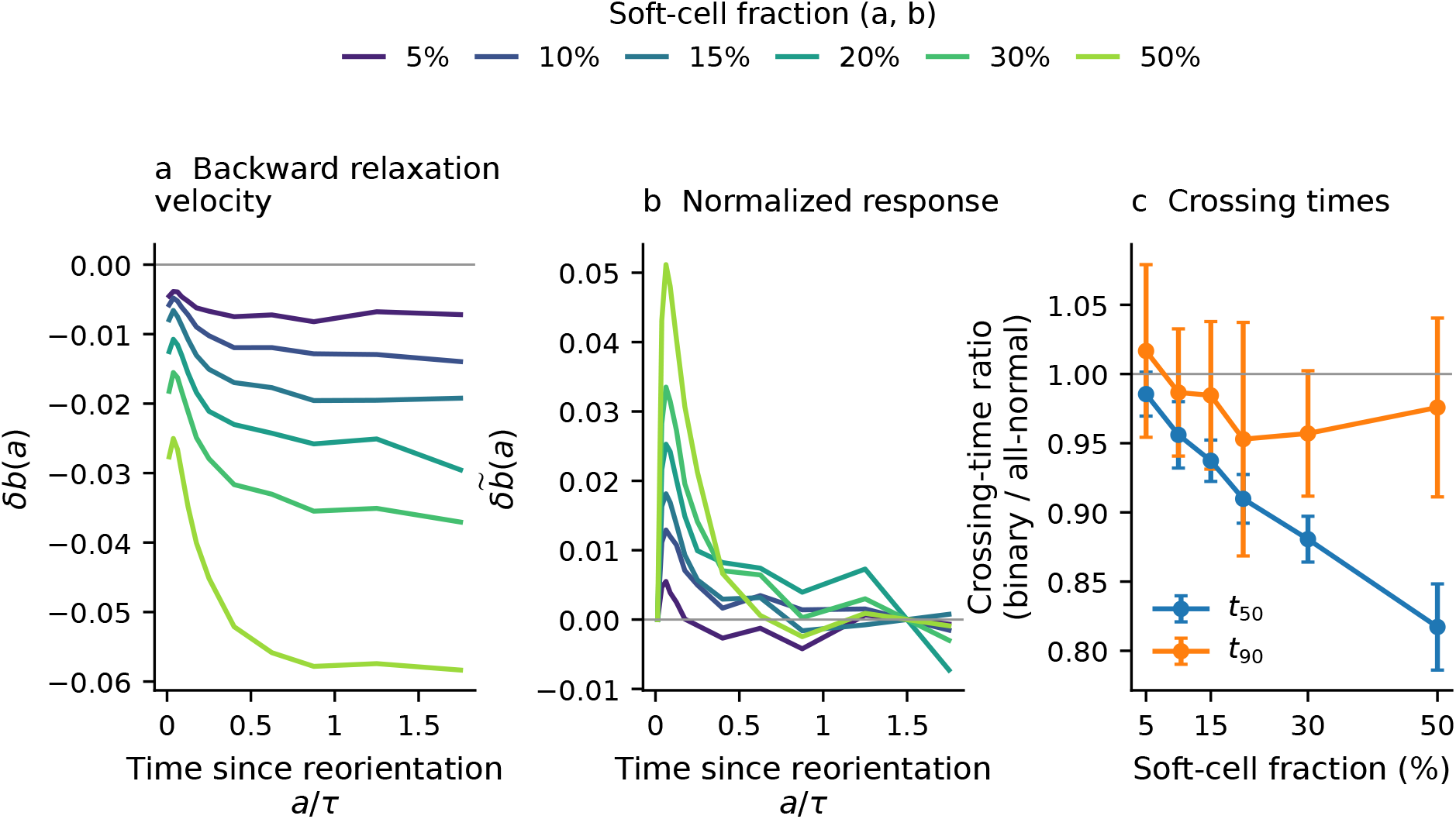
Relaxation after reorientation in normal cells. Binary mixtures across the fraction series relative to matched all-normal controls with 10 paired seeds at each fraction. Left: paired change in backward relaxation velocity at the same time after reorientation. Middle: difference between responses after rescaling each curve to zero in its first time bin and one at late times. Right: binary/control ratios of the 50% and 90% crossing times. Points are paired-seed means and bars are pointwise 95% Student intervals. The late level uses ages 1–2*τ*. The 50% crossing occurs earlier at *f*_*s*_≥ 0.10. All pointwise intervals for the 90% crossing include unity. The observables and averaging are defined in Secs. S2.4 and S7.

**Figure S5:**
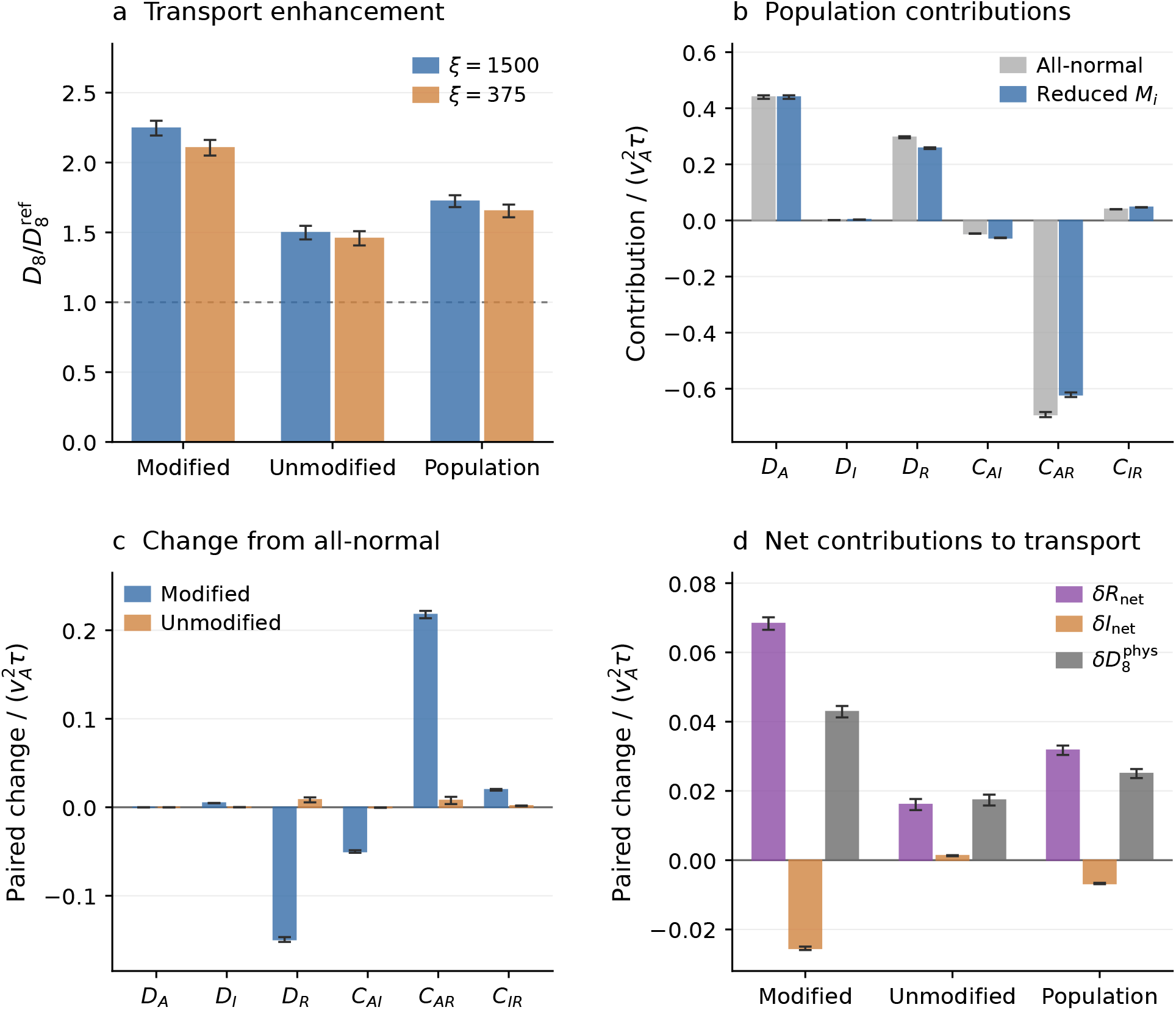
Transport and displacement balance with reduced *M*_*i*_ at fourfold higher advection mobility. (a) Ratios of centroid *D*_8_ between the control with reduced *M*_*i*_ and its all-normal reference for modified cells, unmodified cells and the population, using the same 10 seeds at *ξ* = 1500 and 375. Recorded polarity histories match across both values of *ξ* and both conditions (Sec. S3.2). Bars and error bars show geometric means and pointwise 95% Student intervals. (b) All six displacement contributions for the population at *ξ* = 375, calculated from the recorded velocities. (c) Paired differences from the all-normal reference in each contribution for the modified and unmodified classes. (d) Paired changes in *R*_net_ = *D*_*R*_ + *C*_*AR*_ and *I*_net_ = *D*_*I*_ + *C*_*AI*_ + *C*_*IR*_. Their sum equals the transport change calculated from the recorded velocity contributions because *δD*_*A*_ = 0. Panels b–d show arithmetic means and pointwise 95% intervals across the 10 pairs. Transport contributions are in units of 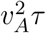. Cells and overlapping windows are averaged within seed. All panels use complete 8*τ* windows within the 50-*τ* active record. The numerical comparison with centroid transport is given in Sec. S3.2.

**Figure S6:**
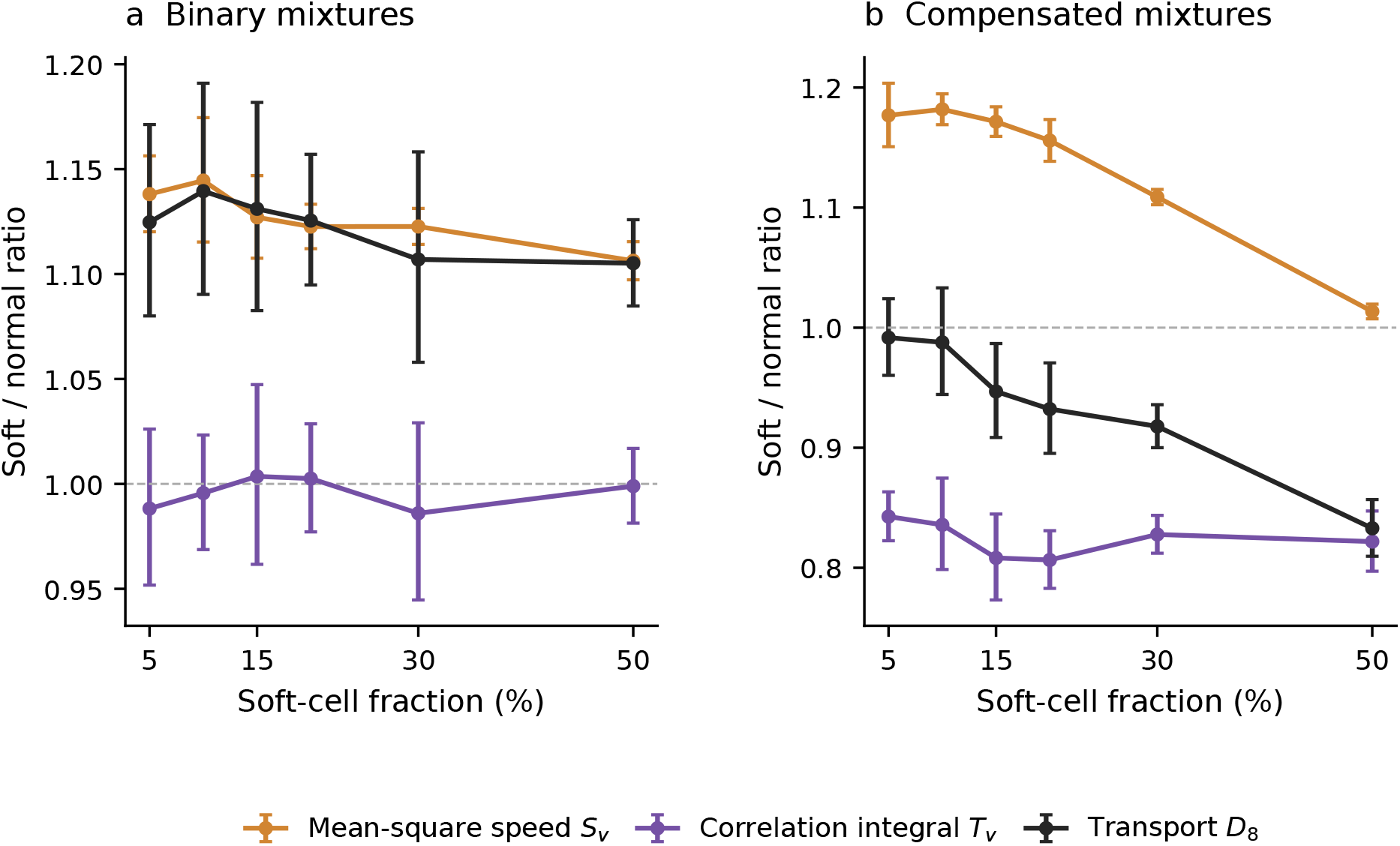
Speed and temporal correlations in binary and compensated mixtures. Soft-to-normal ratios of mean-square centroid speed *S*_*v*_, finite-lag normalized velocity-correlation integral *T*_*v*_ = 2*D*_8_*/S*_*v*_, and *D*_8_ in binary and compensated mixtures. The transport ratio is the product of the other two. Ratios are constructed from class means within each seed. Displayed geometric means and pointwise 95% intervals are calculated across 10 seeds.

**Figure S7:**
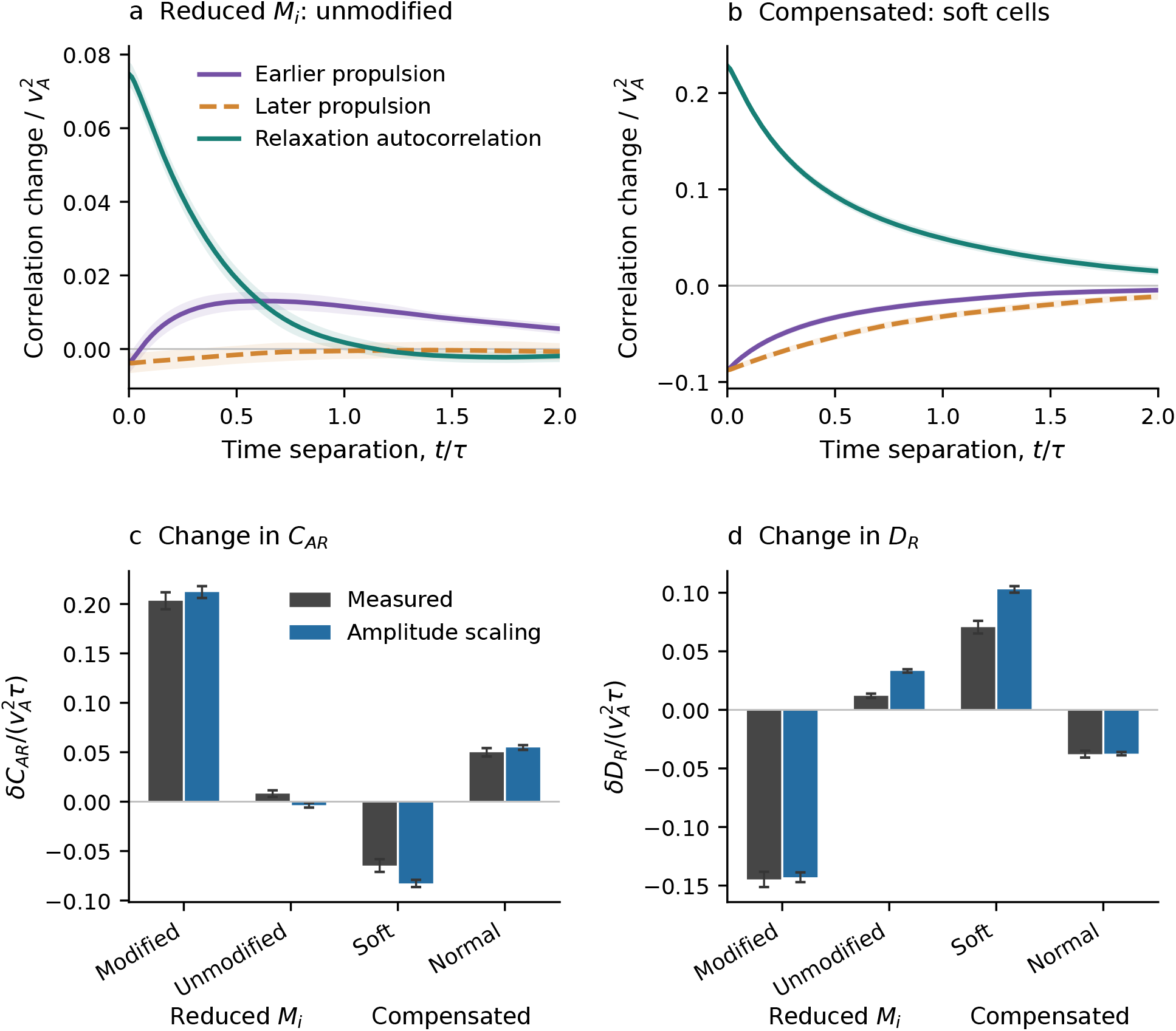
Measured and amplitude-rescaled correlation contributions. All comparisons use *f*_*s*_ = 0.30 and matched all-normal references. (a,b) Changes in *K*_*AR*_(*t*) (earlier propulsion), *K*_*RA*_(*t*) (later propulsion), and *K*_*RR*_(*t*), divided by 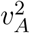, for unmodified cells in the control with reduced *M*_*i*_ (a) and soft cells in the compensated mixture (b). The two ordered cross correlations are not assumed equal. (c,d) Measured displacement-term changes and predictions obtained by scaling each reference correlation by its equal-time amplitude ratio, for both classes in the control with reduced *M*_*i*_ and the compensated mixtures. Contributions are in units of 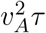. Panels a and b use correlations of velocities averaged over the recorded time intervals. The finite-window calculation uses the weights in Sec. S5.1. Curves and bars show means and pointwise 95% intervals across 10 seeds.

**Figure S8:**
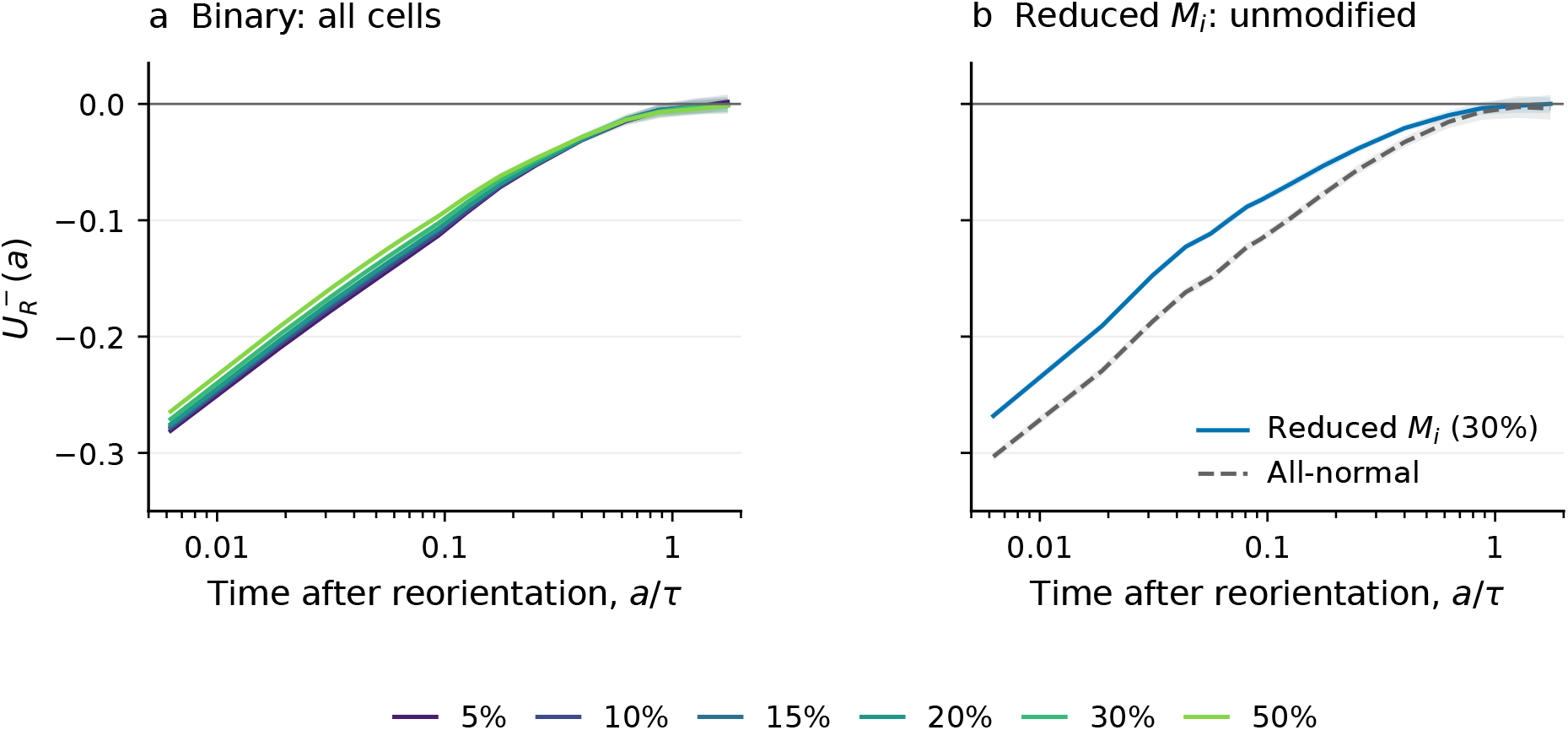
Relaxation projected onto the preceding propulsion direction after reorientation. 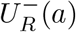 uses the direction immediately before the cell’s latest recorded reorientation. The early 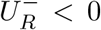 approaches zero with increasing *a*. Each point includes only cells that have not reoriented again. Time is plotted on a logarithmic axis. Panel (a) shows all six fractions for the binary population. Panel (b) compares unmodified cells in the control with reduced *M*_*i*_ with the corresponding all-normal labels at 30%. Shading gives pointwise 95% intervals over 10 seeds. These curves resolve the immediately preceding direction, whereas 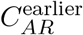 retains correlations with every earlier polarity interval that intersects the displacement window.

**Figure S9:**
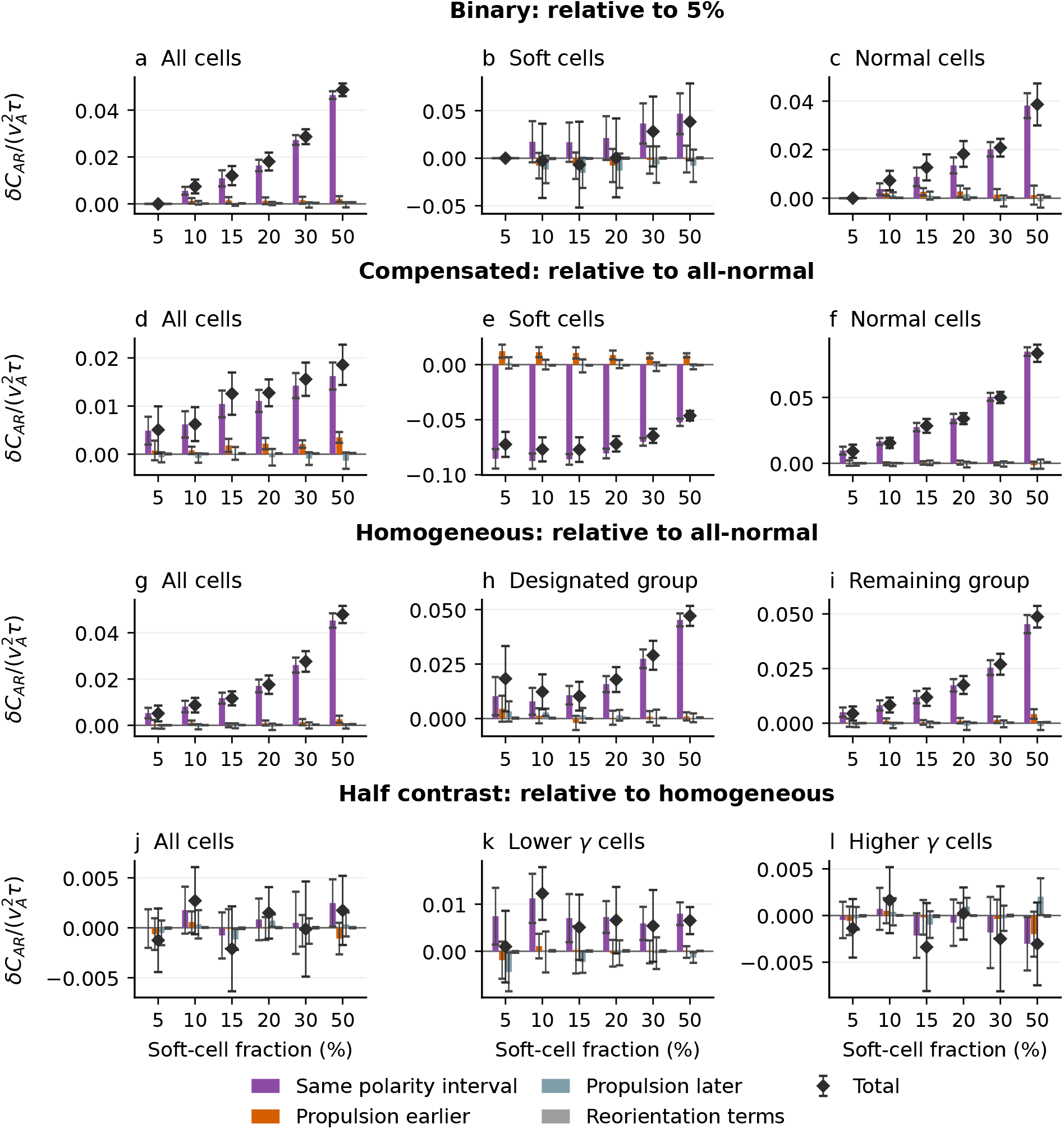
Contributions to the change in *C*_*AR*_ by polarity interval. Columns show all cells, designated cells, and remaining cells. Each row states its reference: 5% within the binary mixture, all-normal for compensated and homogeneous systems, and matched homogeneous mean *γ* for half contrast. The bars separate four contributions to the change in *C*_*AR*_: propulsion and relaxation within the same interval of constant polarity; propulsion before relaxation in different intervals; propulsion after relaxation in different intervals; and terms involving a recording interval containing a reorientation. Diamonds are their sum, the complete change in *C*_*AR*_ at 8*τ*. The initial polarity interval and all age bands enter the sums. Error bars are pointwise 95% intervals over 10 paired within-seed differences. Units of 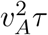.

**Figure S10:**
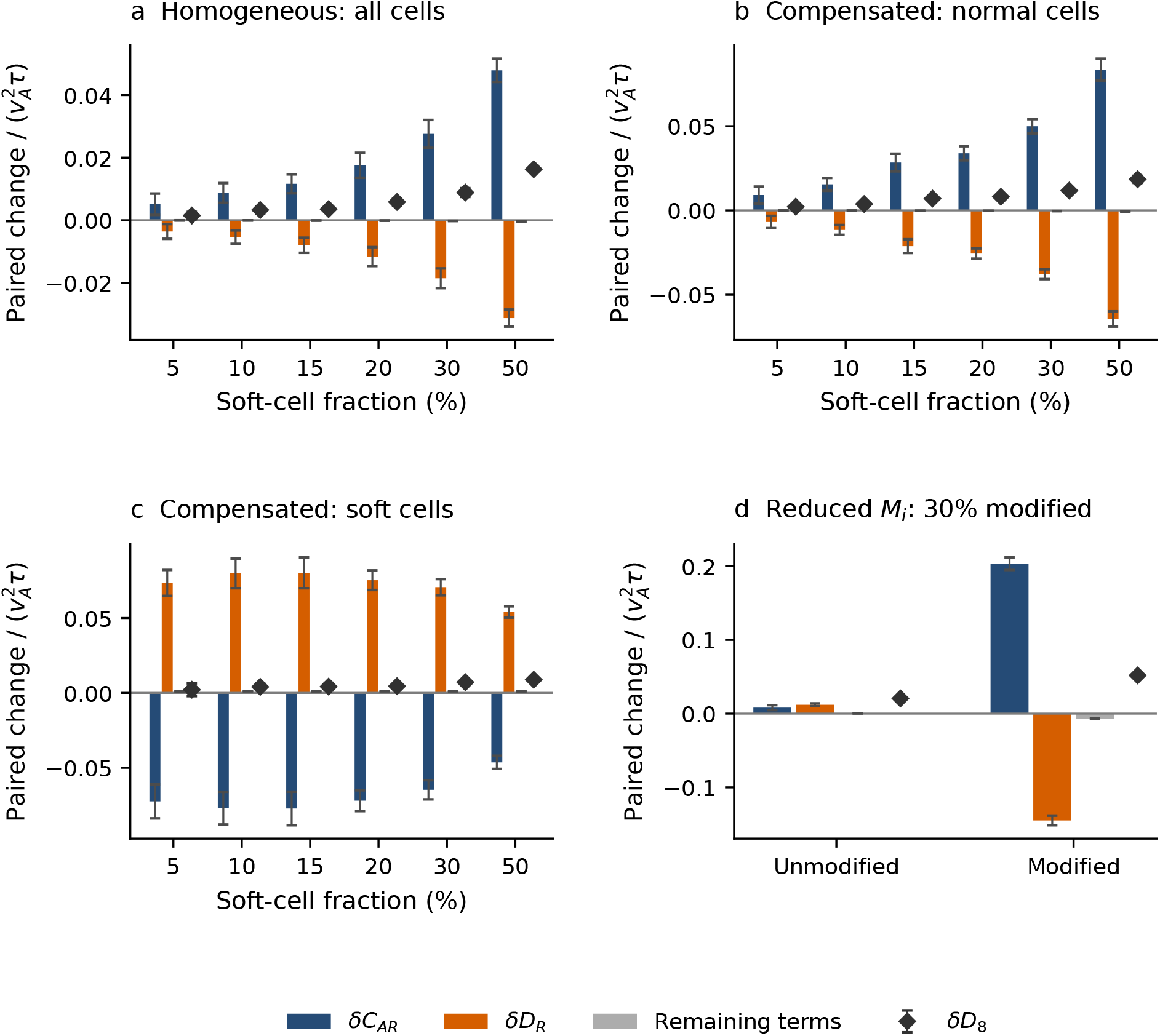
Displacement contributions in the control simulations. Differences from matched all-normal references for (a) homogeneous populations at 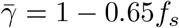, (b,c) normal and soft cells in compensated mixtures and (d) unmodified and modified cells in the control with *M*_*i*_ reduced in 30% of cells. Bars show *δC*_*AR*_, *δD*_*R*_ and *δ*(*D*_*A*_ + *D*_*I*_ + *C*_*AI*_ + *C*_*IR*_). Their sum is *δD*_8_ (diamonds). Relaxation is estimated from centroid displacement minus the integrated advection contributions. Each comparison uses Δ = 8*τ*, 10 seed pairs, and means with pointwise 95% Student intervals. Vertical scales differ between panels.

**Figure S11:**
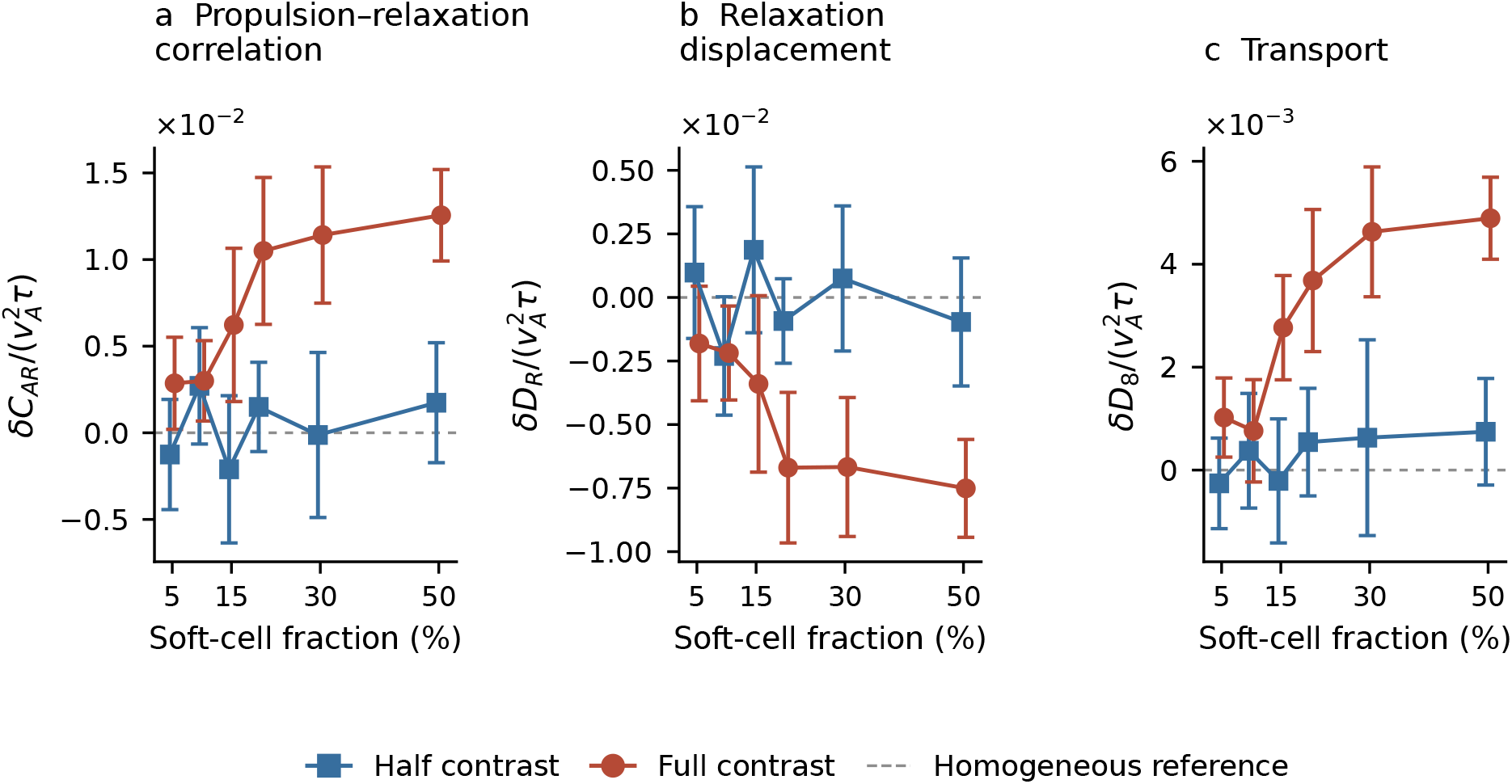
Binary-mixture transport at fixed mean interfacial coefficient. Population differences from the homogeneous system at 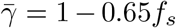, for half and full interfacial contrast. The dashed zero line denotes the homogeneous reference at each fraction. (a) Propulsion–relaxation cross term, *δC*_*AR*_. (b) Relaxation displacement contribution, *δD*_*R*_. (c) Centroid transport, *δD*_8_. Here *δ* denotes mixture minus homogeneous. All three panels use the same 10 matched seeds and common 8*τ* displacement windows. Symbols show seed means and bars show pointwise 95% Student intervals. Horizontal offsets separate the two mixtures visually. Total relaxation is estimated from centroid displacement and recorded advection with sampling and numerical errors assessed in Sec. S2.3. These recordings do not resolve its interfacial and overlap components.

**Figure S12:**
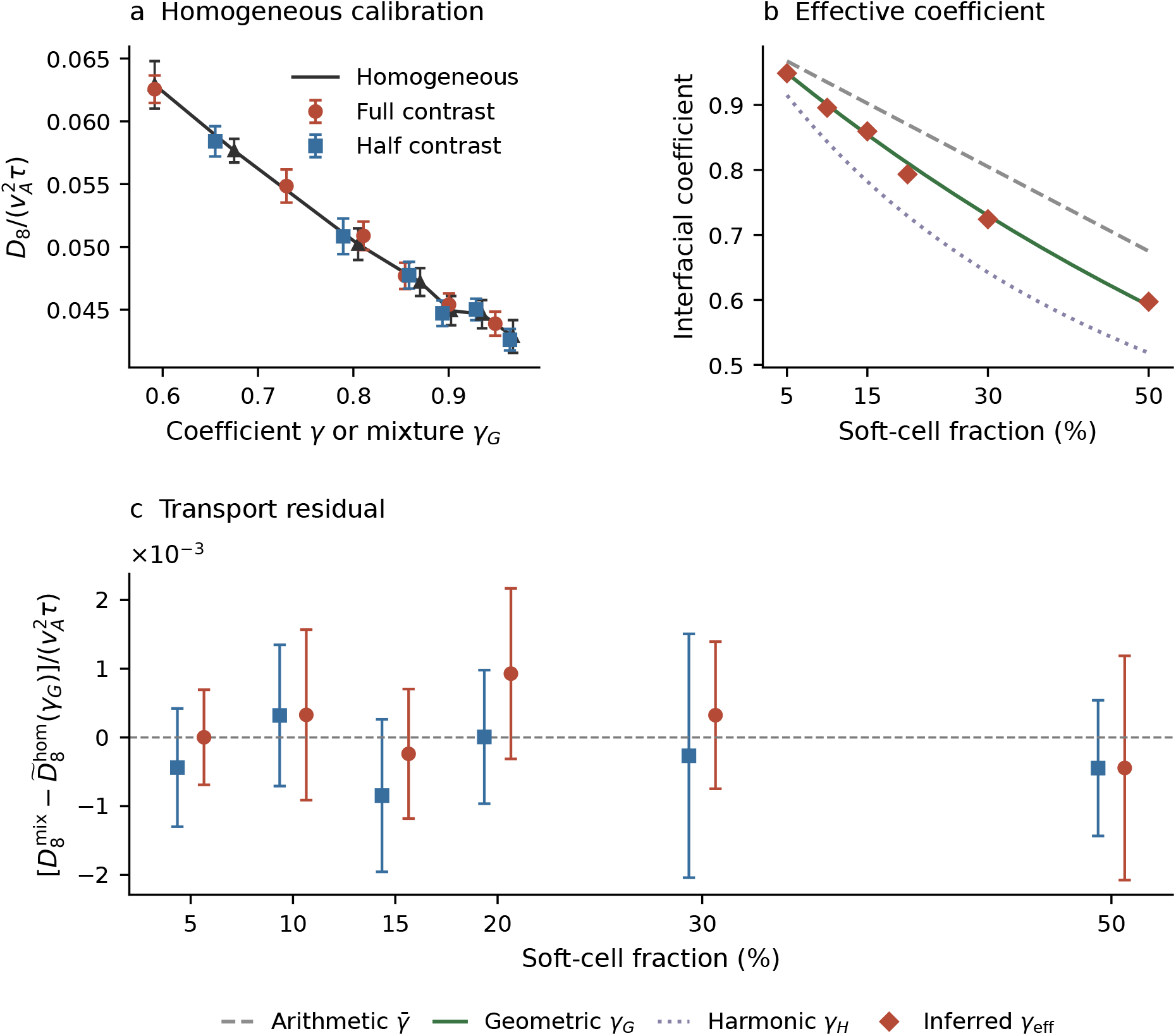
Geometric-mean calibration of population transport. (a) Homogeneous population-mean *D*_8_ versus *γ* with mixture transports plotted at their geometric-mean coefficients *γ*_*G*_. The calibration includes the directly simulated homogeneous control at 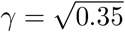. The line interpolates the homogeneous means linearly. (b) Full-contrast effective coefficients inferred by inverting this calibration, compared with arithmetic, geometric and harmonic means. All six inverse estimates lie within the measured calibration range. (c) Mixture-minus-homogeneous transport residuals at *γ*_*G*_, calculated within seed. The 50% full-contrast residual uses the direct control. The other residuals use interpolation. Horizontal offsets distinguish contrast levels. Points and bars show arithmetic means and pointwise 95% Student-*t* intervals. The direct control and the paired 50% residuals use 9 seeds. The other estimates use ten. Each recording averages all 400 cells over complete 8*τ* displacement windows common to the compared recordings within 0.01–50*τ* of activity. The calibration and uncertainty calculation are described in Sec. S6.

**Figure S13:**
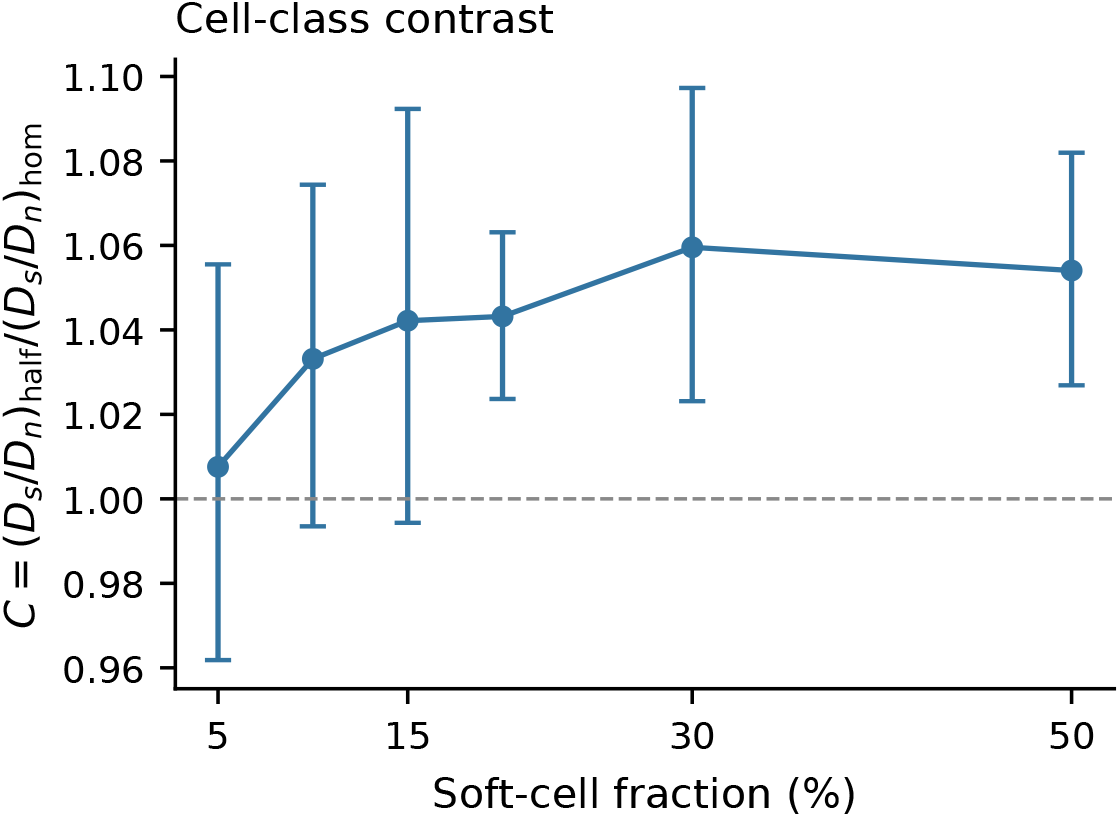
Cell-class transport contrast in half-contrast mixtures. The ratio *C* compares the lower-/higher-*γ* transport ratio with the ratio for the corresponding labels in the homogeneous reference at the same 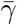 (Eq. S26). The mixture coefficients are given by Eq. S39. Points and bars are geometric means and pointwise 95% confidence intervals across 10 paired seeds, at Δ = 8*τ*. The dashed line marks unity.

**Figure S14:**
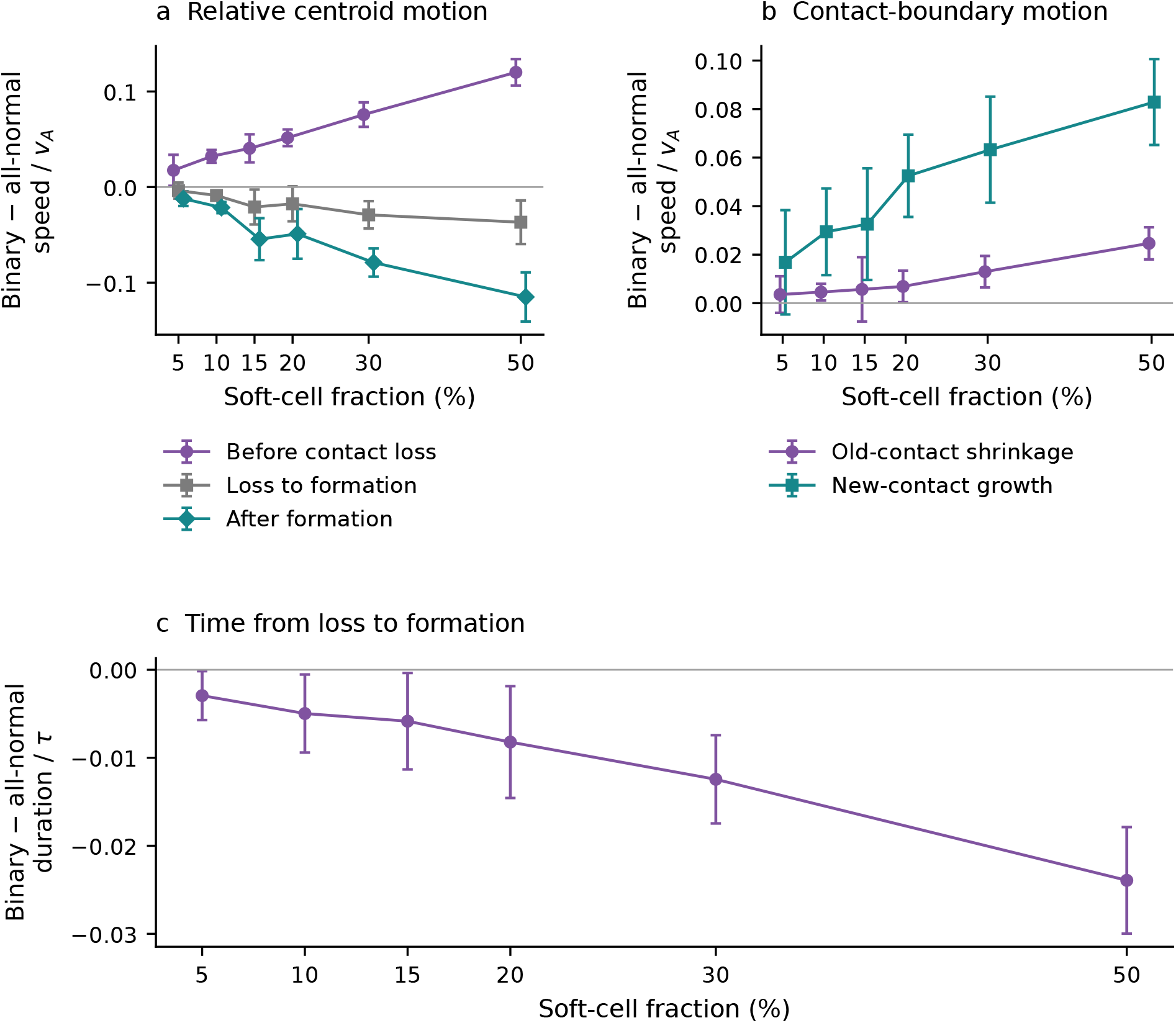
Centroid motion, boundary motion and duration during T1 exchanges. Binary-minus-all-normal changes across soft-cell fractions. (a) Relative centroid speed, defined by *u*_*q*_ = d*q/*d(*t/τ*) for *q* = (*d*_old_ −*d*_new_)*/*(*v*_*A*_*τ*), in the last 0.1*τ* with the old contact intact, the interval from old-contact loss to new-contact formation, and the first 0.1*τ* with the new contact intact. (b) Mean shortening speed of the old contact and growth speed of the new contact in the corresponding 0.1*τ* windows (Eq. S31). (c) Duration *T* = *t*_*n*_ −*t*_*o*_, using the last old-contact and first new-contact frames. The frame interval is 0.01*τ*. These three measurements use the same accepted four-cell exchanges. Events receive equal weight within each seed. Symbols and pointwise 95% Student intervals use five paired seeds at each fraction. Seed means are the independent observations. Events in the two systems need not be the same. The same all-normal simulations provide the reference at each soft-cell fraction, so comparisons at different fractions are correlated.

**Figure S15:**
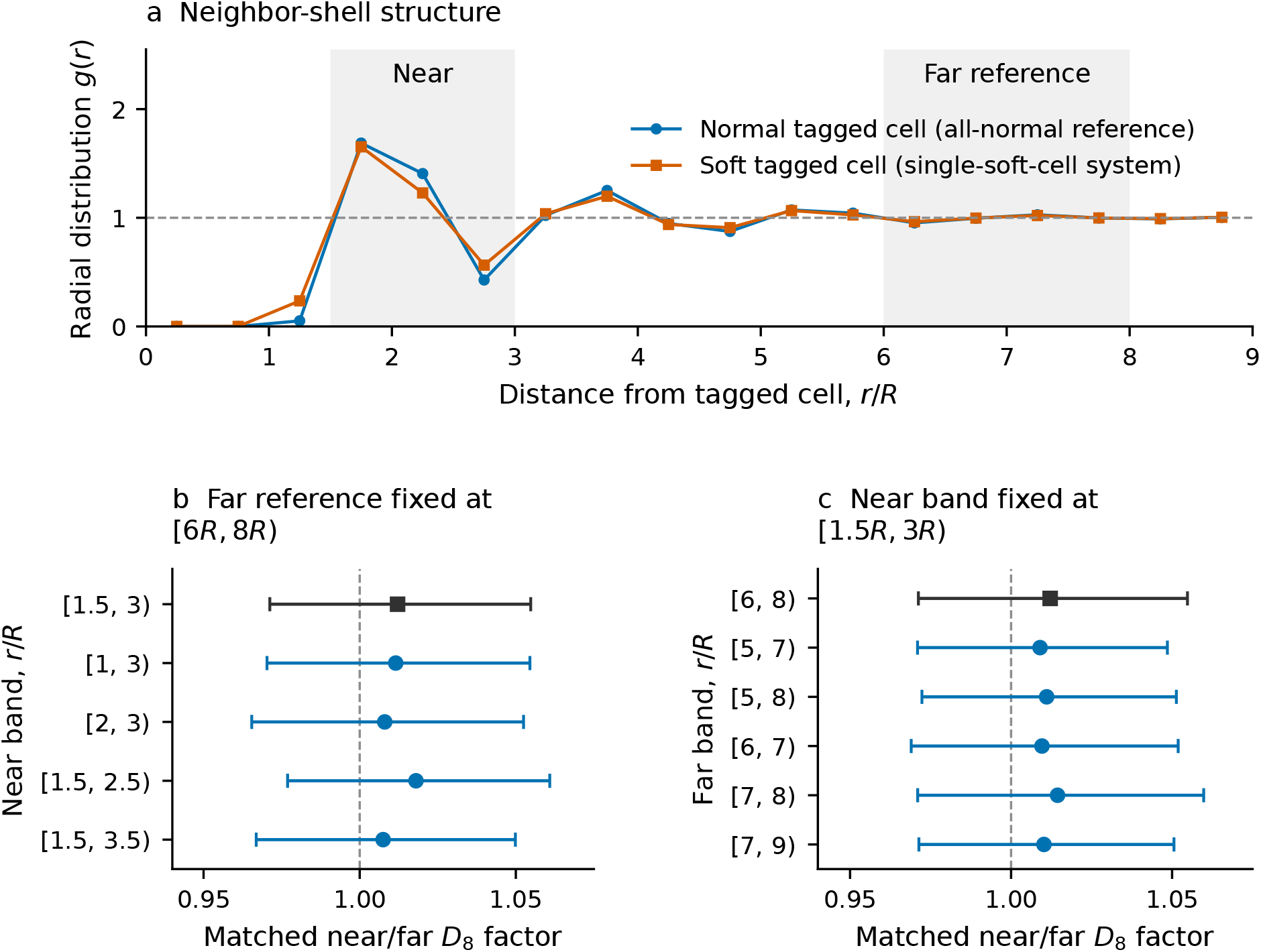
Structural basis and sensitivity of the distance bands. (a) Radial distribution of normal-cell centroids around the tagged cell in the single-soft-cell and all-normal systems. Gray regions mark the original near and far bands. (b,c) Matched near/far *D*_8_ factors when one band is varied and the other retained. Squares show the original comparison. Curves in (a) show arithmetic means, whereas factors in (b,c) show geometric means. Shading and bars give pointwise 95% Student-*t* intervals across 10 replicate groups, each giving equal weight to *N* = 400, 800 and 1600. Each size contributes 10 paired 200*τ* simulations with Δ = 8*τ*. All tested outer reference radii remain below one quarter of the box side.

